# Redirecting the pioneering function of FOXA1 with covalent small molecules

**DOI:** 10.1101/2024.03.21.586158

**Authors:** Sang Joon Won, Yuxiang Zhang, Christopher J. Reinhardt, Nicole S. MacRae, Kristen E. DeMeester, Evert Njomen, Lauren M. Hargis, Jarrett R. Remsberg, Bruno Melillo, Benjamin F. Cravatt, Michael A. Erb

**Affiliations:** Department of Chemistry, The Scripps Research Institute, La Jolla, CA 92037, USA

**Keywords:** FOXA1, covalent chemistry, proteomics, activity-based protein profiling, cysteine, chromatin, pioneer transcription factor, DNA-binding motif, ATAC-seq, ChIP-seq

## Abstract

Pioneer transcription factors (TFs) exhibit a specialized ability to bind to and open closed chromatin, facilitating engagement by other regulatory factors involved in gene activation or repression. Chemical probes are lacking for pioneer TFs, which has hindered their mechanistic investigation in cells. Here, we report the chemical proteomic discovery of electrophilic small molecules that stereoselectively and site-specifically bind the pioneer TF, FOXA1, at a cysteine (C258) within the forkhead DNA-binding domain. We show that these covalent ligands react with FOXA1 in a DNA-dependent manner and rapidly remodel its pioneer activity in prostate cancer cells reflected in redistribution of FOXA1 binding across the genome and directionally correlated changes in chromatin accessibility. Motif analysis supports a mechanism where the covalent ligands relax the canonical DNA binding preference of FOXA1 by strengthening interactions with suboptimal ancillary sequences in predicted proximity to C258. Our findings reveal a striking plasticity underpinning the pioneering function of FOXA1 that can be controlled by small molecules.

## Introduction

Lineage transcription is controlled by cell-type-specific enhancer elements that regulate the activity of distal gene promoters^1^. DNA-binding transcription factors (TF) recognize densely positioned cognate binding sequences within these *cis*-regulatory elements and then recruit transcriptional co-regulators to activate or repress gene expression. Most TFs cannot associate with condensed (e.g., nucleosome-bound) DNA and therefore rely on pioneer TFs to first open such heterochromatin sites in a locus-specific manner^2^. Mechanistically, pioneer TFs are distinguished by the ability to (i) bind closed chromatin and (ii) elicit the subsequent decompaction of that chromatin region. Due to their first-order control of enhancer selection and function, pioneer TFs are critical to gene regulatory changes that underly organismal development and tumorigeneses^3,4^.

Studies of Forkhead box protein A1 (FOXA1) have played a prominent role in defining the properties of pioneer TFs, beginning with the foundational discoveries that it can bind nucleosomal DNA and initiate local chromatin decompaction *in vitro*^5,6^. FOXA1 is critical for liver development^7-9^, and has been identified as an essential gene supporting the growth of breast and prostate cancer cells^10-12^. FOXA1 binds to DNA through its forkhead domain (FKHD), which consists of a helix-turn-helix motif flanked by Wing1 and Wing2 loops that assist in high-affinity DNA binding^13^. Interestingly, in addition to the lineage dependency relationship between wild-type FOXA1 and breast and prostate cancers, mutations in the FKHD domain of this TF are also enriched in these same cancer types, particularly within the Wing2 region^14-19^, and these cancer-related mutations alter FOXA1 pioneer activity^20-22^.

Despite the clear importance of pioneer factors in normal development and tumorigenesis, the mechanisms of action of these key regulatory proteins remain difficult to study in living systems. Investigations of FOXA1 and other pioneer factors have largely relied on correlating *in vitro* biochemical activities with cellular phenotypes derived from genetic disruption or heterologous expression systems^23^. However, inferring the direct biochemical functions of TFs, especially those involved in the dynamic regulation of chromatin structure, from cell models of prolonged genetic perturbation or protein overexpression can be problematic^24,25^. Additionally, the lineage relationship between cancer cells and pioneer TFs like FOXA1 often results in strong growth dependency on these proteins, as revealed in the Cancer Dependency Map^26^, that further hinders their study by constitutive genetic disruption methods. Our understanding of the direct mechanisms by which pioneer TFs function in cells accordingly remain incomplete^23^.

We and others have shown that chemical probes provide a versatile means to acutely perturb and study transcriptional regulatory proteins in cells^25,27-29^. Historically, the discovery of chemical probes for TFs has been challenged by the general lack of structurally ordered small molecule-binding pockets in this class of proteins (nuclear hormone receptors being a notable exception^30^) and their complex modes of cellular regulation, which can be difficult to preserve with purified preparations of protein^29^. We and others have attempted to address these challenges by developing chemical proteomic methods, such as activity-based protein profiling (ABPP), for the global mapping of small molecule-protein interactions directly in native biological systems^31,32^. Integrating ABPP with, for instance, fragment-based^33-35^ or stereochemically defined sets (‘stereoprobes’)^36-39^ of cysteine-directed electrophilic compounds has identified covalent ligands for a wide range of proteins, including many transcriptional regulators^31-39^.

Here we report the chemical proteomic discovery and functional characterization of tryptoline acrylamide stereoprobes that stereoselectively and site-specifically react with a cysteine (C258) in FOXA1 in prostate cancer cells. This cysteine is located in the Wing2 region of FOXA1 in predicted close proximity to the DNA-binding interface and surrounded by a variety of somatic mutations enriched in breast and prostate cancer. Studies with recombinant protein revealed that the stereoprobes react with FOXA1_C258 in a DNA-dependent manner. In prostate cancer cells, the stereoprobes rapidly remodel the chromatin binding of endogenously expressed FOXA1, resulting in both losses and gains of hundreds of sites that show corresponding changes in chromatin accessibility. Mechanistic studies indicate that the stereoprobes strengthen FOXA1 binding to suboptimal DNA sequence motifs both *in vitro* and *in situ*, thus revealing how small molecules can reshape the global chromatin interaction landscape of a pioneer TF in cells.

## Results

### Covalent ligands stereoselectively and site-specifically engage cysteine 258 in FOXA1

As part of a broader effort to expand the ligandability of the human proteome, we performed cysteine-directed ABPP in the prostate cancer cell line 22Rv1 treated with a set of four tryptoline acrylamide stereoprobes (20 µM, 3 h) – WX-02-13, WX-02-23, WX-02-33 and WX-02-43^37^ (**Figure 1A**). In these experiments, stereoselectively liganded cysteines were defined as those showing > 67% decrease in iodoacetamide-desthiobiotin (IA-DTB) reactivity from cells treated with one stereoprobe, where this decrease was also two-fold or greater than that observed from cells treated with the corresponding enantiomeric stereoprobe. Based on these criteria, several stereoselectively liganded cysteines were identified (**Figures S1A**), including a robust interaction between FOXA1_C258 and the (1*R*, 3*S*) stereoprobe WX-02-23 (**Figures 1B**, **C**). Another quantified cysteine in FOXA1 (C268) was unchanged by WX-02-23 (**Figure 1B**), supporting that this compound engaged FOXA1 in a site-specific manner. Across more than 18,000 quantified cysteines, WX-02-23 showed substantial reactivity (> 67%) with only a handful of cysteines in addition to FOXA1_C258, and the majority of these cysteines showed similar reactivity with the enantiomer WX-02-43 (**Figures S1A, B** and **Supplementary Dataset S1**).

**Figure 1.**
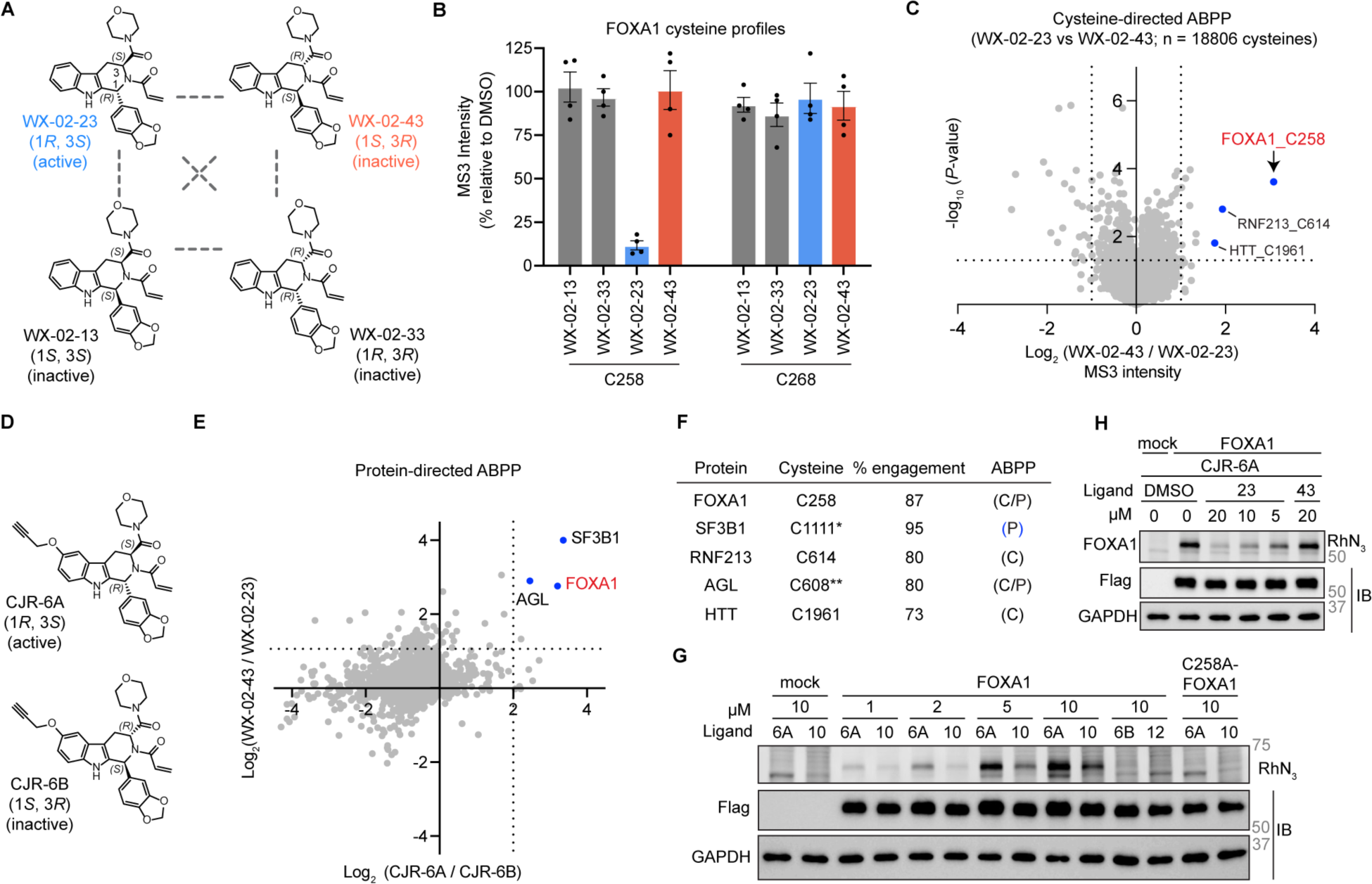
Tryptoline acrylamide WX-02-23 stereoselectively and site-specifically engages cysteine-258 (C258) in FOXA1 in human prostate cancer cells. **A.** Chemical structures of tryptoline acrylamide stereoprobes WX-02-23 (FOXA1_C258 ligand), WX-02-43 (enantiomer of WX-02-23), WX-02-13, and WX-02-33 (diastereomers of WX-02-23). **B.** Cysteine-directed ABPP data for two cysteines (C258, C268) in FOXA1 from 22Rv1 cells treated with tryptoline acrylamide stereoprobes (20 μM, 3 h). N = 4 per group. Mean ± SEM. **C.** Volcano plot showing global stereoselectivity profiles in IA-DTB blockade by WX-02-23 and WX-02-43 in same cysteine-directed ABPP experiments shown in Figure 1B. Outside dashed vertical lines indicate cysteines with substantial (MS3 intensity ratio > 2-fold) stereoselective IA-DTB blockade. Above horizontal dashed lines indicate cysteines with *P*-values < 0.05. Cysteines with blue circle indicate > 66.7 % engagement by WX-02-23. Cysteines were required to originate from proteins with at least two quantified cysteines, and proteins where all cysteines uniformly changed were also removed to ensure only stereoprobe-induced changes in reactivity (rather than potential changes in protein expression) were interpreted. Average values from 2 independent experiments shown. *P*-values were determined by two-tailed Student’s t-test. **D.** Chemical structures of alkyne stereoprobes CJR-6A and CJR-6B. **E.** Protein-directed ABPP data showing proteins stereoselectively enriched by CJR-6A versus CJR-6B (x-axis) and proteins for which enrichment by CJR-6A is stereoselectively competed by WX-02-23 (y-axis). 22Rv1 cells were treated with WX-02-23 or WX-02-43 (20 µM, 3 h), or DMSO, followed by CJR-6A or CJR-6B (5 µM, 1 h), and samples analyzed by protein-directed ABPP as described^37,38^. Right of vertical dashed line indicates proteins showing > 4-fold higher enrichment by CJR-6A vs CJR-6B; above horizontal dashed line indicates proteins showing > 2-fold competitive blockade of CJR-6A enrichment by WX-02-23 vs WX-02-43. Stereoselectively enriched and competed proteins that displayed > 66.7% blockade of enrichment by WX-02-23 are marked with blue circles and UniProt names. N = 4 biological replicates, data are averaged. **F.** Table showing proteins (and the cysteines in these proteins) that were substantially and stereoselectively engaged by WX-02-23 as determined by integrated cysteine-directed (C) and protein-directed (P) ABPP studies. Proteins shown meet the following criteria: Substantial (> 66.7 %) engagement by WX-02-23 that was also > 2-fold higher than WX-02-43. % engagement values are averaged between the 2 chemical proteomic platforms (C and P). *Liganded cysteines assigned previously^37^, **mapped in Ramos cells. **G.** Gel-ABPP data showing stereoselective reactivity of FOXA1_C258 with alkyne stereoprobes CJR-6A (6A) and WX-01-10 (10; see **Figures S1D**). Data are from HEK293T cells recombinantly expressing 2xStrep-Flag epitope-tagged WT-FOXA1 or a C258A-FOXA1 mutant (or mock cells) treated with indicated concentrations of alkyne stereoprobes for 1 h, then lysed, and stereoprobe-labeled protein conjugated to a tetramethylrhodamine (TAMRA)-azide (RhN_3_) reporter tag by copper-catalyzed azide-alkyne cycloaddition (CuACC) chemistry followed by SDS-PAGE and in-gel fluorescence scanning. Also shown are treatments with inactive enantiomeric alkyne stereoprobes CJR-6B (6B) and WX-01-12 (12; see **Figures S1D**). Data are from a single experiment representative of two independent experiments. **H.** Gel-ABPP data showing the concentration-dependent, stereoselective blockade of CJR-6A reactivity with recombinant FOXA1 by WX-02-23 in HEK293T cells. Cells were treated with the indicated concentrations of WX-02-23 (23) or inactive enantiomer WX-02-43 (43) for 2 h followed by CJR-6A (5 µM, 1 h) and analyzed by gel-ABPP as described in Figure 1G. IB, immunoblots for the indicated proteins. Data are from a single experiment representative of two independent experiments.

We recently have shown that protein-directed ABPP experiments using alkyne-modified stereoprobes provide a complementary approach to cysteine-directed ABPP by enabling the discovery of proteins with liganded cysteines that reside on nonproteotypic peptides^38^. We accordingly synthesized alkyne analogs of WX-02-23 and WX-02-43 – CJR-6A and CJR-6B, respectively (**Figure 1D**) – and confirmed stereoselective enrichment of FOXA1 by CJR-6A and blockade of this enrichment by WX-02-23 in 22Rv1 cells (**Figure 1E**). These protein-directed ABPP experiments generally verified the limited global proteomic interactions of WX-02-23, identifying only a handful of stereoselectively liganded proteins beyond FOXA1, including the spliceosomal factor SF3B1, which is a previously characterized target of WX-02-23^37^ (**Figures 1E**, **F** and **S1C,** and **Supplementary Dataset S1**).

We next confirmed by gel-ABPP that CJR-6A engaged ectopically expressed WT-FOXA1 in a stereoselective (compared to inactive enantiomer CJR-6B) and site-specific (compared to a C258A-FOXA1 mutant) manner in transfected HEK293T cells (**Figure 1G**), and that this interaction was blocked in a concentration-dependent manner by WX-02-23, but not WX-02-43 (**Figure 1H**). WT-FOXA1 showed stronger reactivity with CJR-6A than the methyl ester analog WX-01-10 (**Figures 1G**, **S1D,** and **S1E**), which mirrored the respective greater engagement of FOXA1_C258 by WX-02-23 versus the propylamine analog WX-02-26 in cysteine-directed ABPP experiments (**Figures S1F**). This initial SAR analysis demonstrating both stereoselective (1*R*, 3*S* vs 1*S*, 3*R*) and chemoselective (morpholine vs methoxy/propylamine substituents) covalent binding to FOXA1_C258 indicated that the tryptoline acrylamide stereoprobes were interacting with a structurally ordered pocket in FOXA1.

Homology models derived from FOXA2-DNA and FOXA3-DNA crystal structures^40,41^ suggest that FOXA1_C258 is located on a dynamic loop within the Wing2 region that is not resolved in the FOXA2 structure^40^, but located near side chains that make contacts with the minor groove of DNA in the FOXA3 structure (**Figure 2A**). Interestingly, FOXA1_C258 is surrounded by diverse hotspot mutations in breast and prostate cancer that include both missense mutations and small in-frame deletions (some of which include C258) (**Figures 2A** and **B**). These cancer-related mutations have been shown to alter FOXA1 pioneering activity by modifying FOXA1-DNA interactions, leading to increases in FOXA1 chromatin mobility and the enhanced binding to and opening of new chromatin sites^20-22^. FOXA1_C258 and the surrounding region is also conserved in paralogs FOXA2 and FOXA3, but not in other FOX TFs (**Figures 2C** and **S2A**). We found that that all three FOXA paralogs were stereoselectively engaged by CJR-6A (**Figure 2D**); however, prostate cancer cell lines express and depend mainly on FOXA1, but not FOXA2 or FOXA3 (**Figures S2B**), and we therefore inferred that functional studies of tryptoline acrylamide stereoprobe-FOXA interactions performed in 22Rv1 cells would principally assess effects on FOXA1.

**Figure 2.**
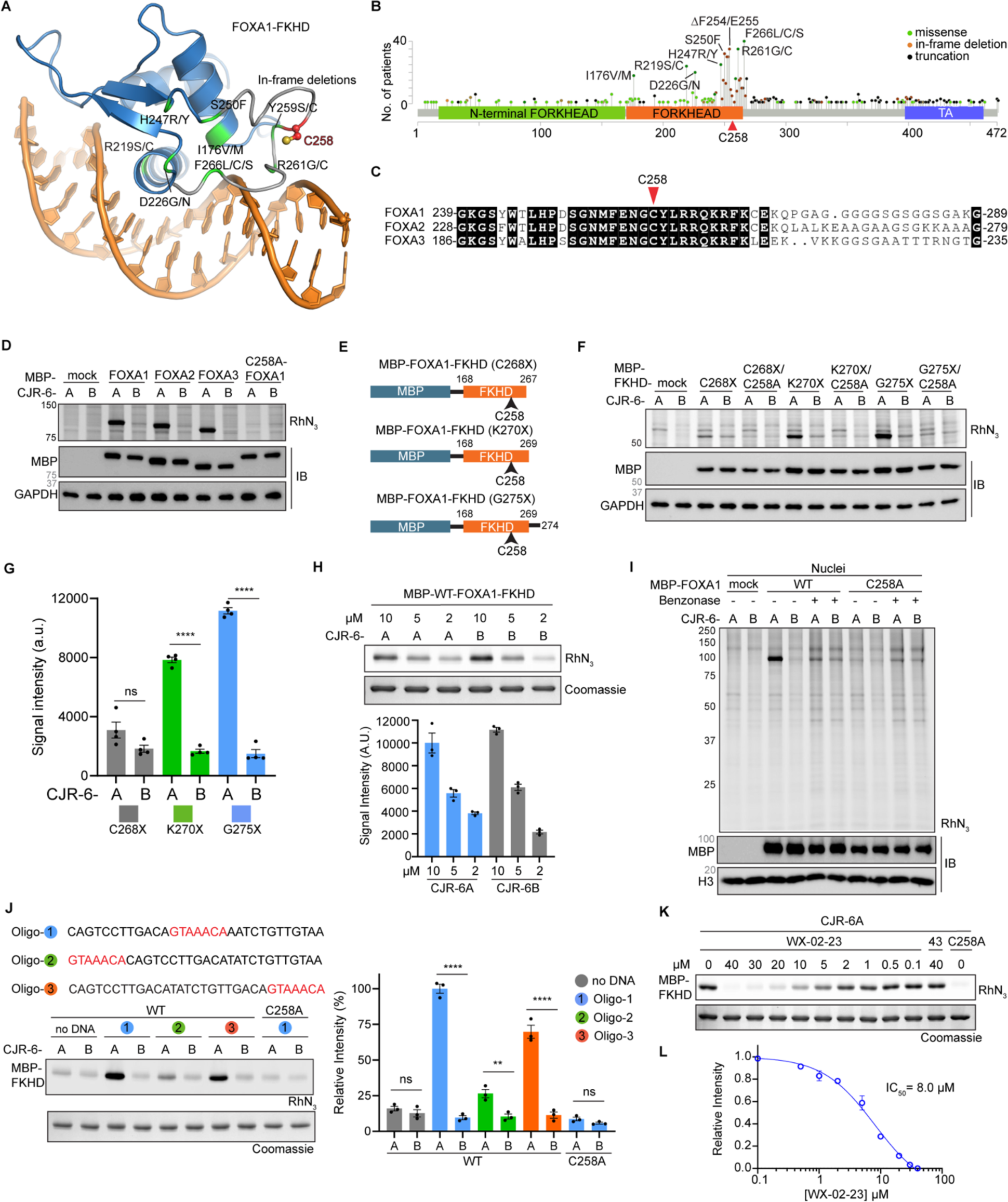
Stereoprobes engage FOXA1_C258 in a DNA-dependent manner. **A.** Homology model of a FOXA1 forkhead domain (FKHD)-DNA complex bound to a canonical DNA-binding motif. The model was generated using the FOXA3 FKHD-DNA structure as a template (PDB: 1VTN)^41^ and aligned with the FOXA2 FKHD-DNA structure (PDB: 5X07)^40^, which contains a canonical DNA-binding motif, to obtain a model with FOXA1-FKHD bound to the canonical motif. FOXA1_C258 is shown as a red stick, and representative cancer-related mutations are shown in green (missense) or grey (in-frame deletion). **B.** Lollipop plot of FOXA1 domain structure and relative position of cancer-related mutations generated from compiled data of breast and prostate cancer studies (cbioportal.org). C258 is indicated with a red arrow and high-frequency point mutations are labeled. **C.** Sequence alignment of human FOXA1, FOXA2, and FOXA3 around the Wing2 region with conserved residues highlighted in black and C258 marked in red. **D.** Gel-ABPP data showing stereoselective engagement of recombinant FOXA1, FOXA2, and FOXA3 by CJR-6A (5 µM; 1h). Data are from HEK293T cells recombinantly expressing maltose-binding protein (MBP)-tagged FOXA1 proteins treated with the indicated alkyne stereoprobes for 1 h and analyzed by gel-ABPP as described in Figure 1G. IB, immunoblots for the indicated proteins. Data are from a single experiment representative of two independent experiments. **E.** Design of MBP-FOXA1-FKHD fusion protein constructs differing in length of C-terminal extension of FOXA1-FKHD component. X designates the final C-terminal residue of each construct. **F.** Gel-ABPP data showing stereoselective and site-specific engagement of MBP-FOXA1-FKHD variants by CJR-6A. Data are from HEK293T cells recombinantly expressing the indicated MBP-FOXA1-FKHD constructs treated with CJR-6A or CJR-6B (10 µM in vitro, 1 h) and analyzed by gel-ABPP. IB, immunoblots for the indicated proteins. Data are from a single experiment representative of four independent experiments. **G.** Quantitation of gel-ABPP data shown in **F**. Data are average values ± SEM for four independent experiments; two-tailed t-test \*\*\*\**P* < 0.0001, ns = not significant. **H.** Gel-ABPP data showing lack of stereoselective engagement of purified MBP-WT-FOXA1-FKHD by CJR-6A. Purified protein (500 nM) was incubated with indicated concentrations of CJR-6A or CJR-6B (2 h) and analyzed by gel-ABPP. Upper image, representative gel-ABPP data; lower image, Coomassie blue stained gel. Lower graph, quantitation of gel-ABPP data. Data are average values ± SEM for three independent experiments. **I.** Effect of benzonase treatment on stereoselective engagement of MBP-FOXA1 by CJR-6A. Benzonase was added to HEK293T nuclear fraction for 30 min prior to addition of CJR-6A or CJR-6B (10 μM, 1 h), after which samples were analyzed by gel-ABPP. IB, immunoblots for the indicated proteins. Data are from a single experiment representative of two independent experiments. **J.** Effects of synthetic oligonucleotides containing a canonical FOXA1-FKHD binding motif on stereoselective and site-specific engagement of purified FOXA1 with CJR-6A. Individual oligonucleotides containing a canonical FOXA1 binding motif at varied positions (oligos-1, -2, -3; 0.5 µM) were incubated with purified MBP-FOXA1-FKHD (WT or C258A) (0.5 µM protein in DPBS supplemented with 2 mM MgCl_2_) for 30 min, followed by treatment with CJR-6A or CJR-6B (10 μM, 1 h) and analysis by gel-ABPP. Left, representative gel-ABPP and protein staining data. Right, quantitation of gel-ABPP data. Data are average values ± SEM for three independent experiments; two-tailed t-test \*\*\*\**P* < 0.0001, \*\**P* < 0.01, ns = not significant. **K.** Gel-ABPP data showing stereoselective blockade of CJR-6A engagement of purified MBP-FOXA1-FKHD by WX-02-23 in presence of oligo. Purified MBP-FKHD-FOXA1 or MBP-C258A-FKHD-FOXA1 proteins were incubated with oligo-1 and the indicated concentrations of WX-02-23 or WX-02-43 for 4 h, followed by treatment with CJR-6A (10 μM, 2 h) and analysis by gel-ABPP. Data are from a single experiment representative of two independent experiments. **L.** Quantitation of gel-ABPP from **K**. Data are average values ± SEM for two independent experiments.

### Covalent ligands engage FOXA1_C258 in a DNA-dependent manner

We next sought to characterize how tryptoline acrylamide stereoprobes engaged FOXA1. We generated maltose binding protein (MBP)-fused WT and C258A mutant FOXA1-FKHD constructs of varying C-terminal lengths for ectopic expression and analysis by gel-ABPP in HEK293T cells (**Figure 2E**). These experiments revealed that CJR-6A stereoselectively and site-specifically engaged the MBP-WT-FOXA1-FKHD proteins truncated at K270 (K270X) and G275 (G275X), but not at C268 (C268X), with the G275X construct (referred to hereafter as MBP-FOXA1-FKHD) showing the strongest overall reactivity (**Figures 2F, G**).

We next purified the MBP-WT-FOXA1-FKHD protein from transfected suspension HEK293 cells (**Figures S2C**) and found, to our surprise, that the purified protein failed to stereoselectively react with CJR-6A across a range of test concentrations (**Figure 2H**). We speculated that additional factors may be needed to support the specific reactivity of FOXA1 with tryptoline acrylamide stereoprobes in cells that were lost during protein purification. We first confirmed that a full-length MBP-WT-FOXA1 fusion protein reacted stereoselectively and site-specifically with CJR-6A in the nuclear fractions of transfected HEK293T cells (**Figures S2D**), where FOXA1 is known to reside on chromatin^42,43^. We then found that adding purified MBP-WT-FOXA1-FKHD to the nuclear lysate of untransfected HEK293T cells restored stereoselective and site-specific reactivity with CJR-6A (**Figures S2E**). Since TFs often undergo structural change upon binding to DNA^44^, we next tested the effects of the universal nuclease benzonase and found that it blocked the stereoselective reactivity of MBP-WT-FOXA1 or MBP-WT-FOXA1-FKHD with CJR-6A (**Figures 2I** and **S2F, G**). Benzonase treatment also blocked the stereoselective reactivity of CJR-6A with FOXA2 and FOXA3 (**Figures S2H**), as well as 2xStrep-Flag epitope-tagged FOXA1 (**Figures S2I**).

We interpreted the effects of benzonase to indicate that tryptoline acrylamides may bind FOXA1 in a DNA-dependent manner. Consistent with this conclusion, incubation of purified MBP-FOXA1-FKHD protein with 31-bp oligonucleotide duplexes containing a canonical binding motif for FOXA1 (GTAAACA) restored stereoselective and site-specific reactivity with CJR-6A (**Figure 2J**). The most robust interactions between MBP-FOXA1-FKHD and CJR-6A occurred in the presence of oligonucleotides bearing central and 3’-end canonical binding motifs, whereas only weak engagement was observed with an oligonucleotide displaying the motif at the 5’ end. DNA titration studies revealed a concentration-dependent increase in stereoselective reactivity between CJR-6A and MBP-FOXA1-FKHD protein that could be visualized across a range of oligonucleotide concentrations (0.2-5 µM) spanning the concentration of MBP-FOXA1-FKHD (0.5 µM) (**Figures S2J**). Finally, we confirmed that pre-treatment with WX-02-23 stereoselectively blocked CJR-6A reactivity with purified MBP-FOXA1-FKHD protein in the presence of oligonucleotide with an IC_50_ value of 8 µM (**Figures 2K, L**).

Having discovered that tryptoline acrylamides specifically react with FOXA1 in the presence of DNA, we next set out to test how these covalent ligands might impact FOXA1-DNA interactions.

### Covalent ligands enhance FOXA1-DNA binding *in vitro*

We quantified FOXA1-DNA interactions using two complementary *in vitro* assays. In the first approach adapted from previous methods^44-47^, we measured binding of a Green Renilla luciferase-tagged FOXA1 (Luc-FOXA1) from nuclear lysates of transfected HEK293T cells with a biotinylated oligonucleotide duplex (31-bp) harboring a canonical FOXA1-binding motif (GTAAACA) immobilized on streptavidin-coated plates (**Figures S3A**). In the second approach, we adapted a NanoBRET (Nano-Bioluminescence Resonance Energy Transfer) assay^48^, where lysates from HEK293T cells expressing NanoLuc (NLuc)-tagged FOXA1 or FOXA1-FKHD proteins were incubated with a TAMRA-tagged oligonucleotide (32-bp) containing a centrally positioned canonical FOXA1-binding motif (or a control oligonucleotide lacking this motif) and BRET signal was measured after NLuc substrate addition (**Figures 3A, B**). Both assays showed strong and specific signals for FOXA1 and FOXA1-FKHD binding to the canonical motif oligonucleotide in comparison to protein (Luc or NLuc protein) or oligonucleotide (non-canonical) controls (**Figures 3C** and **S3B, C)**. The C258A-FOXA1 mutant protein showed a similar, albeit slightly weaker profile of specific binding to the canonical motif oligonucleotide (**Figures 3C** and **S3B, C**), indicating that mutation of C258 did not, on its own, substantially alter FOXA1 interactions with DNA.

**Figure 3.**
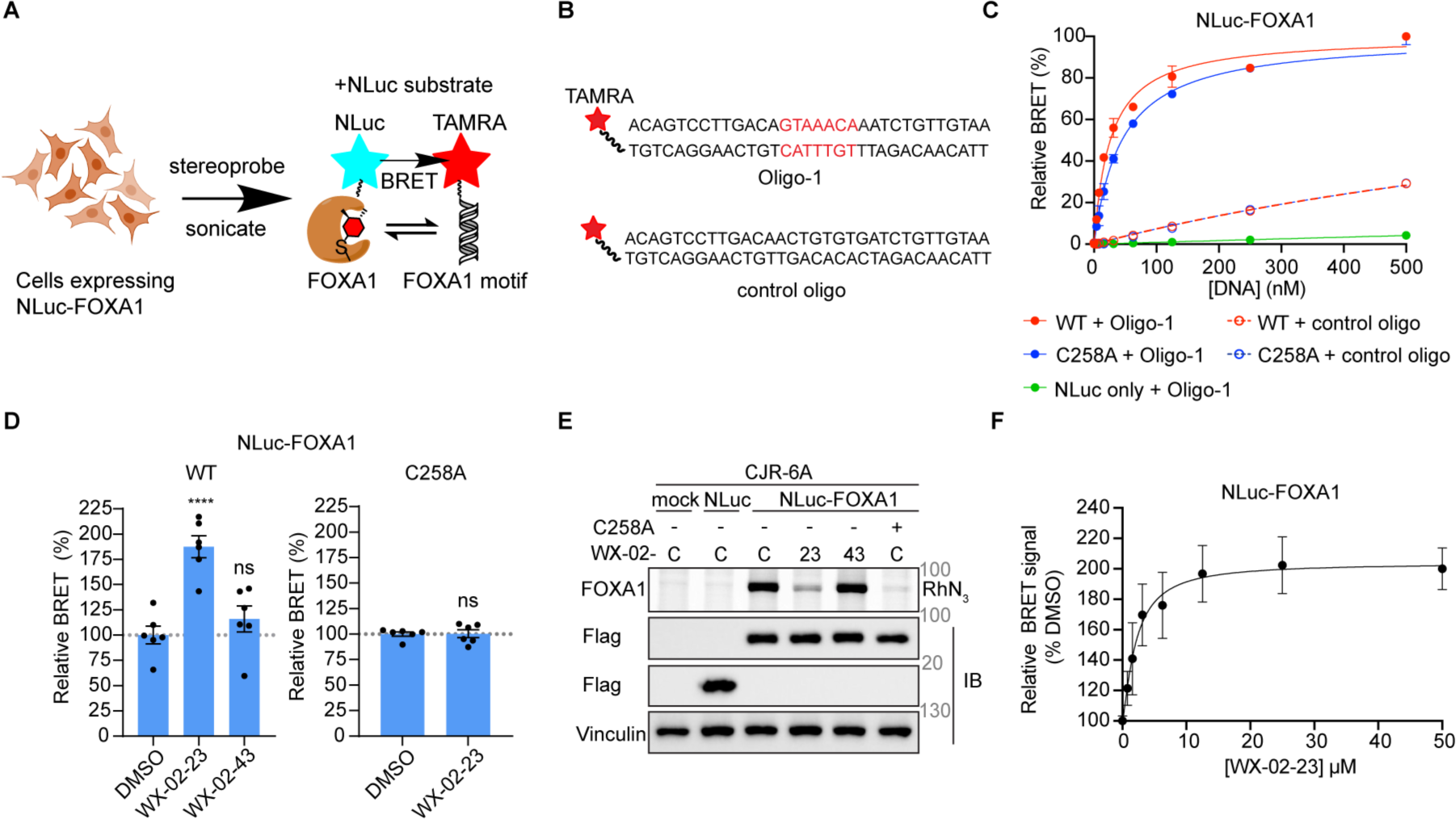
Stereoprobes enhance FOXA1-DNA interactions *in vitro*. **A.** Schematic for nanoBRET-based oligonucleotide-binding assay to measure stereoprobe effects on FOXA1-DNA interactions. HEK293T cells recombinantly expressing a FOXA1-NanoLuciferase (NLuc) fusion protein (C-terminally tagged) or NLuc-FOXA1-FKHD (N-terminally tagged) were treated with WX-02-23 or WX-02-43 (20 µM, 3 h), after which the cells were lysed and incubated with a TAMRA-tagged fluorescent oligo-1 variant for 20 min. NLuc substrate was then added and BRET ratio (ratio of emission from > 610nm (longpass) to 460 ± 80 nm (bandpass)) was measured. **B.** Sequences of TAMRA-conjugated FOXA1-binding DNA oligonucleotide with canonical FOXA1-binding motif in red (oligo-1; above) and control DNA oligonucleotide lacking the canonical FOXA1-binding motif (below) used in the nanoBRET assay. **C.** Binding curves for oligo-1 and control oligo with NLuc-FOXA1 fusion proteins (red:WT and blue:C258A) assayed in lysates of transfected HEK293T cells. Dashed lines indicate binding profile of WT-(red) or C258A-(blue) protein with control oligo, and green indicates oligo-1 binding to a NLuc control protein (also expressed in HEK293T cells; see Figure 3E). NanoBRET signals are reported after subtraction of background signals from samples lacking oligonucleotides. Data are average values ± SEM from 2 independent experiments. **D.** Effects of WX-02-23 and WX-02-43 on NLuc-FOXA1 interactions with oligo-1. HEK293T cells expressing NLuc-WT- or NLuc-C258A-FOXA1 proteins were treated with WX-02-23 or WX-02-43 (20 µM, 3 h), or DMSO, followed by cell lysis and incubation with oligo-1 (15 nM, 20 min), and nanoBRET analysis. NanoBRET data were normalized to DMSO controls for each protein. Data are average values ± SEM; N = 6 per group. Two-tailed t-test \*\*\*\**P* <0.0001 compared to DMSO control. **E.** Gel-ABPP data confirming stereoselective and site-specific engagement of NLuc-WT-FOXA1 by stereoprobes. HEK293T cells expressing NLuc-WT- or NLuc-C258A-FOXA1 (or NLuc control) proteins were treated with WX-02-23 (23) or WX-02-43 (43) (20 μM, 2 h), followed by CJR-6A (5 μM, 1 h), and analyzed by gel-ABPP. C= DMSO control. IB, immunoblots for indicated proteins. Data are from a single experiment representative of two independent experiments. **F.** Concentration-dependent effects of WX-02-23 on nanoBRET signal for NLuc-WT-FOXA1 with oligo-1. HEK293T cells expressing NLuc-WT-FOXA1 were treated with the indicated concentrations of WX-02-23 for 3 h, followed by cell lysis and incubation with oligo-1 (15 nM, 20 min), and nanoBRET analysis. Data are average values ± SEM and presented as the relative maximum percentage increase of nanoBRET signal in WX-02-23-treated samples over DMSO-treated control samples (where DMSO-treated controls are set to 100%); N = 4 per group.

We found that pre-treatment of HEK293T cells expressing the respective Luc/NLuc-FOXA1 and FOXA1-FKHD constructs with WX-02-23 (20 μM, 3 h) led to enhanced FOXA1 interactions with DNA, and this effect was stereoselective (not observed in cells pre-treated with WX-02-43) and site-specific (not observed in Luc/NLuc-C258A-FOXA1-transfected cells treated with WX-02-23) (**Figures 3D** and **S3D-F**). We also confirmed by gel-ABPP that recombinantly expressed Luc/NLuc-FOXA1 proteins showed the expected stereoselective and site-specific reactivity with the CJR6A/WX-02-23 stereoprobes (**Figures 3E** and **S3G-J**). Using the NanoBRET assay, we measured EC_50_ values for WX-02-23-dependent enhancement of FOXA1 (EC_50_ = 2 µM) and FOXA1-FKHD (EC_50_ = 12 µM) binding to DNA (**Figures 3F** and **S3K**), which were similar to the IC_50_ value for WX-02-23 engagement of FOXA1-FKHD domain determined by gel-ABPP (**Figures 2K, L**).

In summary, using two complementary biochemical assays, we found that tryptoline acrylamide stereoprobes that covalently react with C258 of FOXA1 promote the stereoselective and site-specific binding of this protein to DNA.

### Covalent ligands remodel the pioneering activity of FOXA1 in cancer cells

Imaging studies, including single molecule fluorescence microscopy, have indicated that ectopically expressed FOXA1 interacts dynamically with chromatin^42,43,49-52^, suggesting that acute treatment with small molecules may have the potential to impact DNA interactions of endogenous FOXA1 in cells. We accordingly next performed FOXA1 ChIP-seq (chromatin immunoprecipitation followed by sequencing) experiments in 22Rv1 cells treated with WX-02-23 or WX-02-43 (20 µM) for 3 h. In these studies, we also included an additional compound, pladienolide B (PladB)^53^, which is a natural product that binds to the same pocket in the spliceosome as WX-02-23 and phenocopies its effects on splicing^37^. PladB thus provided a control for potential off-target effects of WX-02-23 through engagement of SF3B1 (**Figure 1E**). We did not observe by ChIP-Seq a gross change in the strength of FOXA1 binding to the genome of 22Rv1 cells treated with WX-02-23 (**Figures S4A**, **B,** and **Supplementary Dataset S2**), but instead found that this compound caused a striking redistribution of FOXA1 binding sites across chromatin, as reflected in a much lower correlation with DMSO-treated cells (PCC = 0.62) compared to the correlation displayed by cells treated with control compounds WX-02-43 (PCC = 0.87) or PladB (PCC = 0.94) (**Figure 4A**). The impact of WX-02-23 on FOXA1-chromatin interactions was bidirectional, with substantial numbers of FOXA1 binding sites showing either increased or decreased signals in comparison to DMSO treatment (**Figures 4A**, **B**). In contrast, many fewer changes in FOXA1 binding were caused by WX-02-43 or PladB (**Figures 4A**, **B**). The stereoselective changes in FOXA1 binding in WX-02-23-treated cells occurred without apparent bias to genomic feature, with WX-02-23-responsive and unresponsive FOXA1 binding sites showing equivalent frequencies of localization to intergenic regions (∼50%), intronic regions (∼45%), and promoters and exons (<5%) (**Figures S4C**).

**Figure 4.**
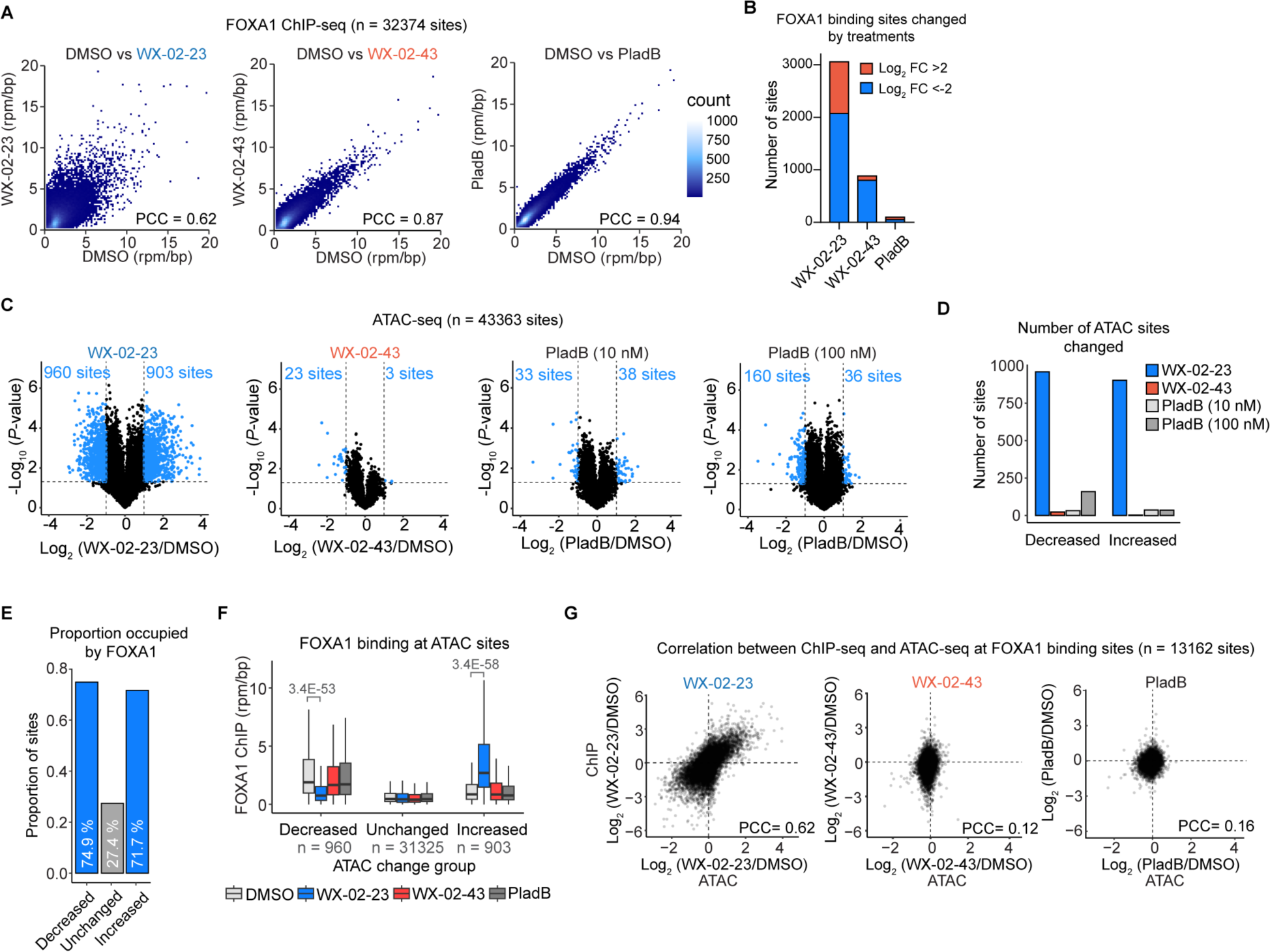
WX-02-23 redirects the pioneering activity of FOXA1 in prostate cancer cells. **A.** Density plots showing correlations of FOXA1 binding intensity for indicated compound treatments (20 μM for each stereoprobe; 10 nM for PladB, 3 h) versus DMSO control in 22Rv1 cells. Density scale is shown on the right. PCC = Pearson’s correlation coefficient. **B.** Number of FOXA1 binding sites with substantial changes induced by compound treatment relative to DMSO control (Log_2_FC < -2 (blue) or Log_2_FC > 2 (red)). WX-02-23, Log_2_FC < -2: 2076 sites, Log_2_FC > 2: 982 sites; WX-02-43, Log_2_FC < -2: 799 sites, Log_2_FC > 2: 86 sites; PladB, Log_2_FC < -2: 48 sites, Log_2_FC > 2: 50 sites. Sites with substantial ChIP signal (rpm/bp > 1) in at least 1 treatment condition were used for analyses. **C.** Chromatin accessibility (ATAC-seq) changes induced by compound treatments (20 µM for each stereoprobe and 10 or 100 nM for PladB) compared to DMSO control in 22Rv1 cells. ATAC-seq sites with substantial ((|log_2_FC| > 1) and significant (*P* < 0.05) changes relative to DMSO are shown as blue dots. Sites with substantial ATAC-seq signal (rpm/bp > 1) in at least 1 treatment condition were used for all analyses. Data are average values from three independent experiments, and *P* values were determined by two-tailed Welch’s *t*-test. The number of significantly changed sites for each treatment group are shown in blue text. **D.** The number of ATAC sites substantially and significantly changed for each compound treatment group (Increased: log_2_FC > 1, *P* < 0.05 or Decreased: log_2_FC < -1*, P* < 0.05). *P* values were determined by two-tailed Welch’s *t*-test. **E.** Proportion of ATAC-seq sites occupied by FOXA1 as determined by ChIP-Seq, grouped by WX-02-23-induced ATAC-seq changes relative to DMSO (decreased, log_2_FC < -1 and *P* < 0.05; increased, log_2_FC > 1 and *P* < 0.05); unchanged, |log_2_FC| < 1 and *P* > 0.05). *P* values were determined by two-tailed Welch’s *t*-test. **F.** FOXA1 ChIP-seq signal at ATAC-seq sites grouped by WX-02-23-induced changes relative to DMSO (decreased, log_2_FC < -1 and *P* < 0.05, *N* = 960 sites; increased, log_2_FC > 1 and *P* < 0.05, *N* = 903 sites; unchanged, |log_2_FC| < 1 and *P* > 0.05*, N* = 32,125 sites). *P* values were determined by two-tailed Welch’s *t*-test between DMSO and WX-02-23. **G.** Correlations between FOXA1 ChIP-seq changes and ATAC-seq changes induced by the indicated compound treatments in 22Rv1 cells. FOXA1-sites with substantial (rpm/bp > 1) FOXA1 ChIP-seq and ATAC-seq signal in at least 1 treatment condition were included for this analysis. PCC = Pearson’s correlation coefficient.

To determine whether the WX-02-23-induced redistribution of FOXA1 binding led to changes in chromatin accessibility, we performed ATAC-seq (assay for transposase-accessible chromatin-sequencing) experiments in 22Rv1 cells^54^. Out of >40,000 quantified sites, >1800 substantial (Log_2_FC < -1 or > 1 compared to DMSO) and significant (*P* < 0.05) changes in ATAC-seq were identified in WX-02-23-treated cells, which contrasted with the minimal perturbations observed in WX-02-43- and PladB-treated cells (< 200 changing sites total) (**Figure 4C** and **Supplementary Dataset S2**). Similar to the FOXA1 ChIP-seq changes caused by WX-02-23, the ATAC-Seq changes were bidirectional, reflected in nearly equivalent numbers of decreased (960 sites) and increased (903 sites) sites (**Figures 4C**, **D**). Notably, the ATAC-seq sites decreased or increased by WX-02-23 both showed a striking enrichment in direct FOXA1 binding sites compared to unchanged ATAC-seq sites (**Figure 4E**). The ATAC-seq sites decreased or increased by WX-02-23 also showed comparatively higher and lower basal FOXA1-ChIP signal in DMSO-treated cells, and these FOXA1-ChIP signals were decreased and increased, respectively by WX-02-23, but not WX-02-43 or PladB (**Figure 4F**). Focusing only on the ATAC-seq signals at FOXA1 binding sites mapped by ChIP-seq, we confirmed the bidirectional pattern of accessibility changes caused by WX-02-23 with negligible effects of WX-02-43 or PladB (**Figures S4D**, **E**). We also observed a robust correlation in the directionality of FOXA1-binding (ChIP-seq) and chromatin accessibility (ATAC-seq) changes caused by WX-02-23 (PCC=0.62), while WX-02-43 and PladB (PCC=0.12 and 0.16, respectively) did not show such a pattern (**Figure 4G**). We interpret these data to indicate that the bidirectional redistribution of FOXA1 binding to chromatin promoted by WX-02-23 results in corresponding changes in accessibility at these sites.

As an additional test whether the WX-02-23-induced stereoselective changes in chromatin binding and accessibility were due to direct covalent engagement of FOXA1, we performed FOXA1 ChIP-seq and ATAC-seq experiments in 22Rv1 cells stably expressing recombinant 2xStrep-Flag-tagged WT- or C258A-FOXA1 under a doxycycline-inducible promoter, which allowed for several-fold higher expression of the recombinant FOXA1 proteins compared to endogenous FOXA1 (**Figures 5A, S5A, and Supplementary Dataset S2**). To further minimize the potential contributions of spliceosome interactions by WX-02-23, we introduced the WT- and C258A-FOXA1 proteins into 22Rv1 cells also expressing a Y36C-PHF5A mutant, which we and others have shown to display resistance to SF3B1 ligands, including WX-02-23 and PladB^37,55^. We found by ChIP-seq analysis that WX-02-23 (20 µM, 3h) cause a much greater redistribution of FOXA1-binding sites in cells expressing WT-FOXA1 (PCC = 0.62 vs DMSO-treated cells) compared to cells expressing C258A-FOXA1 (PCC = 0.84 vs DMSO-treated cells) (**Figure 5B**), and ATAC-seq analysis showed a similar outcome, with WX-02-23 decreasing the chromatin accessibility of more than 1300 sites in WT-FOXA1-expressing cells, while minimally affecting C258A-FOXA1 expression cells (∼60 sites decreasing) (**Figure 5C**). PladB displayed minimal effects on chromatin accessibility in either WT- or C258A-FOXA1-expressing cells (< 50 sites changed; **Figures S5B**).

**Figure 5.**
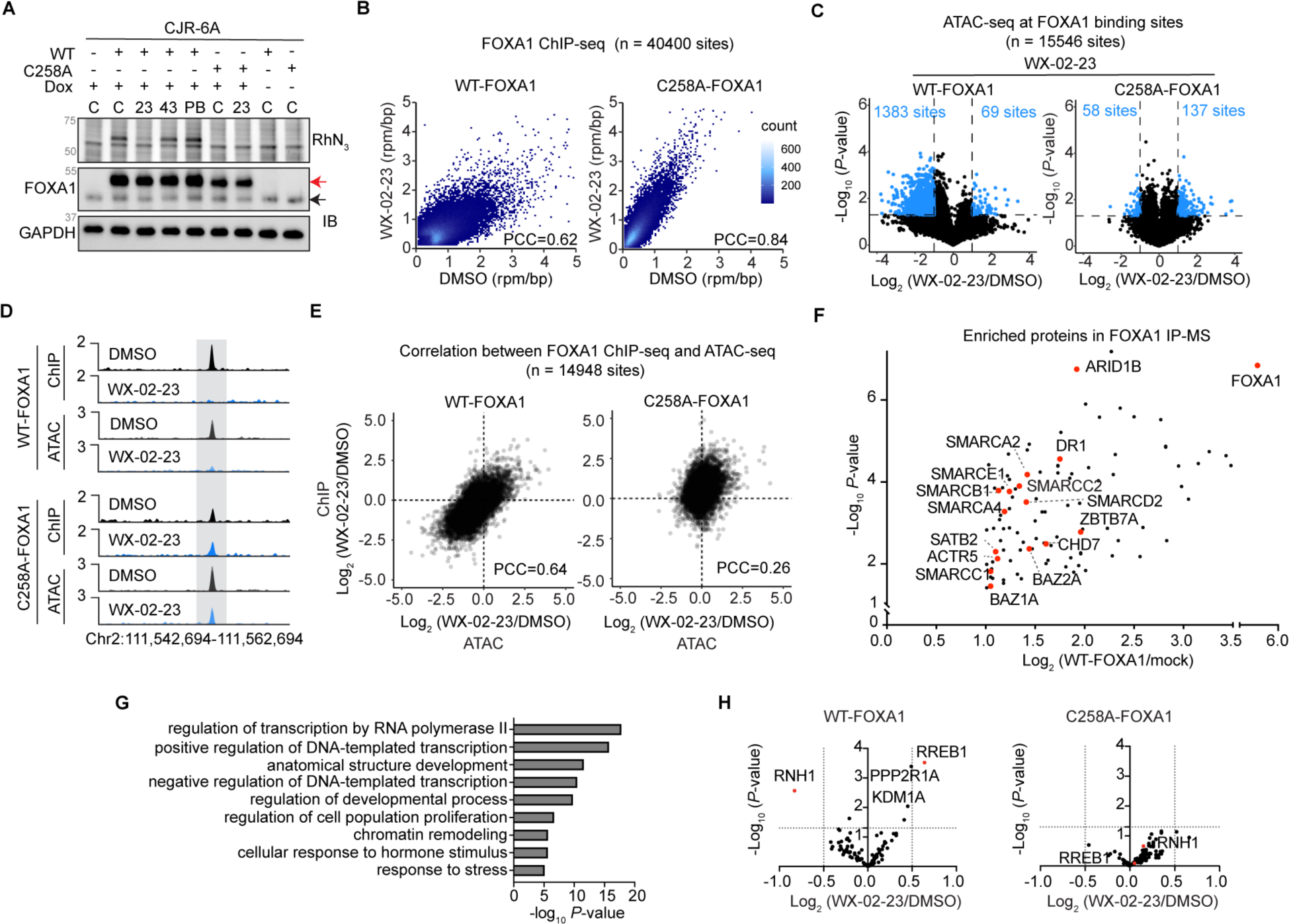
WX-02-23 effects on FOXA1-chromatin interactions depend on engagement of C258. **A.** Gel-ABPP and immunoblot (IB) analysis of doxycycline-induced expression of 2xStrep-flag epitope-tagged WT- and C258A-FOXA1 in 22Rv1 cells stably expressing a Y36C-PHF5A protein that impedes spliceosome binding by WX-02-23. Cells were treated with 0.5 μg/ml doxycycline for 24 h followed by the indicated compounds, or DMSO control (C), for 2 h (20 μM WX-02-23 (23) and WX-02-43 (43); 100 nM PladB (PB)). Cells were then treated with CJR-6A (5 µM, 1 h) and analyzed by gel-ABPP. For IB, the red arrow indicates recombinantly expressed WT- or C258A-FOXA1 and the black arrow indicates endogenous FOXA1. **B.** Density plots showing correlations (DMSO vs WX-02-23 (20 μM, 3h)) of FOXA1 binding intensity in 22Rv1 Y36C-PHF5A cells recombinantly expressing WT-(left) or C258A-FOXA1 (right) via doxycycline induction. Density scale is shown on the right. PCC = Pearson’s correlation coefficient. **C.** Chromatin accessibility (ATAC-seq) changes induced by WX-02-23 (20 μM, 3h) at FOXA1 binding sites in 22Rv1 Y36C-PHF5A cells recombinantly expressing WT- or C258A-FOXA1 via doxycycline induction. ATAC-seq sites with substantial ((|log_2_FC| > 1) and significant (*P* < 0.05) changes relative to DMSO are shown as blue dots. Sites with ATAC-seq signal below 0.5 rpm/bp were excluded from the analysis. Data are averaged values from two independent experiments. *P*-values were determined by two-tailed Welch’s *t*-test. **D.** Representative gene track showing that WX-02-23-induced changes in FOXA1 binding and chromatin accessibility are rescued in cells recombinantly expressing C258A-FOXA1. Y-axis in rpm/bp. **E.** Correlations between FOXA1 ChIP-seq and ATAC-seq changes induced by WX-02-23 in 22Rv1 Y36C-PHF5A cells recombinantly expressing WT- or C258A-FOXA1. FOXA1-sites with substantial signal (rpm/bp > 0.5) are included for this analysis. PCC = Pearson’s correlation coefficient. **F.** Proteins substantially and significantly enriched in anti-Flag immunoprecipitation (IP)-MS experiments performed on 22Rv1 cells expressing 2xStrep-Flag-tagged WT-FOXA1 ((WT-FOXA1/mock) > 2, *P* < 0.05) following doxycycline treatment (24 h) compared to mock 22Rv1 cells. Chromatin remodeling proteins are shown in red dots. Data are average values. N = 4-6 per group. *P* values were determined by two-tailed Welch’s *t*-test. **G.** Gene ontology enrichment for proteins significantly enriched in anti-Flag IP-MS experiments performed on 22Rv1 cells expressing 2xStrep-Flag-WT-FOXA1 vs mock cells (See Figure 5F). *P*-value threshold of 10^-5^ was used. **H.** Volcano plots comparing enriched proteins from anti-Flag IP-MS experiments performed on 22Rv1 cells treated with 2xStrep-Flag-WT- or C258A-FOXA1 and treated with WX-02-23 (20 µM, 3 h) or DMSO. Red dots mark substantially (|Log_2_FC| > 1) and significantly (*P* < 0.05) changing proteins (RNH1 and RREB1) in WX-02-23-treated WT-FOXA1 cells, and the location of these proteins is also marked for WX-02-23-treated C258A-FOXA1 cells. Data are average values. N = 4-9 per group. *P* values were determined by two-tailed Welch’s *t*-test.

In contrast to the bidirectional changes in chromatin accessibility caused by WX-02-23 in parental 22Rv1 cells (**Figure 4C**), this compound mainly caused decreases in chromatin accessibility in 22Rv1 cells ectopically expressing WT-FOXA1 (**Figures 5C**, **D**), which we speculate may reflect the greater basal site occupancy for FOXA1 in cells that overexpress this protein. Potentially consistent with this hypothesis, doxycycline-induced expression led to thousands of new FOXA1-binding sites across the genome that correlated with increases in chromatin accessibility at these sites **(Figures S5C**). Conversely, the much lower, but similar number of chromatin sites showing increased accessibility in WX-02-23-treated WT-FOXA1- and C258A-FOXA1-expressing cells could reflect the action of this compound on endogenous WT-FOXA1, which remained present in both cell models. Even with the increased number of chromatin binding sites found in WT-FOXA1-expressing cells, we still observed a good correlation between the ChIP-seq and ATAC-seq effects of WX-02-23 in this cell model (PCC = 0.64) compared to C258A-FOXA1-expressing cells (PCC = 0.26) (**Figure 5E**). Finally, it is also noteworthy that WX-02-23 not only caused many fewer chromatin accessibility changes in C258A-FOXA1-expressing cells in comparison to WT-FOXA1-expressing cells (**Figures 5C**, **5D**), but also in comparison to parental 22Rv1 cells (**Figure 4C**), which may be explained by the overexpressed recombinant C258A-FOXA1 protein displacing endogenous FOXA1 from many binding sites across the genome, rendering them insensitive to WX-02-23 treatment. These data collectively support that the WX-02-23-induced changes in FOXA1 chromatin binding and chromatin accessibility are due to direct covalent modification of C258 in this protein.

How pioneer TFs like FOXA1 decompact chromatin remains unclear, but this process may involve additional interacting proteins, such as ATP-dependent chromatin remodelers, that also support the maintenance of stable open chromatin in cells^3,56-60^. We took advantage of our 22Rv1 cell models expressing WT-FOXA1 and C258A-FOXA1 to map the interactome of FOXA1 and the effect of WX-02-23 on these protein-protein interactions. We specifically performed immunoprecipitation-mass-spectrometry (IP-MS) studies on WT-FOXA1 and C258A-FOXA1-expressing 22Rv1 cells treated with DMSO, WX-02-23 (20 µM, 3 h), WX-02-43 (20 µM, 3 h), or PladB (100 nM, 3 h) and identified more than 100 proteins that were substantially (fold-change > 2) in and significantly (*P* < 0.05) enriched in anti-FLAG IPs from WT-FOXA1-expressing cells compared to mock-transfected 22Rv1 cells (**Figure 5F** and **Supplementary Dataset S1**). A similar interactome was observed for both WT- and C258A-FOXA1 (**Figures S5D**). The FOXA1-interacting proteins were enriched for processes regulating transcription and chromatin remodeling (**Figure 5G**), including, for instance, members of the SWI/SNF (SWItch/Sucrose Non-Fermentable) complex (**Figure 5F** and **Supplementary Dataset S1**). This FOXA1 interactome was largely unperturbed by WX-02-23 with the exception of two proteins – Ribonuclease Inhibitor 1 (RNH1) and Ras Responsive Element Binding Protein 1 (RREB1) – that were decreased and increased in enrichment, respectively, and these effects were stereoselective and site-specific (**Figures 5H** and **S5E**). Our IP-MS data thus indicate that tryptoline acrylamide stereoprobes targeting FOXA1_C258 largely leave the FOXA1 interactome intact, supporting that these protein complexes should be competent for chromatin remodeling at sites to which FOXA1 is chemically redirected.

### Covalent ligands alter pioneer activity of FOXA1 by relaxing its DNA binding motif

We wondered whether the bidirectional effects of WX-02-23 on chromatin binding of FOXA1 might provide insights into its pioneer function with a temporal resolution that has, to date, been difficult to achieve in cellular systems. Here, we hypothesized that, if WX-02-23 induced *de novo* binding sites on chromatin for endogenous FOXA1 in 22Rv1 cells, then corresponding increases in chromatin accessibility at these *de novo* sites would provide support for a pioneer function (i.e., binding to largely compacted sites of the genome and greatly increasing accessibility at these sites). Through comparative mean-average plots, we observed many sites of increased chromatin accessibility in WX-02-23-treated 22Rv1 cells that had very low basal ATAC-seq signal in DMSO-treated control cells (**Figures 6A** and **S6A**) as well as in WX-02-43- and PladB-treated control cells (**Figures 6A** and **S6A**, **B**). We next grouped FOXA1-binding sites by the changes induced by WX-02-23 (substantially decreased: Log_2_FC < -2; unchanged: |Log_2_FC| < 1; substantially increased: Log_2_FC > 2; in comparison to DMSO-treated cells) and examined ChIP- and ATAC-seq signals for each group across all treatment conditions. This analysis revealed that sites of substantially increased FOXA1 binding in WX-02-23-treated cells harbored ChIP- and ATAC-seq signals that were as strong or stronger than those from unchanged sites in the DMSO control (i.e., compare signals for increased sites in WX-02-23 treatment group to signals for unchanged sites in DMSO treatment group; **Figure 6B**). We accordingly infer that sites of increased FOXA1 binding and chromatin accessibility were often equivalent in signal strength to endogenous FOXA1 binding sites in prostate cancer cells (as opposed to being low occupancy sites). Remarkably, we also found that, for sites with substantially decreased FOXA1 binding in WX-02-23-treated cells, the ChIP-seq signal was reduced in many instances to undetectable levels (**Figure 6B**). Importantly, for DMSO-treated cells, the signals at these sites were comparable to unchanged sites, indicating that WX-02-23 can evict FOXA1 from strong FOXA1-binding sites (**Figure 6B**, **C**). Inspection of individual sites of increased and decreased peaks in WX-02-23-treated cells supported these global observations, with examples of (i) FOXA1 being localized to sites of largely inaccessible chromatin with virtually no basal FOXA1 binding and causing increases in both ATAC- and ChIP-seq signals equivalent to native peaks located nearby (**Figure 6C**); and (ii) robustly accessible sites with abundant FOXA1 binding being diminished to near-undetectable ATAC-seq and ChIP-seq signals (**Figure 6C**).

**Figure 6.**
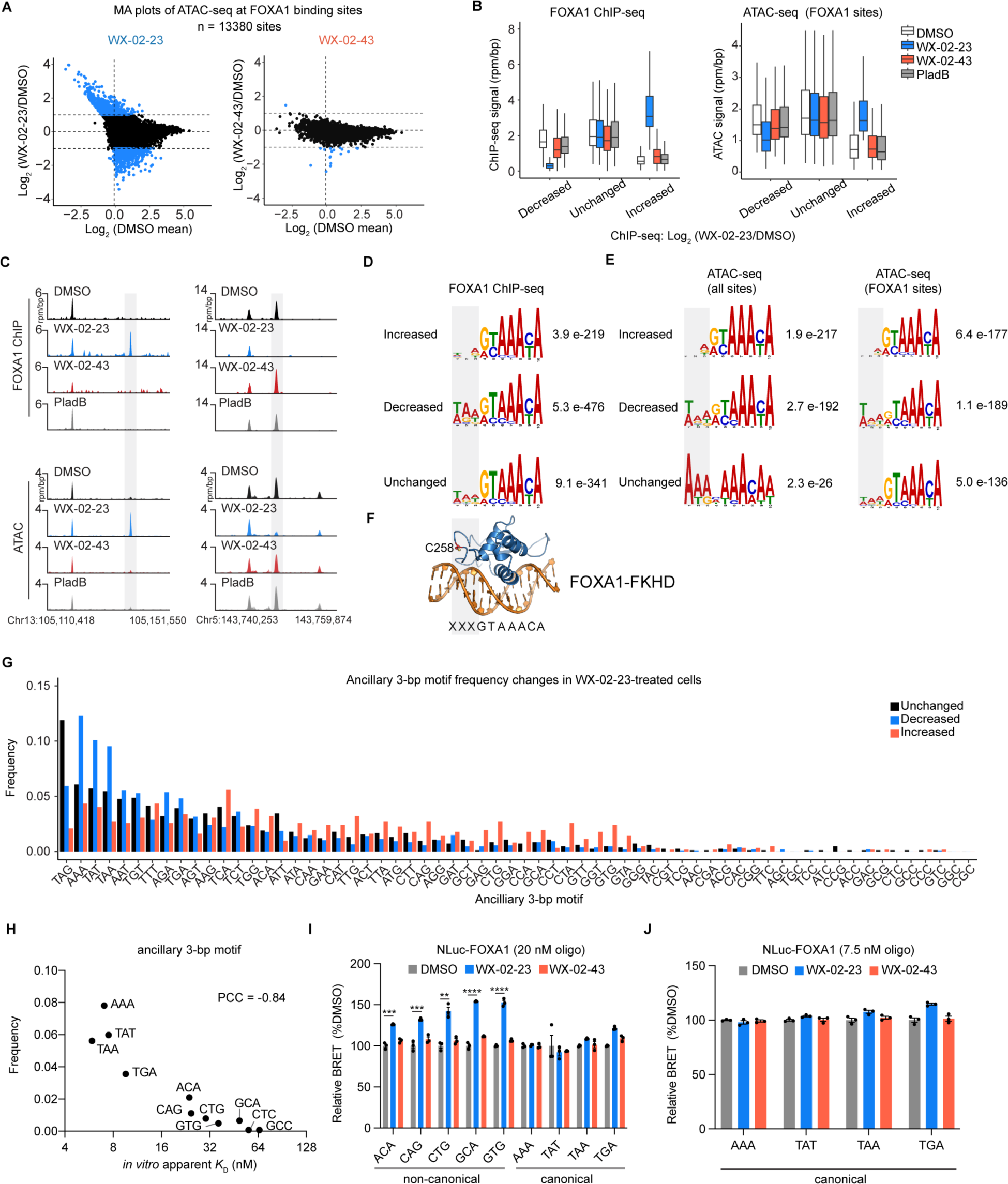
WX-02-23 remodels chromatin interactions of FOXA1 by relaxing its DNA-binding motif preference. **A.** Mean-average (MA) plots showing chromatin accessibility (ATAC-seq) changes induced by WX-02-23 or WX-02-43 (20 μM, 3 h) at ChIP-seq-defined FOXA1-binding sites in 22Rv1 cells (see Figure 4A). The vertical line represents 10th percentile of DMSO mean. FOXA1-binding sites with substantial ATAC signal (rpm/bp > 1) in at least 1 treatment condition were used for all analyses. ATAC-seq sites with substantial and significant changes are shown as blue dots (|log_2_FC| > 1, *P* < 0.05, as determined in **Figures S4D**) Data are average values from three independent experiments. See **Figures S6B** for PladB treatments. **B.** FOXA1 ChIP-seq (left) and ATAC-seq (right) signals (y-axis; rpm/bp) at FOXA1-binding sites grouped by FOXA1 ChIP-seq changes induced by WX-02-23 relative to DMSO (x-axis). Decreased sites: Log_2_(FC) < -2, 528 sites; unchanged sites: -1 < log_2_(FC) < 1, 9,475 sites; increased sites: log_2_(FC) > 2, 329 sites. ATAC-seq data are average values from three independent experiments. White: DMSO, WX-02-23: blue, WX-02-43: red, PladB: grey. FOXA1-sites with substantial (rpm/bp > 1) FOXA1 ChIP-seq and ATAC-seq signal in at least 1 treatment condition were included for this analysis. **C.** Representative gene tracks showing WX-02-23-induced stereoselective increases (left tracks; gray highlight) and decreases (right tracks; gray highlight) in FOXA1-chromatin interactions (FOXA1 ChIP-seq) and corresponding changes in chromatin accessibility (ATAC-seq). For ATAC-seq, all three replicates are included and overlaid in the track. **D.** *De novo* motif discovery analysis of FOXA1-binding sites. Separate analyses were conducted to evaluate binding motifs at sites stereoselectively increased, decreased or unchanged by WX-02-23 relative to DMSO. Increased: log_2_(WX-02-23/DMSO) > 2 and log_2_(WX-02-43/DMSO) < 1, 681 sites. Decreased: log_2_(WX-02-23/DMSO) < -2 and log_2_(WX-02-43/DMSO) > -1, 1,482 sites. Unchanged: 1000 randomly picked sites unaffected by WX-02-23 or WX-02-43 (|Log_2_FC| < 1). *De novo* motif discovery was performed using MEME Suite software and the most significantly enriched sequence is depicted in the figure for each group. For a list of all significantly enriched sequences, see **Figures S6D**. **E.** *De novo* motif discovery analysis based on all ATAC-seq sites (left) and ATAC-seq at FOXA1 binding sites (right) in 22Rv1 cells. Groupings are made according to the ATAC-seq changes induced by WX-02-23 relative to DMSO. Increased: log_2_(FC) > 1 and *P* < 0.05*, N* = 903 for all ATAC-seq sites (left) or 542 at FOXA1 sites (right). Unchanged: |log_2_(FC)| < 1 and *P* > 0.05, *N* = 1000 (all ATAC-seq sites) or 600 (ATAC-seq at FOXA1 binding sites) picked at random. Decreased: log_2_(FC) < -1 and *P* < 0.05*, N* = 960 (all ATAC-seq sites) or 625 (ATAC-seq at FOXA1 binding sites). *De novo* motif discovery was performed using MEME Suite software and the most significantly enriched sequence is depicted in the figure for each group. For a list of all significantly enriched sequences, see **Figures S6E and F**. **F.** FOXA1-FKHD-DNA complex homology model from Figure 2A highlighting the proximal location of C258 near the ancillary 3-bp motif adjacent to the canonical core motif sequence. **G.** Frequencies of ancillary 3-bp motifs at FOXA1 binding sites. Sites are grouped by FOXA1 ChIP-seq changes with the same criteria used in Figure 6D. Frequency is defined as the proportion of individual ancillary 3-bp motifs among all 64 possible 3-bp motifs. Motifs are rank ordered from left to right based on the frequency of each motif in DMSO control treated cells. **H.** Correlation between frequencies of ancillary 3-bp motifs identified in FOXA1 ChIP-seq experiments (DMSO) and the estimated *K*_D_ values for FOXA1 binding to oligonucleotides containing these 3-bp motifs and a canonical core FOXA-binding motif from nanoBRET experiments (**see** Figure 3A) using NLuc-FOXA1. See **Figures S6D** for individual DNA binding curves. PCC = Pearson’s correlation coefficient. **I.** Effects of WX-02-23 on NLuc-FOXA1 interaction with oligo-1 variants with indicated ancillary 3-bp motifs and the core canonical sequence (GTAAACA) using NanoBRET assay described in Figure 3A (20 μM WX-02-23, 3 h) with indicated concentrations of TAMRA-oligonucleotides. Data for each oligo-1 variant are normalized to its DMSO control and are average values ± SEM from three independent experiments. Two-tailed *t*-test between DMSO and WX-02-23 *\*\*\*\*P <* 0.0001, \*\*\**P* < 0.001, *\*\*P <* 0.01. **J.** Additional NanoBRET experiment performed with oligo-1 variants having high affinity ancillary 3-bp motif sequences tested at 7.5 nM (near *K*_D_ values). Data for each oligo-1 variant are normalized to its DMSO control and are average values ± SEM from three independent experiments.

To further investigate the effect of altered FOXA1 binding on chromatin structure, we analyzed the fragment read length distribution of ATAC-seq reads, which can report on nucleosome packing and positioning^54^. Grouping FOXA1-binding sites again by the magnitude of change induced by WX-02-23 (substantially decreased, unchanged, and substantially increased) we analyzed the DNA fragment distribution for each group in cells treated with WX-02-23 or controls (DMSO, WX-02-43, and PladB). While no obvious difference in fragment read length distribution was observed across the control treatments, we found that sites decreased in FOXA1-binding by WX-02-23 had lower proportion of sub-nucleosomal fragments and higher proportion of mono-nucleosomal fragments (**Figures S6C**), indicating that nucleosomal DNA is invading sites formerly bound by FOXA1. Conversely, sites of increased FOXA1 binding exhibited the opposite shift with more subnucleosomal fragments and fewer mononucleosomal fragments, indicating chromatin decondensation at those sites (**Figures S6C**). Taken together, these data support that covalent modification of FOXA1_C258 by WX-02-23 can rapidly remodel the pioneer function of FOXA1, leading to the opening of sites on chromatin that were previously condensed, while also closing sites that were kept decompacted by this protein in its unliganded state.

In considering how WX-02-23 might alter FOXA1 interactions with chromatin, we performed *de novo* motif discovery analysis on FOXA1 ChIP-seq sites from cells treated with this compound. Consistent with previous studies^12,20,21,52,61^, unchanging sites displayed strong enrichment for the canonical core FOXA1 DNA binding motif (GTAAA[CT]A) along with less dramatic enrichment for a 5’-localized 3-base pair motif ([TA][AG][ATG]), referred to hereafter as the canonical ancillary 3-bp motif (**Figures 6D** and **S6D**). Interestingly, FOXA1-binding sites that were substantially and stereoselectively increased in WX-02-23-treated cells (WX-02-23 Log_2_FC > 2 and WX-02-43 |Log_2_FC| < 1) showed a clear diminishment in enrichment for the canonical ancillary 3-bp motif, in particular, in its two most 5’ positions (**Figure 6D**). Conversely, FOXA1-binding sites that were substantially and stereoselectively decreased in WX-02-23-treated cells (WX-02-23 Log_2_FC < -2 and WX-02-43 |Log_2_FC| < 1) displayed the opposite trend and displayed even stronger enrichment for the canonical ancillary 3-bp motif compared to unchanged sites (**Figure 6D**). Motif analysis using ATAC-seq data showed similar results, both when assessing WX-02-23-induced changes across all ATAC-seq sites or when first filtering ATAC-seq sites for those that overlap with FOXA1-binding sites (**Figure 6E** and **S6E**). Again, the canonical ancillary 3-bp motif was overrepresented in ATAC-seq sites that were diminished by WX-02-23 and underrepresented in ATAC-seq sites that were increased by WX-02-23. Remarkably, in the analysis of all ATAC-seq sites, the canonical FOXA1 binding motif was also by far the most dramatically enriched motif at sites altered by WX-02-23 (**Figures S6E**), supporting that the vast majority of chromatin accessibility changes induced by this compound occur through engagement of FOXA1.

A structural homology model of FOXA1 bound to DNA containing both canonical core and canonical ancillary motifs revealed that FOXA1_C258 is located in proximity to the ancillary 3-bp region (**Figure 6F**). Previous studies have also suggested that the FOXA Wing2 region makes contacts with the minor groove of this 3-bp stretch^13,41^. These findings, combined with our discovery that WX-02-23 induces FOXA1 binding to chromatin sites lacking the canonical ancillary 3-bp motif, led us to hypothesize that the covalent liganding of FOXA1_C258 alters DNA interactions with this specific region. We first calculated the frequency of appearance for each of the possible 64 sequences of the 3-bp region in the increasing, decreasing, or unchanged groups of FOXA1-binding sites from WX-02-23-treated cells. As expected from the *de novo* motif analysis, the unchanged and decreasing FOXA1-binding sites showed a marked enrichment for sequences matching the canonical ancillary motif of [TA][AG][ATG], and this motif was also enriched in DMSO-treated control cells (**Figures 6G** and **Supplementary Dataset S2**). In contrast the FOXA1 binding sites increased by WX-02-23, did not show this preference and instead exhibited greater relative enrichment for a spectrum of 3-bp sequences that were scarcely observed in the unchanged or decreasing groups (**Figures 6G**).

We next performed FOXA1 DNA-binding experiments using the NanoBRET assay (**Figure 3A**) with oligonucleotides containing canonical or non-canonical ancillary 3-bp sequences upstream of the canonical core FOXA1 motif. FOXA1 showed apparent *K*_D_ values for the canonical ancillary + canonical core motif oligonucleotides ranging from 6∼10 nM (**Figures S6F**), which were in close agreement with the values determined from previous studies using electrophoretic mobility shift assays^22,62^. On the other hand, FOXA1 displayed lower affinity for non-canonical ancillary + canonical core motif oligonucleotides (apparent *K*_D_ values ranging from 20∼65 nM; **Figures S6F**). We further observed a clear correlation between the relative affinity of FOXA1 for the tested oligonucleotides and their frequencies of occurrence at FOXA1 binding sites in DMSO-treated control 22Rv1 cells (PCC = -0.84) (**Figure 6H**). We interpret this result to indicate that, under basal conditions, the binding affinity displayed by FOXA1 for sequences in the ancillary 3-bp region influences the distribution of this protein across chromatin in cells.

Noting that our earlier *in vitro* DNA binding assays unintendedly used oligonucleotides harboring a canonical core FOXA1 motif and a non-canonical ancillary motif (ACA) (see **Figures 2J** and **3B**), we next considered whether WX-02-23 would similarly impact FOXA1 binding to DNA with a canonical core motif coupled to different ancillary 3-bp sequences. Interestingly, we found that WX-02-23 substantially increased the apparent affinity of FOXA1 for all of the non-canonical ancillary sequences tested, but generally did not alter FOXA1 interactions with canonical ancillary sequences across multiple test concentrations spanning the apparent *K*_D_ values for these oligonucleotides (**Figure 6I, J**). These results thus indicate that covalent liganding of FOXA1_C258 by WX-02-23 relaxes the DNA-binding preferences of this protein by increasing affinity for non-canonical sequences in the ancillary 3-bp region, thus providing a mechanistic rationale for the WX-02-23-induced remodeling of FOXA1-chromatin interactions in cells.

## Discussion

Due to their central roles in human physiology and disease, transcription factors (TFs) represent an exciting, yet still challenging protein class for chemical probe discovery and development^28,29,63^. The dearth of pharmacological tools for TFs has hindered their mechanistic investigations in living systems, as well as the advancement of small-molecule therapeutics targeting these proteins. Our findings highlight potential advantages of both chemical proteomics and covalent chemistry for TF ligand discovery and characterization. First, the chemical probes for FOXA1 were identified in ABPP experiments performed in living cells, which allowed for the discovery of a contextualized small molecule-protein interaction that we later found to reflect DNA-dependent binding. Alternative *in vitro* assays with purified FOXA1 protein may have missed such compounds. Second, homology models suggest that the loop in FOXA1 harboring C258 is dynamic, and we speculate that this region may only transiently adopt a structure compatible with small-molecule binding. Covalent chemistry may have a greater capacity to trap proteins, including TFs, in such rare, but kinetically accessible states^28,64^ Finally, the use of electrophilic stereoprobe libraries furnished a chemical probe (WX-02-23) paired with a physicochemically matched inactive control compound (enantiomer WX-02-43), an alkyne variant for measuring engagement of FOXA1 in cells (CJR-6A), and a probe-resistant mutant variant of FOXA1 (C258A mutant), each of which facilitated confident assignment of pharmacological effects as being on-target.

The integration of our biochemical and cellular data provide a model for WX-02-23 action where covalent engagement of FOXA1_C258 enhances DNA binding to sequences lacking the canonical ancillary 3-bp motif, leading to the redistribution of FOXA1 across the genome and the binding to and opening of previously unoccupied and inaccessible sites bearing these suboptimal motifs. Although the canonical ancillary 3-bp motif has been described in several previous studies assessing the DNA-binding preferences of FOXA1^12,20,52,61^, its functional relevance has not, to our knowledge, been investigated. Our findings thus point to an important and heretofore underappreciated role for this 3-bp motif in contributing to FOXA1-chromatin interactions, and one that is specifically sensitive to modification of FOXA1 by covalent ligands targeting C258. Given (i) the intrinsic effects of the 3-bp ancillary motif on FOXA1-DNA binding affinity, (ii) the predicted proximity of this ancillary motif to C258, iii) the DNA-dependency of stereoprobe binding to FOXA1_C258; and (iv) the altered ancillary sequence preferences exhibited by WX-02-23-bound FOXA1, we speculate that covalent liganding of C258 directly modulates the interactions of FOXA1 with the 3-bp ancillary motif. Structural studies of WX-02-23-modified FOXA1 bound to oligonucleotides displaying canonical or non-canonical ancillary 3-bp motifs may provide further insights into how covalent modification of C258 alters FOXA1-DNA interactions.

The correlated changes in FOXA1 localization and chromatin accessibility induced by WX-02-23 are broadly consistent with past studies establishing a pioneer function for FOXA1^3^ and further support the dynamic mobility of this TF on chromatin as shown in previous single molecule studies with ectopically expressed FOXA1^42,43,50,51^ and here demonstrated pharmacologically with endogenous FOXA1. Indeed, the acute nature of chemical biology studies allowed us to investigate the pioneering capabilities of endogenous FOXA1 while avoiding the potential limitations, such as changes in cell growth or cell state, imposed by prolonged genetic manipulations of this essential lineage TF^42^. In considering how the chemical redirection of FOXA1 to compacted chromatin regions triggers remodeling to achieve an open chromatin state, we suspect, as others have suggested^23,56,65^, that FOXA1 binding likely paves a path for ATP-dependent chromatin remodelers to promote the stable formation of open chromatin. Consistent with this hypothesis, we found by co-IP experiments that FOXA1 associates with chromatin remodeling such as SWI-SNF and these interactions were unperturbed by WX-02-23 engagement.

We anticipate that additional mechanistic insights can be gained in future studies that evaluate the effects of WX-02-23 on FOXA1-chromatin interactions across more time points beyond the 3 h treatment evaluated herein. For instance, we observed that ∼10% of FOXA1 binding sites were substantially changed by WX-02-23 (**Figure 4B**), and we wonder whether even more extensive changes in FOXA1-chromatin interactions may occur following longer treatment times with WX-02-23. Conversely, it would be interesting to understand how soon after WX-02-23 engagement FOXA1 begins to relocalize across the genome. Such shorter time-point analyses would need to take into consideration the time required for WX-02-23 to covalently engage FOXA1 in cells.

While the WX-02-23-induced movement of FOXA1 to new genomic binding sites harboring noncanonical ancillary motifs could be rationalized by our *in vitro* DNA-binding assays, it is less clear why WX-02-23 also depletes FOXA1 from a small percentage of sites harboring canonical motifs. One possibility is that FOXA1 is decreased at these sites as a secondary effect of its enhanced binding to sites with sub-optimal motifs. However, we might have alternatively expected the enrichment of WX-02-23-engaged FOXA1 at *de novo* binding sites to be accompanied by partial losses of FOXA1 at all canonical sites; instead, we observed large losses of FOXA1 from a relatively small number of sites. It is interesting to consider what factors might drive this outcome. Lost interactions could reflect highly dynamic sites of FOXA1 binding with shorter residence times. Other TFs that bind cooperatively with FOXA1 might also contribute to this phenomenon^51,61^, or it could involve local aspects of the chromatin environment, including the number of canonical FOXA1 binding motifs, histone modifications, and higher-order chromatin organization. Alternatively, it is also possible that WX-02-23 acts as a direct antagonist of DNA binding at these sites. We did not observe evidence for such site-specific antagonism *in vitro* (i.e., FOXA1 binding to motifs with canonical 3-bp ancillary sequences were unaffected by WX-02-23 *in vitro*), but we cannot rule out its occurrence in a cellular environment.

Finally, when contemplating the potential translational implications of our findings, there is a wealth of genetic evidence that FOXA1, in its wild type form, serves as an essential lineage factor for breast and prostate cancer cell growth^66,67^. Small-molecule antagonists of FOXA1 thus could represent a compelling class of future cancer therapeutics. Our functional studies indicate that WX-02-23 does not act as a simple antagonist of FOXA1, but rather remodels the pioneer function of this protein. Indeed, the rapid redistribution of FOXA1 across chromatin caused by WX-02-23 is reminiscent of the enhanced mobility observed for cancer-related Wing2 domain mutants of FOXA1^21^, raising the provocative possibility that WX-02-23 may act in an agonistic manner. We also note, however, that alternative base editing-induced mutations of the local sequence encoding Wing2 residues around and including C258 have been found to reduce growth rates of prostate cancer cells^68^, suggesting that analogues of WX-02-23 or structurally distinct chemistries binding the C258 pocket in FOXA1 may exert inhibitory activity by, for instance, blocking FOXA1 binding to DNA. Additionally, one could imagine converting WX-02-23 into heterobifuctional compounds that direct FOXA1 to E3 ligases for degradation^69^ or to additional transcriptional regulators to corrupt the pro-tumorigenic gene expression profiles of FOXA1-dependent cancer cells^70^. It will also be interesting to determine how the cancer-related mutations in the Wing2 domain impact WX-02-23 binding to FOXA1 and, conversely, whether allele-specific covalent ligands can be generated for the subset of FOXA1 mutants that retain C258.

In summary, we have demonstrated that small molecules covalently targeting the DNA-binding domain of the pioneer TF FOXA1 can rapidly remodel its genomic localization to produce correlated changes in chromatin accessibility. These findings provide further evidence to support a striking plasticity underpinning FOXA1’s interactions with chromatin and, more generally, underscore the value of chemical probes as tool to study and remodel the functions of pioneer TFs in their endogenous states in cancer cells. We envision that continued efforts to leverage covalent chemistry and chemical proteomics will enabled the discovery of additional first-in-class ligands for historically challenging proteins like TFs and the broader class of DNA-binding proteins.

### Limitations of the study

WX-02-23, as an early stage tool compound, stereoselectively cross-reacted with a handful of other proteins, one of which – the spliceosome factor SF3B1 – was of particular concern, as we have shown previously this interaction has profound transcriptomic and cell growth effects^37^. Fortunately, we were able to use a chemically unrelated natural product modulator of SF3B1, pladienolide B, as an additional control compound to facilitate the assignment of FOXA1-dependent effects of WX-02-23 in prostate cancer cells. Even with this control, however, some cell biology experiments, such as assessing potential gene expression changes caused by WX-02-23-mediated remodeling of FOXA1-chromatin interactions, were not possible due to SF3B1 cross-reactivity, and therefore an important future objective will be to improve the specificity of stereoprobe ligands for FOXA1. It may also be possible to engineer cancer cell lines where the *SF3B1* gene is edited to express a WX-02-23-resistant mutant (e.g., C1111A-SF3B1), which would provide another way to study the effects of WX-02-23 on FOXA1-related biology without potential confounding impact on spliceosome function.

## Methods

### Cell lines

22Rv1, Ramos, HEK293T, LentiX 293T cells were grown in RPMI 1640 (22Rv1, Ramos) or DMEM (HEK293T, LentiX 293T) supplemented with 10% fetal bovine serum (FBS), 2 mM L-alanyl-L-glutamine (GlutaMAX), penicillin (100 U/ml), and streptomycin (100 μg/ml) and maintained at 37 °C with 5% CO_2_. Expi293F cells were grown in Expi293 Expression Medium (Thermo; A1435101) and maintained at 37 °C, 8% CO_2_ and constant shaking at 120 rpm. All cell lines were routinely inspected for mycoplasma contamination.

### Generation of 22Rv1 Y36C-PHF5A cells with inducible FOXA1 expression

LentiX 293T cells were seeded and once it reached 70% confluency on a 10 cm dish, a mixture of 3.4 μg pINDUCER21_2xStrep-Flag-FOXA1-HA (or C258A), 0.85 μg pMD2.G (Addgene #12259), 1.7 μg psPAX2 (Addgene #12260), 48.8 μg PEI (1 μg/μL) in 200 μL OPTI-MEM were added for lentiviral particle production. After 60 hours, the virus containing media was collected and 250 μL was added to 1 million 22Rv1 cells stably expressing Y36C-PHF5A, which was generated from previous study^37^, in the presence of 8 μg/mL polybrene (3 mL total in 6-well dish). Cells were spin-infected for 45 minutes at 2000 rpm. Cells were expanded to 15 cm dish and sorted for GFP expression by flow cytometry to enrich for transduced cells. The cells were treated with 0.5 μg/mL doxycycline for 24 hours prior to ligand treatments.

### Chemical synthesis of stereoprobes

Synthetic procedures and characterization of CJR-6A and CJR-6B are described in Methods S1. The synthesis and characterization of WX-02-13, WX-02-23, WX-02-33, WX-02-43, WX-01-10, and WX-01-12 have been reported in previous studies^37,71^.

### Cysteine-directed activity based-protein profiling

Cysteine-directed ABPP was performed as previously described^38^. Briefly, 22Rv1 cells (20 million cells in 15 cm dish) or Ramos cells (12.5 mL of 3 million cells/mL) were treated with DMSO or stereoprobes (WX-02-13, WX-02-23, WX-02-33, WX-02-43; 20 μM; 3 h). Cells were washed with ice-cold DPBS (3x), followed by resuspension in DPBS and lysed by probe-sonication (2 x 15 pulses; 10% power output). Proteome was normalized to 2 mg/mL in 500 μL and treated with 0.1 mM iodoacetamide-desthiobiotin (IA-DTB) for 1 hour at room temperature. Proteins were precipitated with ice-cold solvents (600 μL MeOH, 200μL chloroform, and 100 μL water). After vigorous vortexing and centrifugation (16000 x g, 10 min), solvents were aspirated. The protein-disk was washed with 1 mL MeOH and resuspended in 90 μL of reducing buffer (9 M urea, 10 mM DTT, 50 mM triethylammonium bicarbonate (TEAB), pH 8.5). Samples were heated (65 °C, 20 min), followed by incubation with 50 mM iodoacetamide (30 min, 37 °C in dark). Samples were diluted to 2 M urea by adding 50 mM TEAB. Mass-spectrometry grade trypsin (Promega; 1 μg in trypsin resuspension buffer with 25 mM CaCl_2_) was added to each sample and digested overnight at 37 °C. Samples were then incubated with streptavidin-agarose beads (50 μL of slurry resuspended in 300 μL wash buffer (50 mM TEAB, 150 mM NaCl, 0.2% NP-40)) for 2 hours at room temperature. Beads were transferred to BioSpin (Bio-Rad) columns to remove solution and washed (3x 1mL wash buffer, 3x 1 mL DPBS, 3x 1 mL water). Enriched peptides were eluted by elution buffer (300 μL 50% acetonitrile with 0.1 % formic acid) and speedVac to dryness. Peptides were resuspended in 200 mM EPPS (pH 8.0) and Tandem Mass Tag (TMT; 10-plex) labeled (90 min, room temp). The TMT-labeled peptides were de-salted by Sep-Pak C18 cartridges (Waters), followed by fractionation using high-pH high performance liquid chromatography (HPLC). A total of 12 fractions were analyzed by Orbitrap Fusion Mass Spectrometer with Xcalibur v.4.3 (see below for LC-MS instrumentation and analysis).

### Protein-directed activity based-protein profiling

Protein-directed ABPP was performed as previously described^38^. Briefly, 22Rv1 cells were treated with stereoprobes (DMSO, WX-02-23, or WX-02-43; 20 μM; 2h), followed by the alkyne probes (CJR-6A or CJR-6B; 5 μM; 1h). Cells were washed with ice-cold DPBS (3x). Cell pellets were resuspended in DPBS and lysed by probe-sonication (2x15 pulses; 10% power output). Proteome was normalized to 2 mg/ml in 500 μL, and the alkyne probe labeled proteins were conjugated to biotin-PEG4-azide via Cu(I)-catalyzed azide–alkyne cycloaddition (CuAAC; 2 mM CuSO_4_, 100 μM Biotin-PEG4-azide, 1 mM tris(2-carboxyethyl)phosphine (TCEP), 100 μM tris((1-benzyl-4-triazolyl)methyl)amine (TBTA); 1 h. After chloroform/methanol precipitation followed by centrifugation (16,000 g, 10 min, 4 °C), solvents were aspirated and the protein-disk was washed with 1 mL ice-cold MeOH and resuspended in 0.5 mL resuspension buffer (8 M urea, 0.2 % SDS in DPBS). Peptides were reduced (10 mM DTT; 15 min; 65 °C) and alkylated (20 mM iodoacetamide; 30 min; 37 °C). After adding 130 μL 10% SDS, samples were diluted to 6 mL in DPBS and incubated with 100 μL of streptavidin-agarose bead slurry for 90 minutes. Beads were washed (2 x 0.2 % SDS in DPBS, 1 x DPBS, 2x H_2_O, 1 x 200 mM EPPS, pH 8.0) and enriched proteins were on-bead digested with mass-spectrometry grade trypsin in buffer (2 M urea in 200 mM EPPS, 1 mM CaCl_2_) overnight at 37 °C. Digested peptides were TMT-labeled (16-plex) for 90 minutes and quenched by hydroxylamine. The TMT-labeled peptides were desalted by Sep-Pak C18 cartridges and high-pH fractionated using peptide desalting spin columns (Thermo 89852). A total of 10 fractions were analyzed by Orbitrap Fusion Mass Spectrometer with Xcalibur v.4.3 (see below for LC-MS instrumentation and analysis).

### FOXA1 immunoprecipitation-mass-spectrometry (IP-MS)

15 million 22Rv1 Y36C-PHF5A cells with or without (mock) doxycycline-inducible expression of 2xStrep-Flag epitope-tagged FOXA1 were treated with 0.5 μg/ml doxycycline for 24 hours, followed by DMSO, stereoprobes (WX-02-23 or WX-02-43; 20 μM) or PladB (100 nM) for 3 hours. Cells were washed (3 x) with ice-cold DPBS and nuclei fraction was isolated with NE-PER Nuclear and Cytoplasmic Extraction kit (Thermo) supplemented with 1 x Halt Protease Inhibitor Cocktail (Thermo). Instead of using nuclear extraction reagent (NER), crude nuclei fraction was resuspended in IP-lysis buffer (Thermo; 25 mM Tris pH 7.4, 150 mM NaCl, 1 % NP-40, 1 mM EDTA, 5 % glycerol) supplemented with 1 x Halt Protease Inhibitor Cocktail and probe-sonicated (15 x 10% power output). 2 mg of nuclear proteome in 500 μL was incubated with 50 μL of anti-Flag M2 magnetic beads (Sigma) for 3 hours at 4 °C. The beads were quickly washed with ice-cold IP-lysis buffer (3 x) and then DPBS (1 x) on a magnetic stand. Enriched proteins were eluted with 8M urea in DPBS (65 °C; 10 min), reduced (10 mM DTT; 65 °C; 15 min), and alkylated (25 mM iodoacetamide; 37 °C; 30 min). Samples were diluted to 2 M urea by adding 200 mM EPPS (pH 8.0) and trypsin (2 μg) digested overnight at 37 °C. Digested peptides were TMT-labeled and quenched by hydroxylamine. The TMT-labeled peptides were desalted by Sep-Pak C18 cartridges and high-pH fractionated using peptide desalting spin columns (Thermo 89852). A total of 3 fractions were analyzed by Orbitrap Eclipse Mass Spectrometer with Xcalibur v.4.3 (see below for LC-MS instrumentation and analysis).

### TMT liquid chromatography-mass-spectrometry (LC-MS) analysis

All samples in this study used the same instruments and workflow settings as described previously^37,38^. Briefly, all samples were analyzed by Orbitrap Fusion Tribrid Mass Spectrometer (Thermo Scientific; all cysteine- and protein-directed ABPP studies) or Orbitrap Eclipse Tribrid Mass Spectrometer (FOXA1 IP-MS studies) coupled to an UltiMate 3000 Series Rapid Separation LC system and autosampler (Thermo Scientific Dionex). Data was acquired using an MS3-based TMT method (see ref.^37^ for the detailed settings) and the RAW files were uploaded and processed by Integrated Proteomics Pipeline (IP2, v.6.7.1), and searched using the ProLuCID algorithm^72^ using a reverse concatenated, non-redundant variant of the Human UniProt database (release 2016). Cysteine residues were searched with a static modification for carboxyamidomethylation (+57.02146 Da) and for cysteine-directed ABPP, a dynamic modification for IA-DTB labeling (+398.25292 Da) was included for a maximum number of 2 differential modifications. N-term and lysine static modification from the tags (+229.1629 Da for 10-plex in Cys-directed ABPP, +304.2071 Da for 16-plex in protein-directed ABPP and FOXA1 IP-MS) were included. Peptides were required to be at least 6 amino acids long and filtered to through DTASelect (version 2.0) to obtain a peptide false-positive rate below 1%. The MS3-based peptide quantification was performed with reporter ion mass tolerance set to 20 ppm with Integrated Proteomics Pipeline (IP2).

#### -Data processing of cysteine-directed ABPP

The census output files from Integrated Proteomics Pipeline 2 (IP2, v.6.7.1) were further processed to calculate cysteine engagement ratios (probe vs DMSO) by dividing each TMT reporter ion intensity by the average intensity for the DMSO channels. Peptide-spectra matches were grouped based on protein ID and the cysteine residue number. Peptides with summed reporter ion intensities < 10000, coefficient of variation for DMSO channels > 0.5 were excluded from analysis. TMT ion intensities were median normalized per TMT channel. Two independent replicates of 10-plex experiments in 22Rv1 cells were analyzed together (total n = 2-4 per treatment condition). One 10-plex experiment in Ramos cells (n = 2 per treatment condition) was analyzed separately. Data were averaged from all replicates. A cysteine site was considered stereoselectively liganded if the variability corresponding to the probe leading to highest blockade of iodoacetamide-desthiobiotin (IA-DTB) did not exceed 20%, and the following additional criteria were met: (i) the average IA-DTB blockade by the probe was > 66.7% and > 2-fold that of its enantiomer; (ii) the same probe led to < 25% IA-DTB blockade of at least one other cysteine in the same protein; (iii) the corresponding protein did not show uniform changes in peptide enrichment by IA-DTB, indicative of protein abundance changes rather than legitimate liganding of a given site. Specifically, if a protein had multiple cysteines quantified and verified both average IA-DTB blockade > 25% and SEM < 15%, it was subject to manual inspection to determine whether a uniform change in reactivity was observed across all quantified cysteine-containing peptides.

#### -Data processing of protein-directed ABPP

The census output files from IP2 were further processed to calculate enrichment engagement ratios (probe vs probe) by dividing each TMT reporter ion intensity by the sum of intensity for all the channels. Each spectrum-peptide match was grouped based on protein ID, excluding peptides with summed reporter ion intensities < 10,000, coefficient of variation of > 0.5, < 2 unique peptides per protein ID. TMT reporter ion intensities were normalized to the median summed signal intensity across channels. A protein was considered stereoselectively liganded by WX-02-23 if the following criteria were met: (i) > 4-fold higher enrichment by CJR-6A vs CJR-6B; (ii) > 2-fold higher residual enrichment by CJR-6A following blockade by WX-02-43 vs WX-02-23; (iii) > 66.7% blockade of CJR-6A by WX-02-23.

#### -Data processing of FOXA1 IP-MS

The census output files from IP2 were further processed to compute enrichment engagement ratios (probe vs probe) by dividing each TMT reporter ion intensity by the sum of intensity for all the channels. Each spectrum-peptide match was grouped based on protein ID, excluding peptides with summed reporter ion intensities < 10,000, coefficient of variation of > 0.5. Peptide hosting FOXA1_C258 was removed to account for FOXA1_C258A mutation and loss of signal in the WT samples due to modification by WX-02-23. 3 independent replicates were pooled and averaged. Enriched proteins were defined as proteins that were substantially ((WT-FOXA1/mock) >2) and significantly (*P* < 0.05) enriched in ≥ 2 independent datasets (out of 3), ≥ 2 unique peptides quantified in at least 1 dataset, ≥ 2 average spectral counts.

### Plasmid construction

To generate plasmids for recombinant protein expression, cDNA of hFOXA1 (Gene ID: 3169), hFOXA2 (Gene ID: 3170) and hFOXA3 (Gene ID: 3171) were PCR amplified with indicated primers (see Table S1) and inserted into backbone plasmids via restriction digest and ligation (all enzymes from NEB) or Gateway cloning (Thermo). All mutangenesis were performed by Q5 Site-Directed Mutagenesis Kit (NEB; E0554S). For pCAG-OSF-FOXA1 (N-terminus 2xStrep-Flag-tagged; see **Figure 1**), pCAG-OSF-VPS34 (addgene # 99327) plasmid was digested with Kpn1 + Xho1 to remove VPS34 and insert FOXA1. For MBP-tagged FOXA expressions (see **Figure 2**), pHLmMBP-8 (addgene #72347) was digested with indicated restriction enzymes (see Table S1) and FOXA or FOXA1-FKHD cDNA was ligated-in. To generate plasmids for the Luciferase-tagged proteins (see **Figure 3**), pMCS-Green Renilla Luc vector (Thermo; 16152) was used to obtain the Luciferase cDNA and first inserted into pCAG-OSF-VPS34 digested with Kpn1 + Xho1, and FOXA1 was inserted after digestion with Xho1 and Bsu36I. For NanoBRET plasmids (see **Figure 3**), 2xStrep-Flag tag was inserted to pHLmMBP-8 after digestion with EcoR1 + Not1 (removing the MBP-tag), then NanoLuc cDNA from pNL1.1.CMV[Nluc/CMV] (Promega; N1091) was inserted at Not1 + Sac1 (NLuc-FOXA1-FKHD) or MluI + Bsu36I (full-length FOXA1-NLuc) restriction digest sites. FOXA1 construct of interest was inserted after digestions with Sac1 + Xho1 (N-term NanoLuc) and Xho1 + MluI (C-term NanoLuc). For doxycycline inducible expression of FOXA1 (see **Figure 5**), 2xStrep-Flag-FOXA1-HA was introduced to pINDUCER21 (Addgene #46948) to generate pINDUCER21_FOXA1 by Gateway cloning protocol (Thermo).

### Transient transfection of epitope-tagged proteins of interest

HEK293T cells were seeded in 6-well dishes and 48 hours later (70-80% confluent) transfected with gene of interest cloned into expression vectors by incubating 2 μg plasmid and 6 μg PEI (1 μg/μL) in 200 μL OPTI-MEM (Thermo; 31985062) for 20 minutes. The mixture was added drop-wise into refreshed media. Experiments were performed 24-48 hours later. For protein purification purposes, manufacturer’s guidelines for ExpiFectamine™ 293 Transfection Kit (Thermo; A14524) was used with modifications. Expi293F cells were diluted to 2.5 million cells/mL with fresh media. Per 100 mL cell culture transfection, 100 μg of plasmid (MBP-FOXA1-FKHD) was diluted in 5 mL OPTI-MEM and added to 5 mL OPTI-MEM containing 270 μL ExpiFectamine 293 reagent and incubated for 15 minutes. The mixture was added to cells and after 22 hours, a mixture of Expifectamine 293 Transfection Enhancer 1 (500 μL) and Enhancer 2 (5 mL) was added. Cells were collected 60 hours later and frozen for protein purification.

### Gel-based activity based-protein profiling

HEK293T cells expressing recombinant protein of interest were treated with steroprobes as indicated (0.1 % DMSO final in media) or collected and lysed in DPBS for *in vitro* treatments. Cell lysates were normalized to 1 mg/ml in 44 μL and CuAAC reaction was performed by adding 6 μL of click-master mix (1 μL 1.25 mM TAMRA-azide, 1 μL 50 mM CuSO4, 3 μL 1.7 mM TBTA in 1:4 DMSO: t-butanol, 1 μL 50 mM TCEP) and incubated for 1 hour at room temperature. Samples were resolved on SDS-PAGE and in-gel fluorescence was acquired by Bio-Rad Chemidoc imager. ImageJ software was used for fluorescent band intensity quantification.

### Immunoblot analysis

Proteins were resolved by SDS-PAGE and transferred to either nitrocellulose membrane using a power blotter semidry transfer system (Thermo) or PVDF membrane (60 V; 2 h in Towbin buffer 20% methanol). The membrane was blocked with 5% non-fat milk in tris-buffered saline with Tween (TBST) buffer (20 mM Tris-HCl pH 7.5, 150 mM NaCl, 0.1% tween 20) for 1 h at room temperature. All blots were incubated with primary antibody (1:1000) overnight at 4 °C, followed by appropriate HRP-conjugated secondary antibody (1:1000) at room temperature (1h). After several washes, 1:1 mix of SuperSignal West Pico PLUS stable peroxide and luminol/enhancer (Thermo) was used to initiate chemiluminescence. All images were acquired by Bio-Rad Chemidoc imager.

### Generation of the 3-D structure of FOXA1-FKHD homology model

The 3-D model of FOXA1-FKHD (AA168-268) was generated by SWISS-MODEL^73^ using FOXA3-FKHD-DNA complex (PDB: 1VTN; resolution 2.5 Å)^41^ as a template, which has 95 % identity and has FOXA1_C258 equivalent cysteine resolved. Since FOXA3-FKHD-DNA complex did not use canonical motif DNA, to build a DNA model in our structure, we aligned the FOXA1-FKHD homology model with FOXA2-FKHD-DNA complex (PDB: 5X07)^40^, which used 16-bp DNA containing canonical motif, including optimal ancillary 3-bp (AATGTAAACA).

### Protein purification

Frozen Expi293F cells transfected with MBP-FOXA1-FKHD were thawed in lysis buffer (HEPES pH7.5, 300mM NaCl, 10 mM MgCl2, 1% glycerol, 0.5 mM TCEP, 1 x Halt protease inhibitor cocktail) and lysed by probe sonication. Lysate was cleared by centrifugation at 20000 x g for 45 minutes at 4 °C. Supernatants were incubated with pre-washed Amylose resin (NEB) and incubated with gentle rotation for 2 hours at 4 °C. Resin was washed thoroughly with wash buffer (10 mM HEPES pH7.5, 300 mM NaCl, 10 mM MgCl_2_, 0.5 mM TCEP) and eluted with elution buffer (10 mM HEPES pH7.5, 150 mM NaCl, 10 mM MgCl_2_, 0.5 mM TCEP, 10mM maltose). Eluents were further purified by fast protein liquid chromatography (FPLC; Superdex200 column; wash buffer constant gradient), concentrated and stored in wash buffer supplemented with 10 % glycerol.

### DNA-dependent stereoprobe labeling

HEK293T cells recombinantly expressing FOXA1 or FOXA1-FKHD (AA168-274) were collected and resuspended thoroughly with 0.1 % NP-40 in DPBS, followed by centrifugation at 10000 x g (5min at 4 °C). After saving supernatant (cytosol), the pellet was washed again with 0.1 % NP-40 in DPBS and centrifuged again to remove supernatant. The nuclei fraction was then resuspended in DPBS and probe sonicated (20 pulses). Cell fractions were normalized to 1 mg/ml in 43 μL and treated with 1 μL of benzonase (Sigma E1014; 30 min, 37 °C water-bath). Cell lysate was then incubated with CJR-6A or -6B (10 μM; 1h) and processed for gel-ABBP as described above. In case of purified proteins, MBP-FKHD (500 nM) was incubated with indicated concentration of DNA oligonucleotide duplex (annealed by mixing equal moles of matching ssDNA, heated in 95 C heat block for 2 minutes and then gradually cooled to room temperature) for 30 minutes in DPBS supplemented with 2 mM MgCl_2_, followed by addition of stereoprobes. Samples were processed for gel-ABPP as described above.

### Luciferase (Luc)-based immobilized oligonucleotide assay

HEK293T cells expressing recombinant Green Renilla Luc-FOXA1 or Luc-FOXA1-FKHD were treated with stereoprobes (48 hr post transfection; 20 μM; 3h). Cells were washed with DPBS (3 x) and after collection, nuclei extract was isolated by NE-PER kit (Thermo) supplemented with 1 x Halt protease inhibitor cocktail (Thermo) according to manufacturer’s guidelines. Pierce Streptavidin coated white 96-well plates (Thermo 15118) were blocked with blocking buffer (10 mM Tris pH7.5, 50 mM KCl, 1.25 % glycerol, 0.025 % NP-40, 5 mM MgCl_2_, 0.1 % BSA w/v) for 10 minutes and washed with the same buffer without BSA (wash buffer). Biotinylated oligonucleotide duplexes (3 μM in 100 uL wash buffer) was added to each well and incubated for 1 hour at room temperature. After 3 washes, nuclear lysates (free of chromatin; 0.1 mg in 100 μL) and 1 μg poly(dI-dC) (Thermo) were incubated on plate with constant shaking (200 rpm; room temp; 1h). After 3 washes in wash buffer (5 min intervals with shaking), 1x coelenterazine (Thermo 1862430) in DPBS was added and immediately measured luminescence on microplate reader (Clariostar; em 480-80 nm). Signals were normalized to appropriate controls in each experiment (eg. DMSO).

### NanoBRET-based oligonucleotide binding assay

Overall procedure was adapted from a previous study^48^. HEK293T cells expressing recombinant NLuc-tagged FOXA1 or FOXA1-FKHD were treated with stereoprobes (24 hr post transfection; 20 μM; 3h). Cells were washed with DPBS (3 x) and lysed by probe sonication (20 pulses) in DPBS supplemented with 1 x Halt protease inhibitor cocktail. Cell lysates were normalized to 0.2 mg/ml in DPBS supplemented with 1 x Halt protease inhibitor cocktail and indicated concentrations of TAMRA-tagged oligonucleotide duplexes were incubated for 20 minutes. After transferring mixtures to white 384-well microplate, 3 μL of 1:100 diluted NanoBRET Nano-Glo substrate (Promega N1571) was added, and NanoBRET signal was measured immediately on microplate reader (Clariostar). Multichromatic settings with appropriate filters were used to measure emission from both Acceptor (TAMRA-DNA; >610 nm (longpass filter)) and Donor (NLuc; 460±80 nm (bandpass filter)). Each condition included a control blank sample without Acceptor (no TAMRA-DNA) to subtract any background bleed-through signal. BRET = ((Acceptor_sample_/Donor_sample_) – (Acceptor_control_/Donor_control_)). Data were normalized to relevant control (eg. DMSO samples).

### Gene ontology analysis

A list of significantly and substantially enriched proteins in FOXA1-IPMS (WT: DMSO/mock) was analyzed using Gene Ontology Enrichment Analysis and Visualization Tool (GOrilla)^74^. *P*-value threshold of 10^-5^ was used for gene ontology enrichment of biological processes and the output was visualized by REVIGO^75^ (version 1.8.1) to remove redundant terms.

### ChIP-seq

Parental 22Rv1 or 22Rv1cells recombinantly expressing Y36C-PHF5A and FOXA1 (WT or C258A; induced by 0.5 μg/mL doxycycline for 24 hours prior to the experiment) were treated with stereoprobes (WX-02-23, WX-02-43; 20μM) or PladB (10 nM for parental cells and 100 nM for recombinant Y36C-PHF5A expressing cells) for 3 hours (15 cm dish; 70-80% confluency). In case of FOXA1 knock-down, cells were transfected with 225 pmol control siRNA (ThermoFisher Invitrogen Silencer Select, 4390843) or FOXA1 siRNA (Thermo Fisher Ambion FOXA1 Silencer Select, s6688) using Lipfectamine 2000 for 24 hours prior to DMSO treatment. Cells were washed, trypsinized, quenched and fixed with formaldehyde buffer (1.1 % formaldehyde final in 50 mM HEPES pH 7.5, 100 mM NaCl, 1 mM EDTA, 0.5 mM EGTA) for 10 minutes. The crosslinking reaction was quenched with 125 mM glycine. Cells were washed with ice-cold DPBS (3x) and frozen at -80 °C for processing. Cells were resuspended in lysis buffer 1 (LB1; 50 mM HEPES, 140 mM NaCl, 1 mM EDTA, 10% glycerol, 0.5 % NP-40, 0.25 % triton x-100, 1 x Halt Protease Inhibitor Cocktail) and incubated for 10 minutes at 4 °C with gentle rotation. After removing supernatant, cells were washed again in lysis buffer 2 (LB2; 10 mM Tris pH 7.5, 200 mM NaCl, 1 mM EDTA, 0.5 mM EGTA, 1 x Halt Protease Inhibitor Cocktail). The isolated nuclei were resuspended in sonication buffer (50 mM HEPES pH 7.5, 140 mM NaCl, 1 mM EDTA, 1 mM EGTA, 1 % Triton x-100, 0.1 % Na-deoxycholate, 0.5 % SDS, 1 x Halt Protease Inhibitor Cocktail) and water-bath sonicated (Diagenode; 25 cycles, 30 sec on and off). Sheard chromatin was diluted to achieve 0.1 % SDS and incubated with FOXA1 antibody (Novus Biologicals, NBP2-45354) pre-conjugated on Dynabeads (Thermo, Protein G, 10004D) overnight at 4 °C. Beads were washed with sonication buffer (2x, without SDS), the sonication buffer with higher salt (1x, 500 mM NaCl, without SDS), LiCl wash buffer (1x, 20 mM Tris pH7.5, 1 mM EDTA, 250 mM LiCl, 0.5 % NP-40, 0.5 % Na-deoxycholate), and TE supplemented with 50 mM NaCl. Sheared chromatin was eluted from the beads by incubating with elution buffer (50 mM Tris pH7.5, 10 mM EDTA, 1% SDS; 65 °C for 15 min) and reverse crosslinked at 65 °C for 13 hrs. Samples were treated with RNaseA (37 C, 2h), ProteinaseK (55 C, 30 min), and phenol:chloroform:isoamyl alcocol (25:24:1). The top layer was isolated, followed by ethanol precipitation (-80 C for 2-3 h) to isolate DNA fragments ranging 300-500bp. Samples were processed for DNA library preparation (ThruPLEX DNA-seq Kit, Takara Bio), cleaned up with AMpure XP beads (Beckman Coulter), and sequenced on Illumina Nextseq 2000 (single-end 75-bp).

### ChIP-seq analysis

Sequencing reads were aligned to human genome build hg19 and RefSeq genes by Bowtie2 with the following parameters: -N 1 -p 10 -x. Using the unenriched input sample as background, the signal-enriched regions were identified by Model-based Analysis of ChIP-seq (MACS) peak-finding algorithm (v1.4.1) with the *P* value threshold of 1e-9. A merged peak list including all FOXA1-binding sites from all relevant conditions (DMSO, WX-02-23, and WX-02-43 for ChIP-seq in parental 22Rv1 cells) was created with bedtools (v2.17.0) for downstream analyses. Using the read density calculator Bamliquidator (v1.0) (http://github.com/BradnerLab/pipeline/wiki/bamliquidator) to map sequencing reads to designated loci with a 200 bp extension window in either direction differential analyses were performed across samples. Samples were normalized by total number of mapped reads (reads per million, rpm) and read density per base pair was calculated (rpm/bp). Sites with high background signal (input sample > 0.5 rpm/bp) were excluded. For the differential analyses, an arbitrary threshold (1 rpm/bp for ChIP-seq with parental 22Rv1 and 0.5 rpm/bp for ChIP-seq with engineered 22Rv1) was used to filter out the loci with low signals in all treatments to minimize the drastic fold changes induced by random noise. To select for the sites with WX-02-23-induced-stereoselective changes for analyses in **Figures S4C, 6D,** and **6G** the filters as following are used: stereoselectively increased, log_2_(WX-02-23/DMSO) >2 and Log_2_(WX-02-43/DMSO)<1; unchanged, -1<Log_2_(WX-02-23/DMSO)<1 and -1<Log_2_(WX-02-43/DMSO)<1); stereoselectively decreased, log_2_(WX-02-23/DMSO) <-2 and Log_2_(WX-02-43/DMSO)>-1. All box plots represent 25-75 percentile with whiskers extending 1.5 interquartile range (IQR) and the center line represents the median.

### ATAC-seq

Parental 22Rv1 or 22Rv1 cells recombinantly expressing Y36C-PHF5A and FOXA1 (WT or C258A; induced by 0.5 μg/mL doxycycline for 24 hours prior to the experiment) were treated with stereoprobes (WX-02-23, WX-02-43; 20μM) or PladB (10 nM and 100 nM for parental 22Rv1 cells, and 100 nM for engineered 22Rv1 cells) for 3 hours (6-well dish; 70-80% confluency). Cells were washed with DPBS (3x) and trypsinized to detach from the surface and washed again with DPBS (3x). ATAC-seq kit (Active Motif, 53150) was used to process samples according to manufacturer’s guidelines. In summary, cell pellets were resuspended with ice-cold ATAC lysis buffer and removed supernatant to isolate nuclei. Nuclei were resuspended in Tagmentation Master Mix (1x Tagmentation buffer, 1x PBS, 0.01 % digitonin, 0.1% tween-20, assembled Transposomes). The mixture was incubated at 37 °C for 30 minutes on thermal cycler with gentle mixing every 15 minutes. Samples were processed for DNA purification and library preparation according to the manufacturer’s guidelines. Samples were further cleaned by SPRI beads, followed by gel-purification to obtain fragments below 700-bp and were multiplex sequenced on the Illumina NextSeq 2000 with 50-bp paired-end reads.

### ATAC-seq analysis

Sequencing reads were first mapped to human genome build hg19 and RefSeq genes by Bowtie2 with the following parameters: bowtie2 --local --very-sensitive-local --no-mixed --no-discordant --dovetail --phred33 -I 10 -X 700 --threads 12 -x. Unmapped, duplicated, and low-quality reads were filtered out with sambamba (v0.7.1). Reads mapped to mitochondria genome and blacklist regions defined in data file wgEncodeDacMapabilityConsensusExcludable.bed.gz were also excluded by using samtools (v1.10). Enriched regions were identified by MACS2 with following parameters: -q 0.01 --nomodel --shift -100 --extsize 150. For each sample, adjacent peaks (distance < 1kb) were stitched. A merged peak list with all ATAC sites identified from all samples was created for relevant downstream analyses. Differential analyses were performed with the same method used for ChIP-seq experiments as described in the previous section. To only include the sites with strong ATAC signal in differential analyses and minimize the random noise, an arbitrary threshold (1 rpm/bp for ChIP-seq with parental 22Rv1 and 0.5 rpm/bp for ChIP-seq with engineered 22Rv1) was used to include the sites of which mean ATAC signal from the triplicate is above the threshold in at least one relevant condition. In the analyses showing the correlation of FOXA1 ChIP changes and ATAC changes, only sites with signals above both the ChIP threshold and the ATAC threshold were included to minimize the effects by random noise. The sites with significant ATAC changes are defined by <Log2FC < -1 or > 1 and *P* value <0.05. The fragment distribution was summarized by using samtools. All box plots represent 25-75 percentile with whiskers extending 1.5 interquartile range (IQR) and the center line represents the median.

### Motif analysis

For de novo motif discovery, 500-bp fragments around the center of each enriched site from each sample were extracted and were used as inputs for MEME-ChIP in MEME Suite^9^ with default setting under ‘classic mode’ to search for motif width of 10 bp. Top enriched motifs were shown in the figures. To further investigate the prevalence of the 3-bp ancillary motifs upstream of the canonical FOXA1-binding motif, the sequences of the enriched sites for each sample were extracted. The 3-bp ancillary motifs were first identified by locating the canonical motif (GTAAA[CT]A) in the sequences of a given sample and the number of the appearance of each motif was summarized for each sample. The frequencies of the ancillary 3-bp motifs among all identified motifs (64 possible motifs in total plus ‘no motif’ if the canonical motif appears at either end of the enriched region) were then calculated based on the number of their appearances in the corresponding sample.

### Quantification and statistical analysis

Unless otherwise stated, quantitative data are expressed in bar and line graphs as mean ± SEM (error bar) shown. Data quantification and statistical analysis were performed with Graphpad Prism 10 or Excel. All details are included in figure legends.

## Data and code availability

The mass spectrometry proteomics data have been deposited to the ProteomeXchange Consortium via the PRIDE^76^, partner repository with the dataset identifier PXD050685. All ChIP-seq and ATAC-seq data have been deposited to NCBI under GEO: GSE261803 and GSE261801, respectively. Processed proteomics and sequencing data are provided in Supplementary Dataset S1 and S2, respectively. All raw proteomic data and sequencing data are publicly available as of the date of publication. This paper does not report original code.

## Acknowledgements

This work was supported by the NIH (R35 CA231991 (B.F.C.), DP5-OD26380 (M.A.E.), F32 CA265211 (C.J.R.)), Ono Pharma Foundation, HHMI Hanna H Gray Fellowship (E.N., GT15176), and Jane Coffin Childs Memorial Fellowship (K.E.D.). The authors thank Melissa Dix for mass spectrometry instrumentation, Daisuke Ogasawara for the proteomic data analysis assistance, the Scripps Research Genomics Core for the NGS experiments, Xuedong Liu and Bing Chen (WuXi AppTech) for compound resynthesis, Jason Chen, Brittany Sanchez, Quynh Nguyen Wong, and Jason Lee (Scripps Automated Synthesis Facility) for support with high-resolution mass spectrometry (HRMS) and chiral SFC.

## Author contributions

S.J.W., Y.Z., B.F.C., and M.A.E. conceived the study. S.J.W., K.E.D., and E.N. generated proteomic data. S.J.W., B.M., and B.F.C. analyzed proteomic data. S.J.W. and N.S.M. performed gel-ABPP and biochemical experiments. C.J.R. synthesized compounds. S.J.W., Y.Z., and C.J.R. performed ChIP-seq experiments. S.J.W. and Y.Z. performed ATAC-seq experiments. Y.Z. and M.A.E. analyzed sequencing data. S.J.W., Y.Z., C.J.R., N.S.M., K.E.D., E.N., L.M.H., and J.R.R. provided resources. S.J.W., Y.Z., B.F.C., and M.A.E. wrote and edited the manuscript. B.F.C. and M.A.E. supervised the study.

---

**Figure S1.**
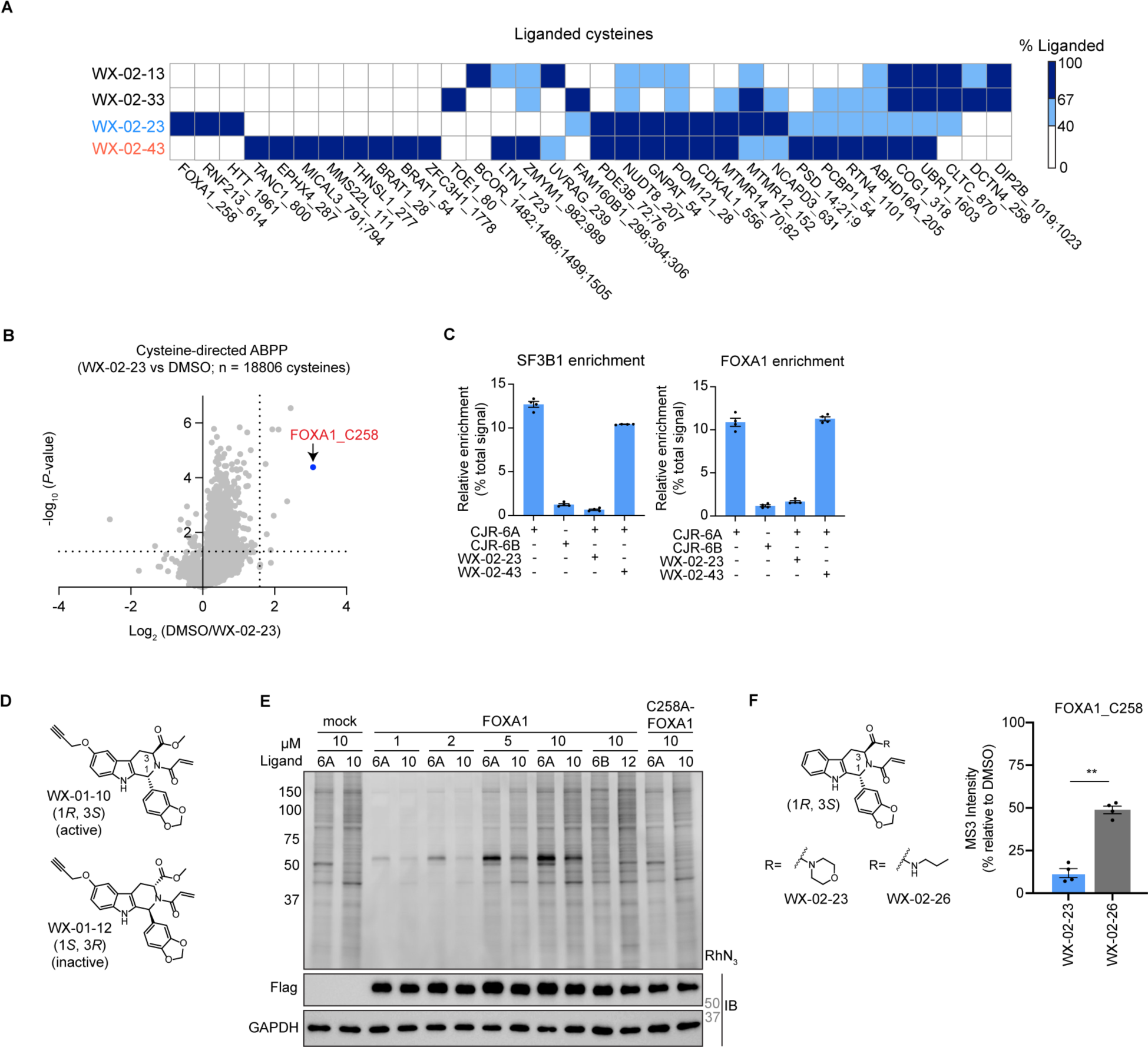
Tryptoline acrylamide WX-02-23 stereoselectively and site-specifically engages cysteine-258 (C258) in FOXA1 in human prostate cancer cells, related to Figure 1. **A.** Heatmap showing all cysteines substantially engaged by the WX-02-13/33/23/43 stereoprobe set in 22Rv1 cells (20 µM; 3 h). Cysteines shown meet the following criteria: > 66.7% IA-DTB blockade by at least one stereoprobe; quantification of ≥ 1 other unchanging cysteine (< 25% decrease in IA-DTB labeling) in the same protein. IA-DTB blockade is denoted using the following color scale: dark blue: > 66.7 %, light blue: 40-66.7%, white: < 40% probe engagement. Data are average values (N = 2 - 4 per group). **B.** Volcano plot showing global cysteine reactivity profiles for active stereoprobe WX-02-23 from cysteine-directed ABPP experiments shown in Figure 1B. Right of vertical dashed line indicates cysteines showing > 66.7 % engagement by WX-02-23. Above horizontal dashed line indicates cysteines with *P*-values < 0.05. Cysteines were required to originate from proteins with at least two quantified cysteines, and proteins where all cysteines uniformly changed were also removed to ensure only stereoprobe-induced changes in reactivity (rather than potential changes in protein expression) were interpreted. Average values from 2 independent experiments shown. *P*-values were determined by two-tailed Student’s t-test. **C.** Protein-directed ABPP data for SF3B1 and FOXA1 from 22Rv1 cells treated with WX-02-23 or WX-02-43 (20 µM, 2 h) followed by CJR-6A or CJR-6B (5 µM, 1 h). Data are average values ± SEM (N = 4 per group). **D.** Chemical structures of WX-01-10 and WX-01-12. **E.** Uncropped image from gel-ABPP experiment shown in Figure 1G. IB, immunoblots for the indicated proteins. **F.** Left, chemical structures of WX-02-23 and WX-02-26. Right, Cysteine-directed ABPP data for FOXA1_C258 in 22Rv1 cells treated with stereoprobes (20 μM, 3 h). WX-02-26 data were taken from another study (Njomen et al)^38^. N = 4 biological replicates pooled from 2 independent experiments. Data are average values ± SEM (N = 4 per group). \*\**P* < 0.01 determined by paired Student’s t-test.

**Figure S2.**
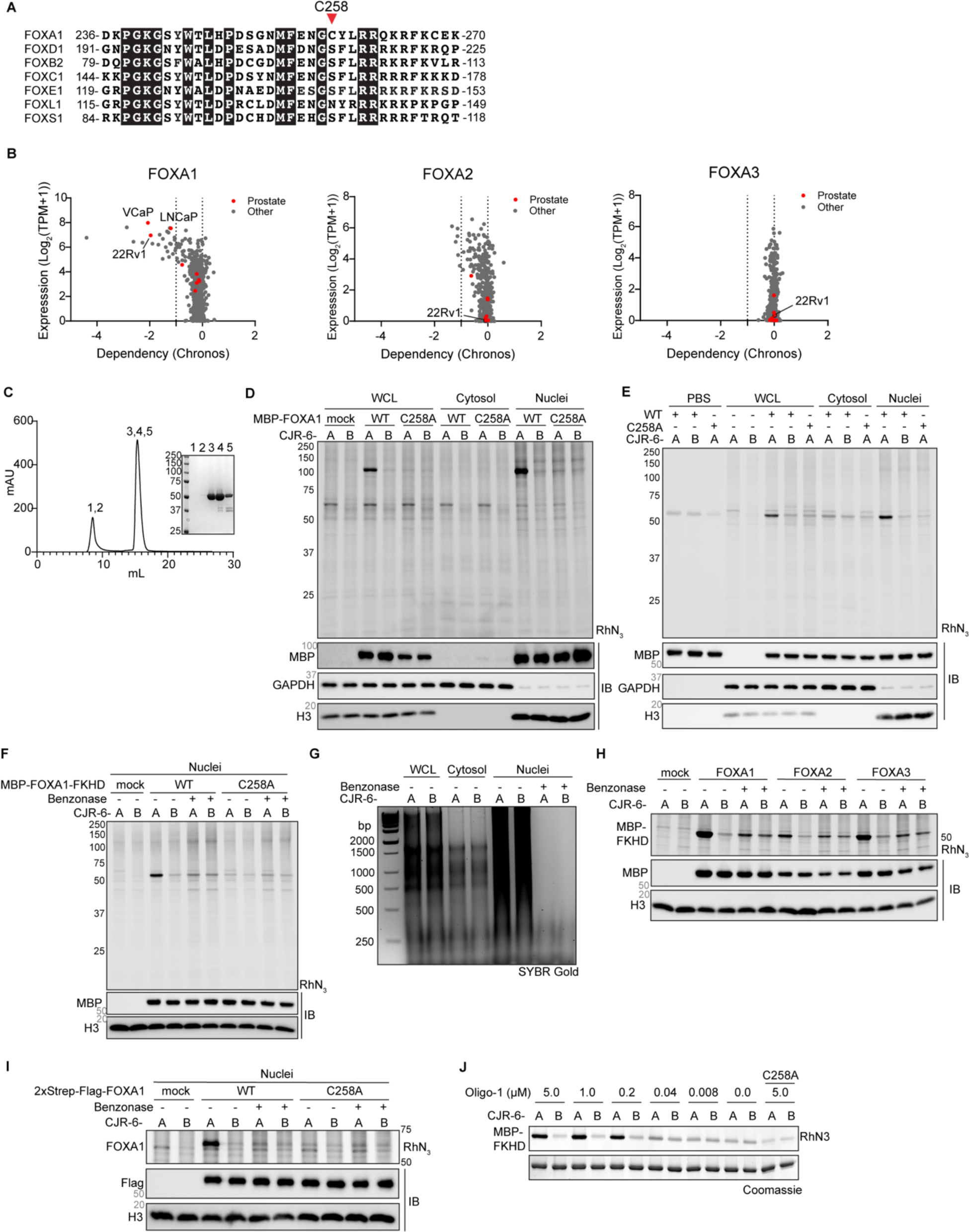
Stereoprobes engage FOXA1_C258 in a DNA-dependent manner, related to Figure 2. **A.** Protein sequence alignment of region of human FOXA1 surrounding C258 (red arrow) other representative human FOX transcription factors. **B.** *FOXA* gene dependencies across human cancer cell lines studied in the Cancer Dependency Map (https://depmap.org/portal/). Plots show *FOXA1*, *FOXA2* and *FOXA3* dependency scores (Chronos score, data release 23Q2) versus gene expression levels. Prostate cancer cell lines are highlighted in red, and the 22Rv1 cell line is marked in each plot. **C.** Size-exclusion chromatography (SEC) trace as final step of purification of recombinant MBP-FOXA1-FKHD protein. Fractions 3 and 4 were pooled and used for subsequent experiments. Coomassie stain of indicated SEC fractions (as labeled above SEC peaks). **D.** Gel-ABPP data for indicated (sub)cellular fractions of HEK293T cells recombinantly expressing WT and C258A-MBP-FOXA1 proteins (or mock cells). WCL, whole cell lysates. Each fraction was incubated with CJR-6A or CJR-6B (10 μM *in vitro*; 1 h) and analyzed by gel-ABPP as described in Figure 1G. IB, immunoblots for the indicated proteins. Data are from a single experiment representative of two independent experiments. **E.** Gel-ABPP data showing that addition of purified MBP-FOXA1-FKHD protein to HEK293T cell lysates restores stereoselective and site-specific engagement by CJR-6A. Purified WT and C258A proteins (500 nM) were incubated with the indicated (sub)cellular fractions (1 mg/mL; 50 μL) in the presence of CJR-6A or CJR-6B (10 µM, 1 h) and then analyzed by gel-ABPP as described in Figure 1G. IB, immunoblots for the indicated proteins. Data are from a single experiment representative of two independent experiments. **F.** Effect of benzonase treatment on stereoselective engagement of MBP-FOXA1-FKHD by CJR-6A. Benzonase was added to lysates from HEK293T cells recombinantly expressing WT or C258A MBP-FOXA1-FKHD protein for 30 min prior to addition of CJR-6A or CJR-6B (10 μM *in vitro*, 1 h), after which samples were analyzed by gel-ABPP. IB, immunoblots for the indicated proteins. Data are from a single experiment representative of two independent experiments. **G.** An agarose gel confirming DNA degradation following benzonase treatment protocol used in Figures 2I and **S2F.** **H.** Effect of benzonase treatment on stereoselective engagement of MBP-FOXA1-FKHD, MBP-FOXA2-FKHD, and MBP-FOXA3-FKHD by CJR-6A. Each construct was ectopically expressed in HEK293T cells, followed by cell fractionation and benzonase treatment. Cell lysates were treated with CJR-6A or CJR-6B (10 µM *in vitro*, 1 h) and then analyzed by gel-ABPP. IB, immunoblots for the indicated proteins. Data are from a single experiment representative of two independent experiments. **I.** Gel-ABPP data showing that stereoselective engagement of 2xStrep-Flag epitope-tagged full-length WT-FOXA1 by CJR-6A and the benzonase sensitivity of this interaction in nuclear lysates from HEK293T cells. HEK293T cells recombinantly expressing WT- or C258A-FOXA1 were lysed and fractionated, after which the nuclear fraction was treated with benzonase (30 min, 37 °C), followed by CJR-6A or CJR-6B (10 μM *in vitro*, 1 h), and analyzed by gel-ABPP. IB, immunoblots for the indicated proteins. Data are from a single experiment representative of two independent experiments. **J.** Concentration-dependent effects of oligo-1 on stereoselective engagement of MBP-FOXA1-FKHD by CJR-6A. Purified MBP-FKHD-FOXA1 (0.5 µM) was incubated with the indicated concentrations of oligo-1 for 30 min, followed by treatment with CJR-6A or CJR-6B (10 μM, 1 h), and analysis by gel-ABPP.

**Figure S3.**
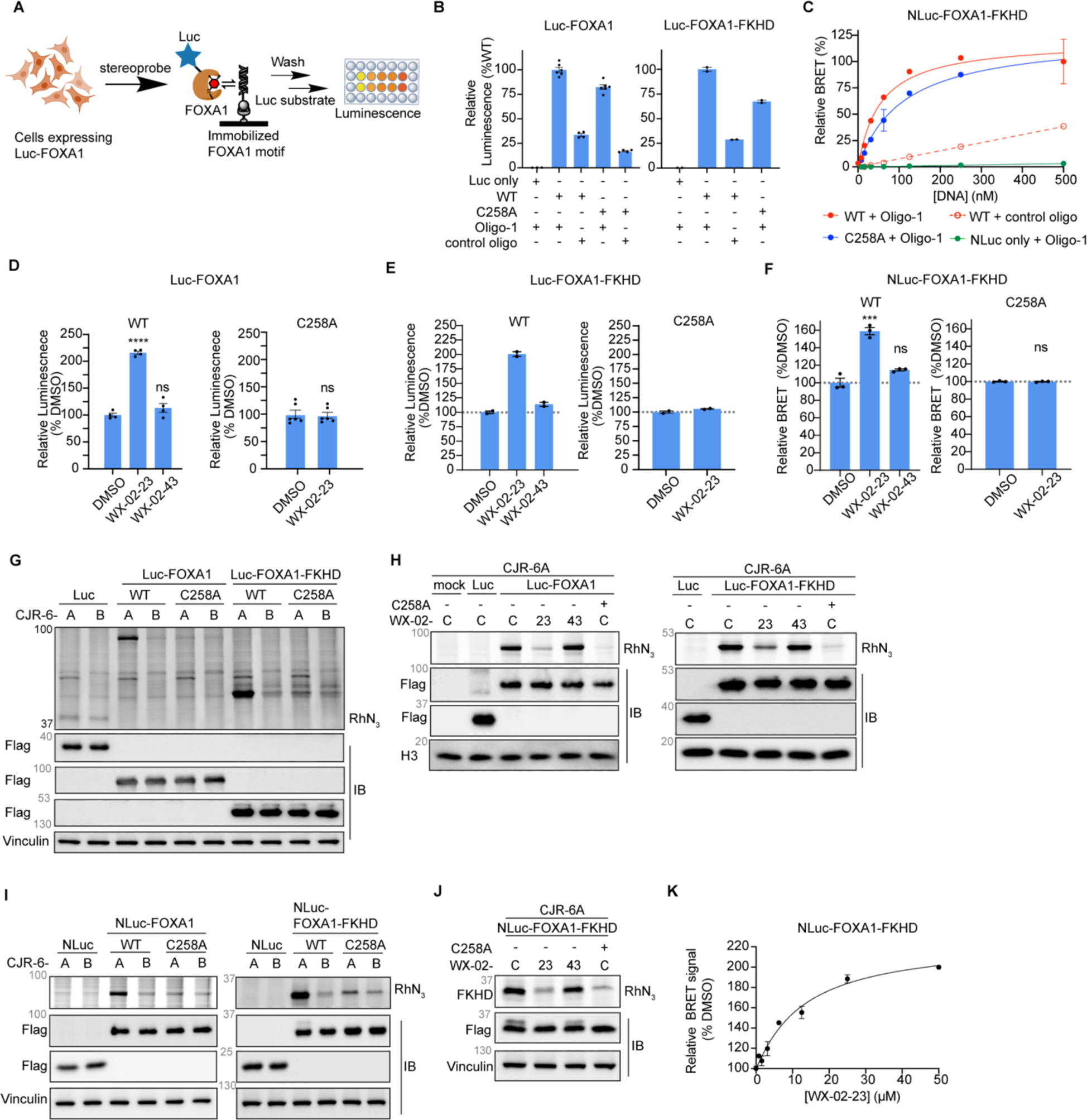
Stereoprobes enhance FOXA1-DNA interactions *in vitro*, related to Figure 3. **A.** Schematic for luciferase (Luc)-based immobilized oligonucleotide assay to measure stereoprobe effects on FOXA1-DNA interactions. HEK293T cells recombinantly expressing a Luc-FOXA1 fusion protein (N-terminally tagged) were treated with WX-02-23 or WX-02-43 (20 µM, 3 h), after which the nuclear proteome was isolated and incubated with biotinylated oligo-1 or control DNA oligonucleotide lacking the canonical FOXA1-binding motif pre-immobilized onto a streptavidin-coated plate. Following washing of the plate, Luc substrate was added and luminescence was measured. **B.** Binding profiles of Luc-FOXA1 or Luc-FOXA1-FKHD proteins (WT and C258A) to oligo-1 in Luc-based immobilized oligonucleotide assay. ‘Luc only’ indicates HEK293T cells expressing the Luc protein. Nuclear proteomes containing indicated constructs were incubated with indicated oligonucleotides immobilized on streptavidin-coated plate (1 h), after which Luc substrate was added and measured. Data average values ± SEM and normalized to each WT + oligo-1 condition; N = 3-6 per group for Luc-FOXA1 and N = 2 per group for Luc-FOXA1-FKHD. **C.** Binding curves for nanoBRET-based oligonucleotide-binding assay data for oligo-1 and control oligo with NLuc-FOXA1-FKHD fusion proteins (red: WT and blue: C258A) assayed in lysates of transfected HEK293T cells. Dashed line indicates binding profile of WT protein with control oligonucleotide, and green indicates oligo-1 with a NLuc control protein. NanoBRET signals are reported after subtraction of background signals from samples lacking TAMRA-oligonucleotides. Data are average values ± SEM from two independent experiments. **D.** WX-02-23 stereoselectively increases Luc-WT-FOXA1-DNA interactions, but not Luc-C258A-FOXA1-DNA interactions, in the immobilized oligonucleotide assay. Experiments were performed as described in **Figures S3A**. Data were normalized to DMSO controls for each protein. Data are average values ± SEM; N = 4 for WT-FOXA1 and N = 6 for C258A-FOXA1. Two-tailed t-test \*\*\*\**P* <0.0001 in comparison to DMSO-treated HEK293T cells. **E.** The same experiment as **Figures S3D**, except Luc-FOXA1-FKHD constructs were used. Data are average values ± SEM; N = 2 per group. **F.** The same experiment as Figure 3D, except NLuc-FOXA1-FKHD constructs (WT and C258A) were used. Effects of WX-02-23 and WX-02-43 on NLuc-FOXA1-FKHD interactions as measured by the nanoBRET-based oligonucleotide-binding assay data. HEK293T cells expressing each FOXA1 protein were treated with WX-02-23 or WX-02-43 (20 µM, 3 h), or DMSO, followed by cell lysis and incubation with oligo-1 (15 nM, 20 min), and nanoBRET analysis. Left and right bar graphs within each panel show effects of stereoprobes on WT-FOXA1 and C258A-FOXA1 proteins, respectively. NanoBRET data were normalized to DMSO controls for each protein. Data are average values ± SEM; N = 3 per group. Two-tailed t-test \*\*\**P* < 0.001 compared to DMSO control. **G.** Gel-ABPP data showing stereoselective and site-specific engagement of Luc-FOXA1 and Luc-FOXA1-FKHD proteins by CJR-6A. Data are from HEK293T cells recombinantly expressing the indicated FOXA1 proteins treated with CJR-6A or CJR-6B (10 µM *in vitro*, 1 h) and analyzed by gel-ABPP. IB, immunoblots for the indicated proteins. Data are from a single experiment representative of two independent experiments. **H.** Gel-ABPP data showing stereoselective engagement of Luc-FOXA1 (left) and Luc-FOXA1-FKHD (right) by WX-02-23. HEK293T cells expressing the indicated WT- and C258A-FOXA1 proteins were treated with DMSO control (C), WX-02-23, or WX-02-43 (20 μM of compounds, 2 h), followed by CJR-6A (5 μM, 1 h) and analyzed by gel-ABPP. IB, immunoblots for the indicated proteins. Data are from a single experiment representative of two independent experiments. **I.** Gel-ABPP data showing stereoselective and site-specific engagement of NLuc-FOXA1 and NLuc-FOXA1-FKHD constructs by CJR-6A. Data are from HEK293T cells recombinantly expressing the indicated FOXA1 proteins treated with CJR-6A or CJR-6B (10 µM *in vitro*, 1 h) and analyzed by gel-ABPP. IB, immunoblots for the indicated proteins. Data are from a single experiment representative of two independent experiments. **J.** Gel-ABPP data showing stereoselective engagement of NLuc-FOXA1-FKHD fusion proteins by WX-02-23. HEK293T cells expressing the indicated NLuc-FOXA1-FKHD proteins were treated with WX-02-23 (23) or WX-02-43 (43), (20 μM, 2 h), followed by CJR-6A (5 μM, 1 h), and analyzed by gel-ABPP. C= DMSO control. IB, immunoblots for the indicated proteins. Data are from a single experiment representative of two independent experiments. **K.** Concentration-dependent effects of WX-02-23 (3 h) on nanoBRET signal for NLuc-FOXA1-FKHD with oligo-1. HEK293T cells expressing NLuc-FOXA1-FKHD were treated with the indicated concentrations of WX-02-23 for 3 h, followed by cell lysis and incubation with TAMRA-Oligo 1 oligonucleotide (15 nM, 20 min), and nanoBRET analysis. Data are average values ± SEM and presented as the relative maximum percentage increase of nanoBRET signal in WX-02-23-treated samples over DMSO-treated control samples (where DMSO-treated controls are set to 100%); N = 2 per group.

**Figure S4.**
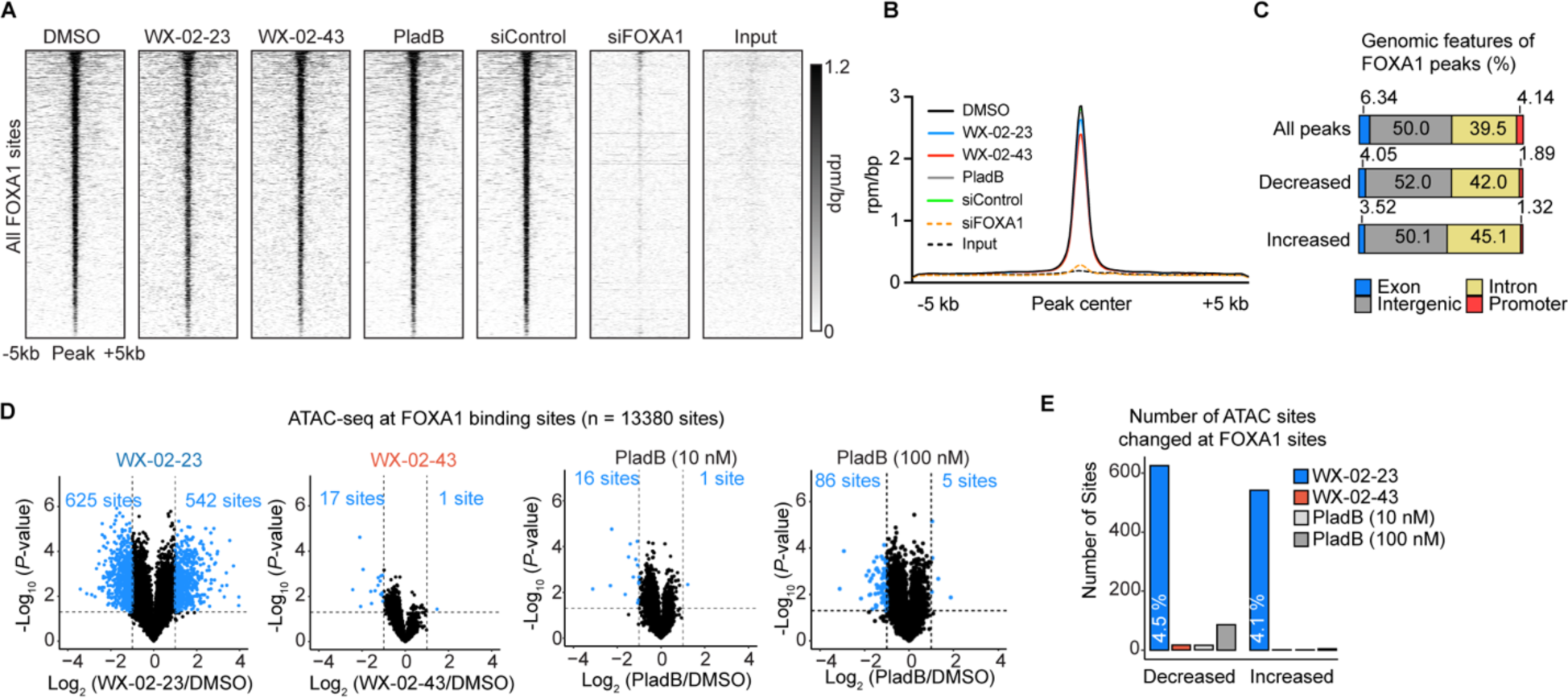
WX-02-23 redirects the pioneering activity of FOXA1 in prosate cancer cells. related to Figure 4. **A.** Rank-ordered heatmaps showing FOXA1 ChIP-seq signal. Binding sites are ranked by the FOXA1 signal in the DMSO sample (total 30,557 sites) and signal intensity is plotted by color intensity. **B.** Meta-track representations of average FOXA1 ChIP signal (rpm/bp) in each sample at merged binding sites identified with a fixed width of 10 kb window centered around the center of the site. **C.** Genomic feature distribution of FOXA1-binding sites identified in ChIP-seq experiments at all sites and sites with FOXA1 binding stereoselectively changed by WX-02-23 compared to DMSO. Increased: log_2_(WX-02-23/DMSO) > 2 and Log_2_(WX-02-43/DMSO) < 1; *N* = 681. Decreased: log_2_(WX-02-23/DMSO) < -2 and Log_2_(WX-02-43/DMSO) > -1; *N* = 1,482. All peaks: *N* = 29,831. Sites with substantial ChIP signal (rpm/bp > 1) in at least 1 treatment condition were used for analyses. **D.** Chromatin accessibility (ATAC-seq) changes induced by compound treatments (20 µM for each stereoprobe and 10 or 100 nM for PladB) compared to DMSO control at FOXA1-binding sites in 22Rv1 cells. The FOXA1 binding sites were determined from FOXA1 ChIP-seq experiments (see Figure 4A). ATAC-seq sites with substantial ((|log_2_FC| > 1) and significant (*P* < 0.05) changes relative to DMSO are shown as blue dots. FOXA1-binding sites with substantial ATAC-seq signal (rpm/bp > 1) in at least 1 treatment condition were used for all analyses. Data are average values from three independent experiments, and *P* values were determined by two-tailed Welch’s *t*-test. The number of significantly changed sites are shown. **E.** The number of ATAC sites substantially and significantly changed for each compound treatment group at FOXA1 binding sites. The percentages of such sites compared to all FOXA1-binding sites are shown for WX-02-23 treatment. Increased: log_2_FC > 1, *P* < 0.05. Decreased: log_2_FC < -1, *P* < 0.05.

**Figure S5.**
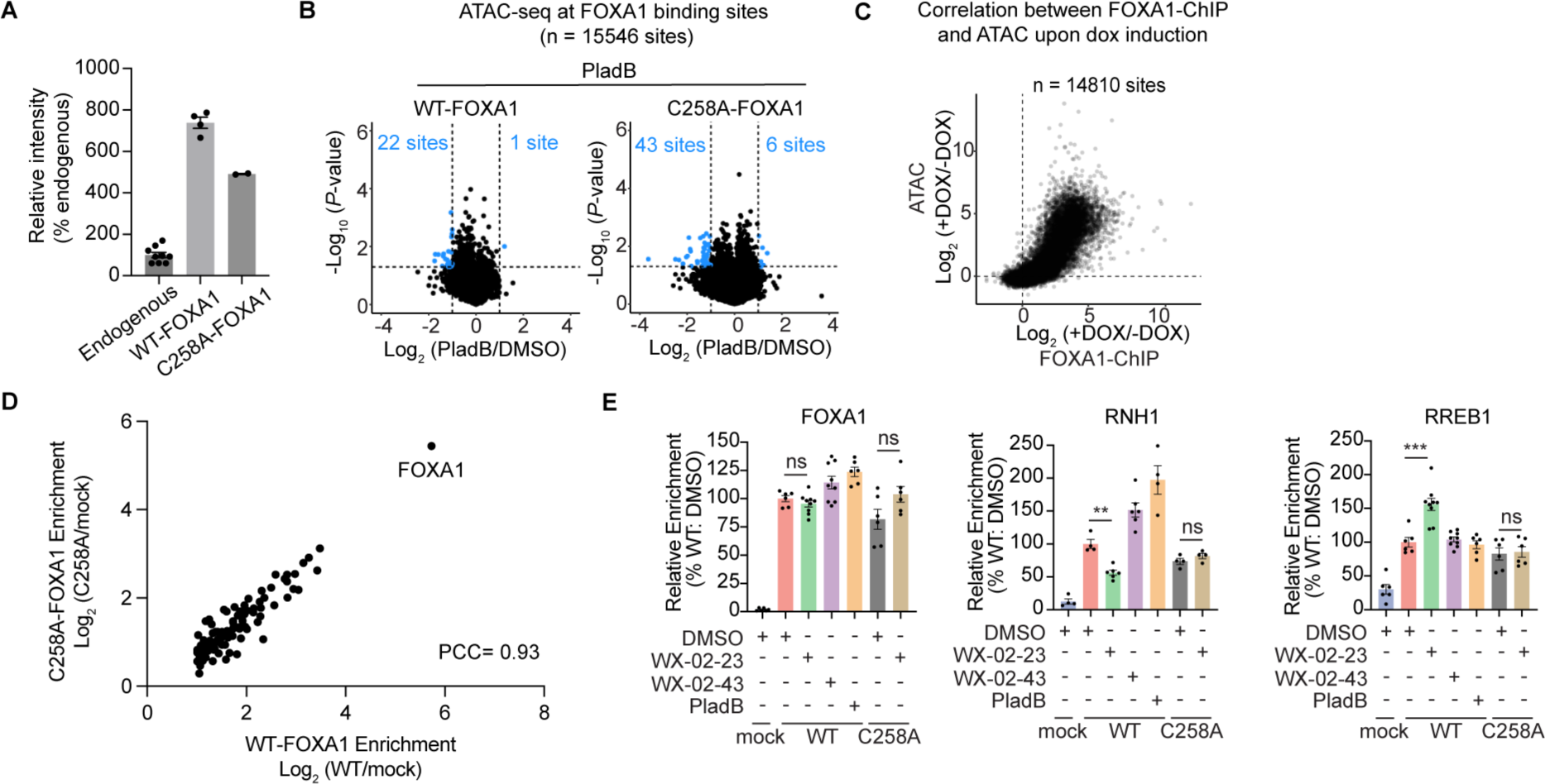
WX-02-23 effects on FOXA1-chromatin interactions depend on engagement of C258, related to Figure 5. **A.** Quantification of FOXA1 expression as measured by immunoblotting in Figure 5A. Data are average value after normalization to endogenous FOXA1. N = 2-9 per group. **B.** Chromatin accessibility (ATAC-Seq) changes induced by PladB (100 nM, 3 h) at FOXA1 binding sites in 22Rv1 Y36C-PHF5A cells recombinantly expressing WT- or C258A-FOXA1 via doxycycline induction. ATAC-seq sites with substantial ((|log_2_FC| > 1) and significant (*P* < 0.05) changes relative to DMSO are shown as blue dots. Sites with ATAC-seq signal below < 0.5 rpm/bp were excluded from the analysis. Data are average values from two independent experiments. *P* values were determined by two-tailed Welch’s *t*-test. **C.** Correlation of changes in FOXA1 binding and changes in chromatin accessibility induced by doxycycline induction of recombinant WT-FOXA1 expression in 22Rv1 Y36C-PHF5A cells. **D.** Correlation plot of proteins significantly (> 2-fold vs mock, *P* < 0.05) enriched in anti-Flag IP-MS experiments performed on 2xStrep-Flag-WT-versus C258A-FOXA1. 22Rv1 Y36C-PHF5A cells were treated with doxycycline (24 h) prior to IP-MS analysis. Data are average values. N = 4-6 per group. PCC = Pearson’s correlation coefficient. **E.** Bar graphs showing relative quantification of FOXA1, RNH1, and RREB1 in anti-Flag IP-MS experiments performed across the different indicated treatment groups (20 μM stereoprobes, 100 nM PladB; 3h). Data are normalized to DMSO-treated 2xStrep-Flag-WT-FOXA1-expressing cells. Data are average values ± SEM. N = 4-9 per group. Two-tailed Welch’s *t*-test between DMSO and WX-02-23. \*\*\**P* < 0.001, *\*\*P <* 0.01, ns = not significant.

**Figure S6.**
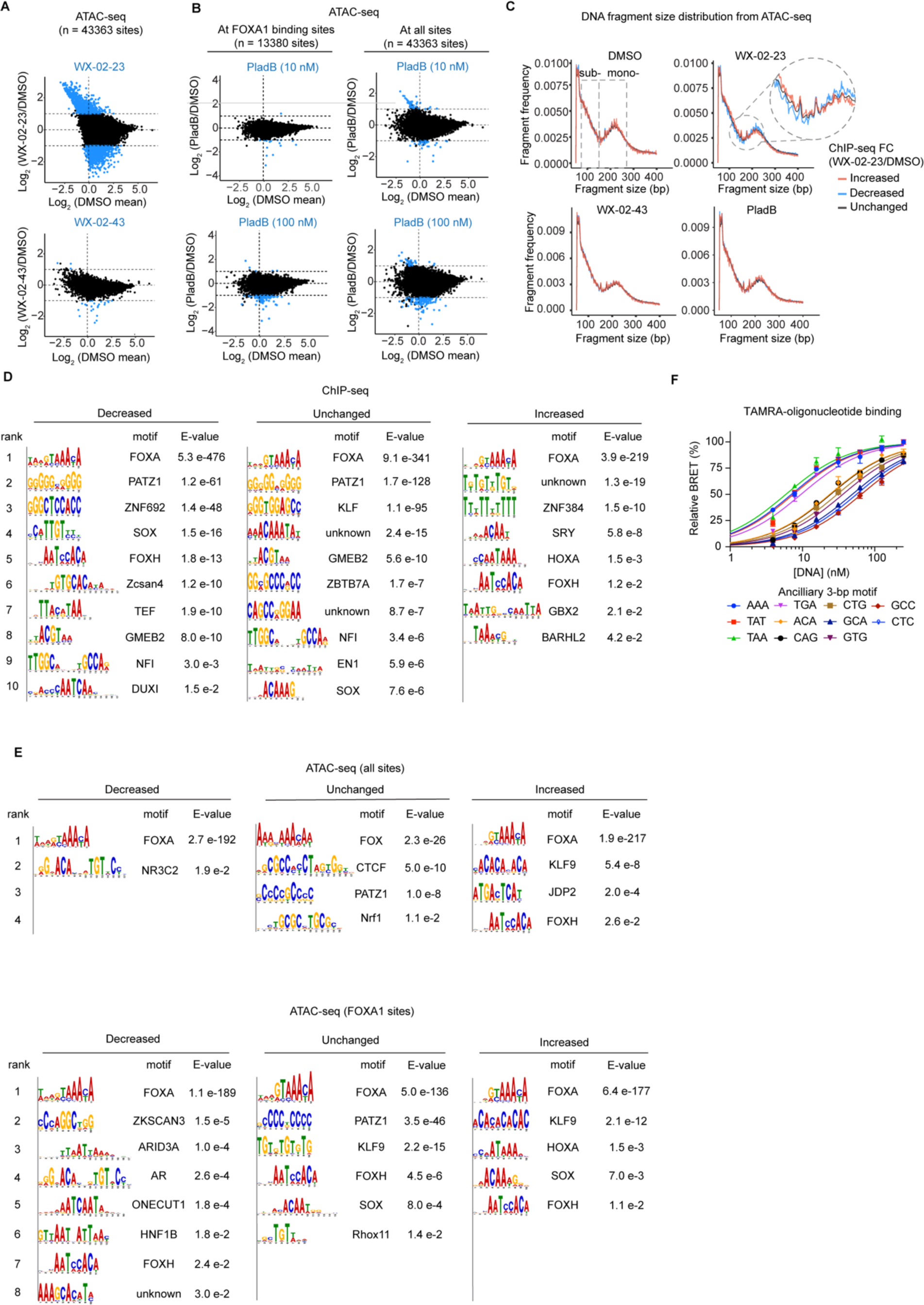
WX-02-23 remodels chromatin interactions of FOXA1 by relaxing its DNA-binding motif preference, related to Figure 6. **A.** MA plots as shown in Figures 6A, but for all ATAC-seq sites (rpm/bp > 1). **B.** Mean-average plots showing chromatin accessibility (ATAC-seq) changes induced by PladB (upper = 10 nM, below = 100 nM; 3 h) at FOXA1-binding sites (left) and at all sites (right) in 22Rv1 cells. The FOXA1 binding sites are determined from FOXA1 ChIP-seq (see Figure 4). The vertical line represents 10th percentile of DMSO mean. FOXA1-binding sites with substantial ATAC-seq signal (rpm/bp > 1) in at least 1 treatment condition are used for all analyses. ATAC-seq sites with substantial and significant changes are shown as blue dots (|log_2_FC| > 1, *P* < 0.05 as determined in Figures 4C and **S4D**). *N* = 3 biological replicates. Data are averaged and *P* values were determined by two-tailed Welch’s *t*-test. **C.** DNA fragment length distribution measured by ATAC-seq at FOXA1 binding sites in 22Rv1 cells. FOXA1-binding sites are grouped based on FOXA1 ChIP-seq changes induced by WX-02-23 relative to DMSO (decreased: log_2_FC < -2, 528 sites; unchanged: |log_2_FC| < 1, 9,475 sites; increased: log_2_FC > 2, 329 sites). Subnucleosomal and mononucleosomal fragments are highlighted in dotted grey boxes. **D.** Table showing significantly (E-value ≤ 0.05) enriched motifs revealed by d*e novo* motif discovery (MEME Suite software) in FOXA1 ChIP-seq. Separate analyses were conducted to evaluate motifs in sites that were stereoselectively increased, decreased, or unchanged by WX-02-23 relative to DMSO. The number of sites and criteria are the same as Figure 6D. The name of the best-matched known motif is shown beside each sequence. For the decreased and unchanged sites, the top 10 significant enriched motifs are shown. **E.** Table showing all significantly (E-value ≤ 0.05) enriched motifs revealed by d*e novo* motif discovery (MEME Suite software) in ATAC-seq (upper: all sites; below: at FOXA1 sites). Separate analyses were conducted to evaluate motifs in sites that were substantially and significantly increased, decreased, or unchanged by WX-02-23 relative to DMSO. The number of sites and criteria are the same as Figure 6E. The name of the best-matched known motif is shown beside each sequence. **F.** Binding curves of NLuc_FOXA1 with oligo-1 variants containing the indicated ancillary 3-bp sequences as measured with the NanoBRET assay described in Figure 3A. Data are average values ± SEM from two independent experiments.

