## Supplementary material for "Redirecting the pioneering function of FOXA1 with covalent small molecules": Methods S1

**Instruments and Materials.**

Solvents and reagents were purchased and used as received from commercial vendors. Analytical thin layer chromatography (TLC) was performed on Millipore Sigma TLC Silica gel 60G $F_{254}$ 25 glass plates (20 x 20 cm) and visualized by UV light (254 nm). Preparative thin layer chromatography (prep-TLC) was performed on Miles Scientific silica gel GF UV254 plates (20 x 20 cm; 500, 1000, or 2000 μm thickness).

$^{1}H$ and $^{13}C$ NMR spectra were acquired at 25 °C on a Bruker AV III HD 600 MHz NMR spectrometer. Chemical shifts (δ) are reported in parts per million (ppm) relative to the residual solvent peaks. The following abbreviations are used to describe coupling constants (*J*, in Hertz): singlet (s), doublet (d), triplet (t), quartet (q), quintet (quint), multiplet (m), broad singlet (bs) or a combination thereof. Spectra were visualized and analyzed using MestReNova (version 14.2).

Routine LC-MS analysis was performed on a Waters I-Class LC and a Waters Acquity QDA MS with a Waters Cortecs C18 column (1.6 um, 2.1x55 mm) using a 5 - 99% B gradient lasting 2.5 minutes (A: 0.1% aqueous formic acid, B: 0.06% formic acid in acetonitrile, 0.8 mL/min flow rate, 35 °C column temperature). The product purity was quantified on a combined 220 nm + 260 nm UV channel and the product identity was verified by mass spectrometry.

Chiral, analytical supercritical fluid chromatography (SFC) was performed on a Waters UPC2 SFC with a Daicel IH column (3 μm, 4.6 x 250 mm) under isocratic conditions (3.3 mL/min, 50% MeOH/$CO_2$, 1600 psi backpressure) at 30 °C. The enantiomers were detected by UV light (212 nm). Instead of a racemate, methods were developed using a mixture of independently prepared enantio-enriched samples in an ~1:1 ratio.

Mass measurements for high-resolution mass spectrometry (HRMS) were performed on a Waters Xevo G2-XS TOF calibrated against sodium formate clusters and using a LeuEnk lockmass. Expected monoisotopic masses were calculated using MassLynx 4.1 and the m/z values for calibrant and lockmass were MassLynx-default values.

All final compounds are greater than 95% pure by LC-MS and greater than 95% ee prior to use in biological experiments.

### Synthetic Procedures.

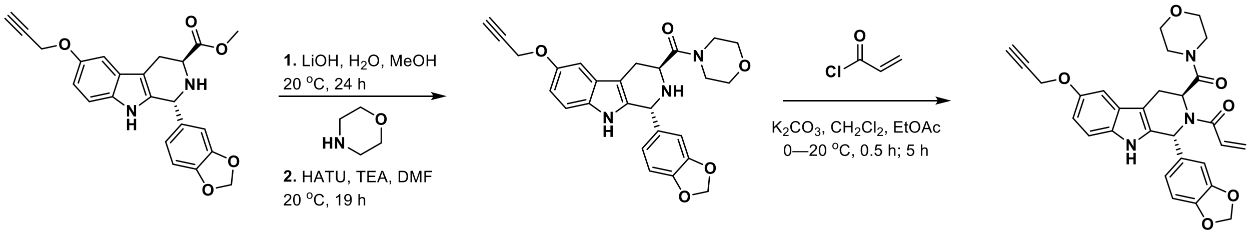

S15 S15.1 CJR-6A

#### ((1*R*,3*S*)-1-(benzo[d][1,3]dioxol-5-yl)-6-(prop-2-yn-1-yloxy)-2,3,4,9-tetrahydro-1H-pyrido[3,4-b]indol-3-yl)(morpholino)methanone (S15.1)

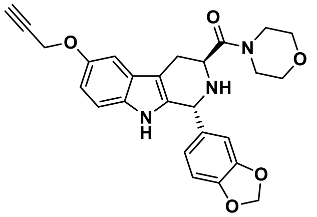

To a solution of **S15**[1] (52.5 mg, 0.13 mmol, 1 equiv.) in methanol (0.65 mL) was added lithium hydroxide monohydrate (8.54 mg, 0.204 mmol, 1.6 equiv.) and water (23.4 mg, 1.30 mmol, 23.4 µL, 10 equiv.). The mixture was stirred at room temperature for ~24 h. The reaction was monitored by LCMS. Once complete, reaction mixture was concentrated under reduced pressure to afford a residue which was azeotroped once with toluene (1 mL) to remove residual water. The resulting solid was dissolved in DMF (0.3 mL) and treated with HATU (74.1 mg, 0.195 mmol), DIPEA (33.6 mg, 0.260 mmol, 45.3 µL) and morpholine (227 mg, 2.60 mmol, 0.22 mL). The mixture was stirred at room temperature for ~19 h. The reaction was diluted in EtOAc and partitioned against brine. The product was extracted 3x with EtOAc and the combined organic layers were dried over anhydrous sodium sulfate. The solution was concentrated and purified via preparatory TLC (silica) eluting with EtOAc. Pure product was recovered in EtOAc and concentrated to afford a foam (17.8 mg, 38.7 µmol, 30% yield over two steps).

**LC-MS** m/z calculated for $[M+H]^+$ 460.19. Found. 460.19. Retention time: 0.95 min.

**HRMS** m/z calculated for $[M+H]^+$ 460.1872 Found 460.1870.

**$^1$H NMR** (600 MHz, $CD_3OD$) δ 7.17 (d, *J* = 8.7 Hz, 1H), 7.09 (d, *J* = 2.4 Hz, 1H), 6.80 (dd, *J* = 8.7, 2.5 Hz, 1H), 6.77 – 6.70 (m, 2H), 6.65 – 6.58 (m, 1H), 5.91 (m, 2H), 5.20 (s, 1H), 4.72 (d, *J* = 2.4 Hz, 2H), 3.93 (dd, *J* = 10.2, 4.7 Hz, 1H), 3.71 – 3.50 (m, 5H), 3.50 – 3.42 (m, 1H), 3.29 – 3.20 (m, 2H), 2.95 (ddd, *J* = 15.4, 10.3, 1.3 Hz, 1H), 2.89 (t, *J* = 2.4 Hz, 1H), 2.87 (dd, *J* = 15.4, 4.7 Hz, 1H), 2 exchangeable protons not observed.

**$^{13}$C NMR** (151 MHz, $CD_3OD$) δ 173.49, 153.15, 149.17, 148.55, 137.17, 135.43, 133.74, 128.43, 123.42, 113.12, 112.55, 110.40, 109.43, 108.69, 103.43, 102.44, 80.67, 76.16, 67.85, 67.71,

57.87, 56.08, 49.33, 47.03, 43.56, 25.70.

**1-((1*S*,3*R*)-1-(benzo[d][1,3]dioxol-5-yl)-3-(morpholine-4-carbonyl)-6-(prop-2-yn-1-yloxy)-1,3,4,9-tetrahydro-2H-pyrido[3,4-b]indol-2-yl)prop-2-en-1-one (CJR-6A)**

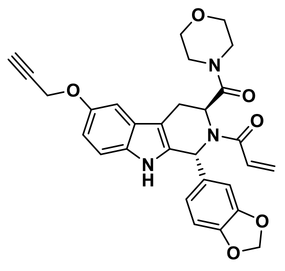

To an ice-cold solution of **S15.1** (17.8 mg, 38.7 µmol, 1 equiv.) in EtOAc (0.78 mL) was added potassium carbonate (16.1 mg, 0.116 mmol, 3 equiv.) and 0.45 M solution of acryloyl chloride in $CH_2Cl_2$ (0.13 mL, 58 µmol, 1.5 equiv.). The mixture was stirred at the same temperature for 30 min. Next, the ice bath was removed, and the reaction allowed to continue at room temperature for 5 h. The reaction was monitored by TLC. The reaction mixture was diluted in EtOAc, quenched with aq. sat. sodium bicarbonate, and then extracted 3x with EtOAc. The organic layer was dried over sodium sulfate and then concentrated under reduced pressure to give a residue. The product was purified via preparatory TLC (silica) eluting with 75% EtOAc/hex. Pure product was recovered in EtOAc, concentrated to an oil, crashed out in diethyl ether with sonication, and dried to under reduced pressure to afford an off-white solid (4.4 mg, 8.57 µmol, 22% yield).

**LC-MS** m/z calculated for $[M+H]^+$ 514.19. Found. 514.20. Retention time: 1.32 min.

**HRMS** m/z calculated for $[M+H]^+$ 514.18998. Found. 514.1987.

**$^1$H NMR** (600 MHz, $CD_3OD$) δ 7.17 (d, *J* = 8.7 Hz, 1H), 7.08 (d, *J* = 2.5 Hz, 1H), 6.98 – 6.87 (m, 2H), 6.85 (d, *J* = 7.7 Hz, 1H), 6.79 (dd, *J* = 8.8, 2.5 Hz, 2H), 6.35 (s, 1H), 6.23 (dd, *J* = 16.6, 1.8 Hz, 1H), 5.92 (d, *J* = 4.4 Hz, 2H), 5.74 (dd, *J* = 10.6, 1.8 Hz, 1H), 5.05 (s, 1H), 4.71 (d, *J* = 2.4 Hz, 2H), 3.65 – 3.33 (m, 6H), 3.30 – 3.05 (m, 4H), 2.88 (t, *J* = 2.4 Hz, 1H), 1 exchangeable proton not observed.

**$^{13}$C NMR** (151 MHz, $CD_3OD$) δ 171.26, 169.76, 153.36, 149.73, 149.11, 136.03, 134.44, 133.94, 129.84, 129.72, 127.85, 122.08, 113.50, 112.83, 109.37, 108.72, 107.29, 103.26, 102.78, 80.58, 76.22, 67.73, 67.34, 59.20, 57.73, 53.80, 47.09, 43.93, 23.93.

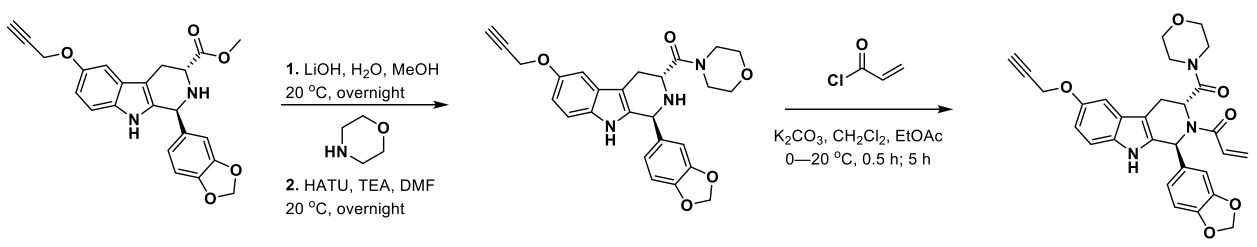


*ent*-S15 ent-S15.1 CJR-6B

**((1*S*,3*R*)-1-(benzo[d][1,3]dioxol-5-yl)-6-(prop-2-yn-1-yloxy)-2,3,4,9-tetrahydro-1H-pyrido[3,4-b]indol-3-yl)(morpholino)methanone (*ent*-S15.1)** Prepared from *ent*-S15[1] in analogous fashion to the synthesis of S15.1 (302 µmol scale, 47% yield over two steps).

**LC-MS** m/z calculated for $[M+H]^+$ 460.19. Found. 460.19. Retention time: 0.95 min.

**HRMS** m/z calculated for $[M+H]^+$ 460.1872 Found. 460.1866.

**$^1$H NMR** (600 MHz, $CD_3OD$) δ 7.17 (d, *J* = 8.7 Hz, 1H), 7.10 (d, *J* = 2.4 Hz, 1H), 6.80 (dd, *J* = 8.7, 2.5 Hz, 1H), 6.77 – 6.72 (m, 2H), 6.63 (dd, *J* = 8.2, 1.8 Hz, 1H), 5.91 (dd, *J* = 1.1, 1.1 Hz, 2H), 5.23 (s, 1H), 4.72 (d, *J* = 2.4 Hz, 2H), 3.95 (dd, *J* = 10.1, 4.9 Hz, 1H), 3.71 – 3.52 (m, 5H), 3.50 – 3.44 (m, 1H), 3.27 (s, 2H), 2.95 (ddd, *J* = 15.4, 10.1, 1.3 Hz, 1H), 2.92 – 2.86 (m, 2H), 2 exchangeable protons not observed.

**$^{13}$C NMR** (151 MHz, $CD_3OD$) δ 173.31, 153.17, 149.20, 148.63, 136.87, 135.19, 133.76, 128.38, 123.50, 113.18, 112.57, 110.41, 109.35, 108.72, 103.44, 102.47, 80.65, 76.16, 67.84, 67.70, 57.87, 56.10, 49.37, 47.02, 43.57, 25.64.

**1-((1*R*,3)-1-(benzo[d][1,3]dioxol-5-yl)-3-(morpholine-4-carbonyl)-6-(prop-2-yn-1-yloxy)-1,3,4,9-tetrahydro-2H-pyrido[3,4-b]indol-2-yl)prop-2-en-1-one (CJR-6B)**

Prepared from *ent*-S15.1 in analogous fashion to the synthesis of CJR-6A (126 µmol scale, 38% yield).

**LC-MS** m/z calculated for $[M+H]^+$ 514.19. Found 514.20. Retention time: 1.32 min.

**HRMS** m/z calculated for $[M+H]^+$ 514.18998. Found. 514.1987.

**$^1$H NMR** (600 MHz, $CD_3OD$) δ 7.17 (d, *J* = 8.7 Hz, 1H), 7.08 (d, *J* = 2.4 Hz, 1H), 6.99 – 6.86 (m, 2H), 6.85 (d, *J* = 8.0 Hz, 1H), 6.79 (dd, *J* = 8.7, 2.5 Hz, 2H), 6.35 (s, 1H), 6.23 (d, *J* = 16.6 Hz, 1H), 5.92 (s, 2H), 5.74 (dd, *J* = 10.7, 1.8 Hz, 1H), 5.05 (s, 1H), 4.71 (d, *J* = 2.4 Hz, 2H), 3.66 – 3.33 (m, 6H), 3.29 – 3.11 (m, 4H), 2.88 (t, *J* = 2.4 Hz, 1H), 1 exchangeable proton not observed.

$^{13}$**C NMR** (151 MHz, $CD_3OD$) δ 171.26, 169.75, 153.36, 149.73, 149.10, 136.03, 134.43, 133.94, 129.81, 129.75, 127.84, 122.08, 113.50, 112.83, 109.37, 108.71, 107.29, 103.26, 103.24, 102.77, 80.59, 80.57, 76.27, 76.18, 67.73, 67.34, 59.20, 57.75, 57.72, 57.69, 53.81, 47.09, 43.93, 23.93.

### NMR Spectra.

$^{1}H$ NMR (600 MHz, $CD_3OD$) for S15.1.

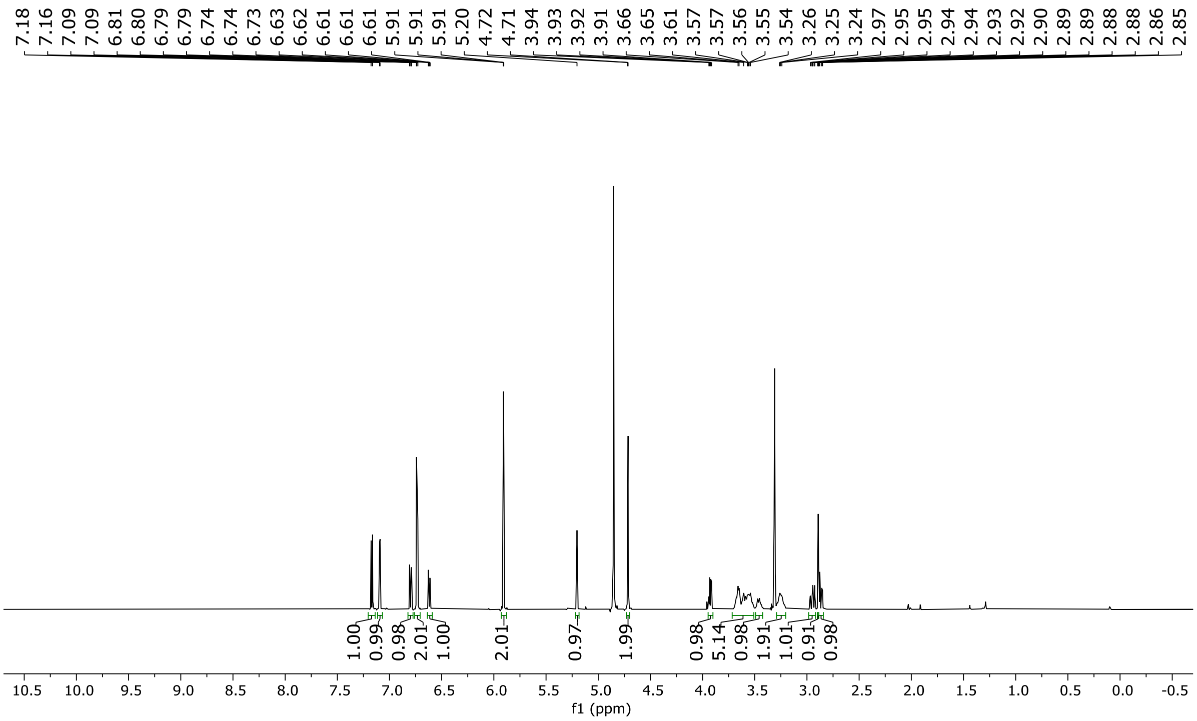

$^{13}C$ NMR (151 MHz, $CD_3OD$) for S15.1.

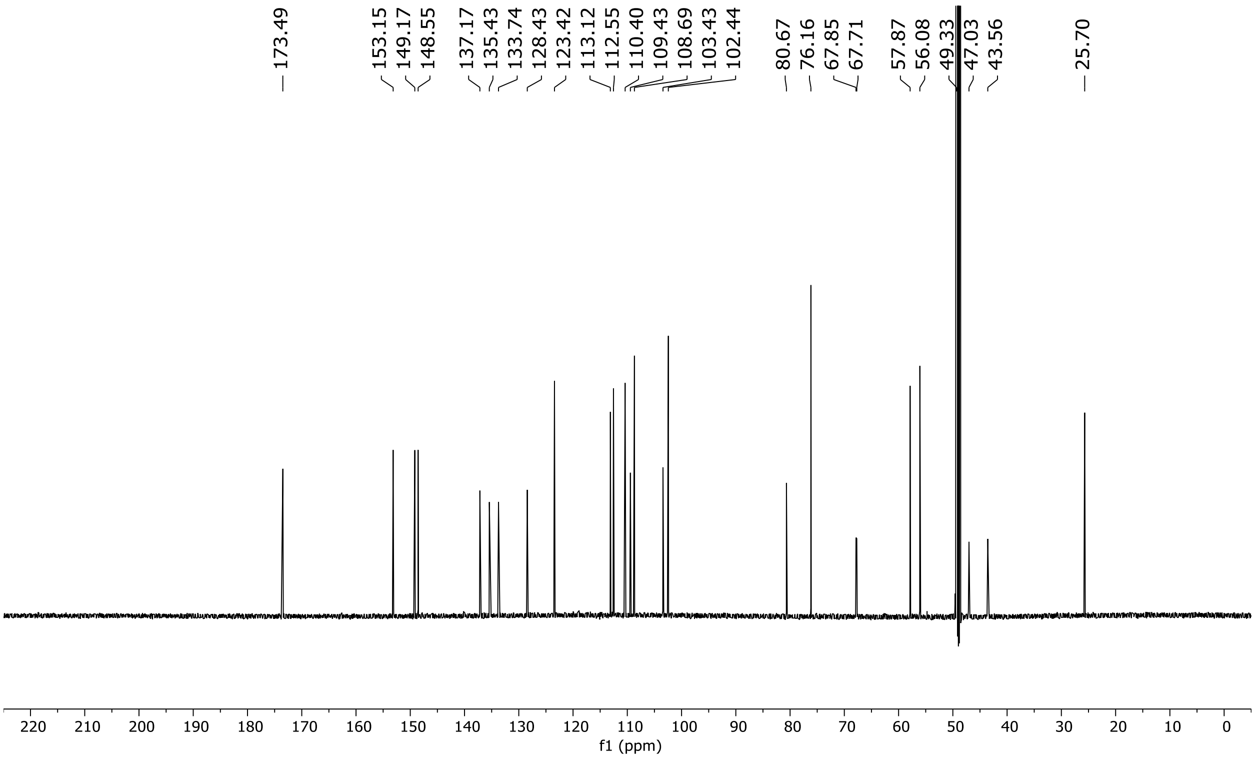

[1]H NMR (600 MHz, $CD_3OD$) for CJR-6A.

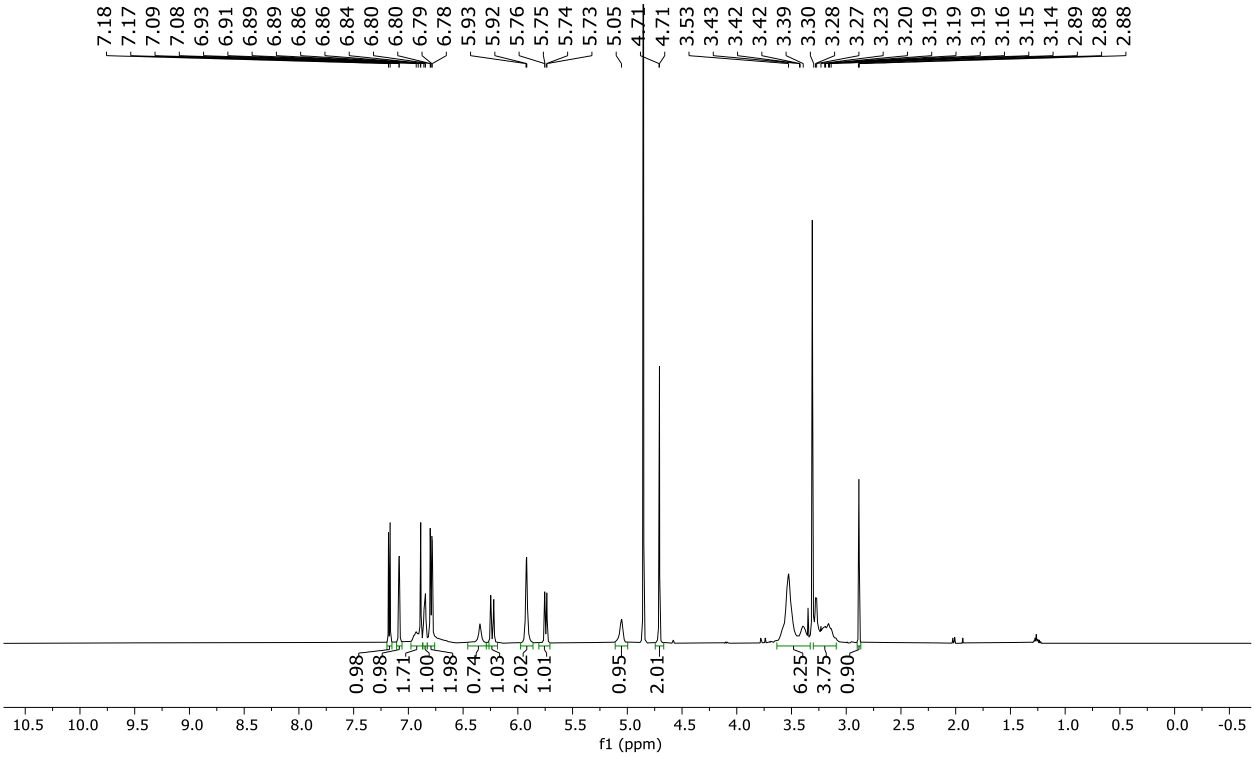

$^{13}C$ NMR (151 MHz, $CD_3OD$) for CJR-6A.

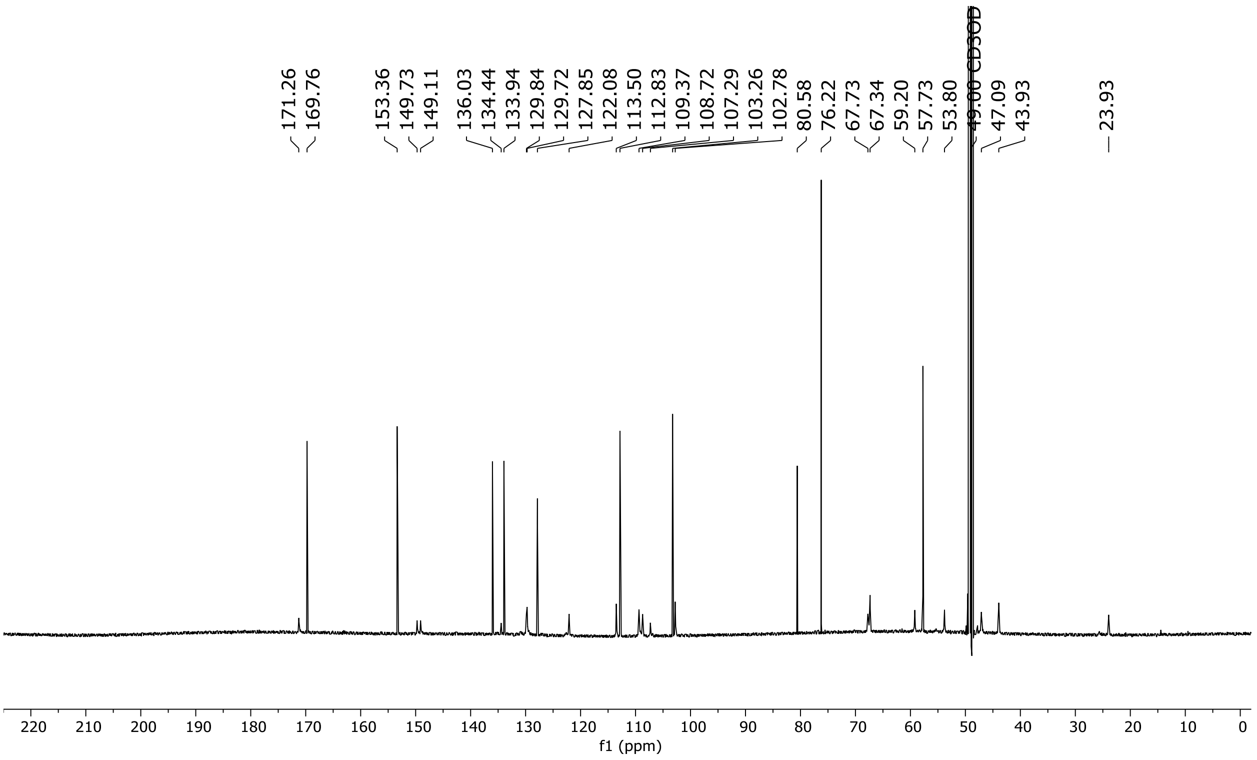

$^{1}$H NMR (600 MHz, $CD_3OD$) for ent-S15.1.

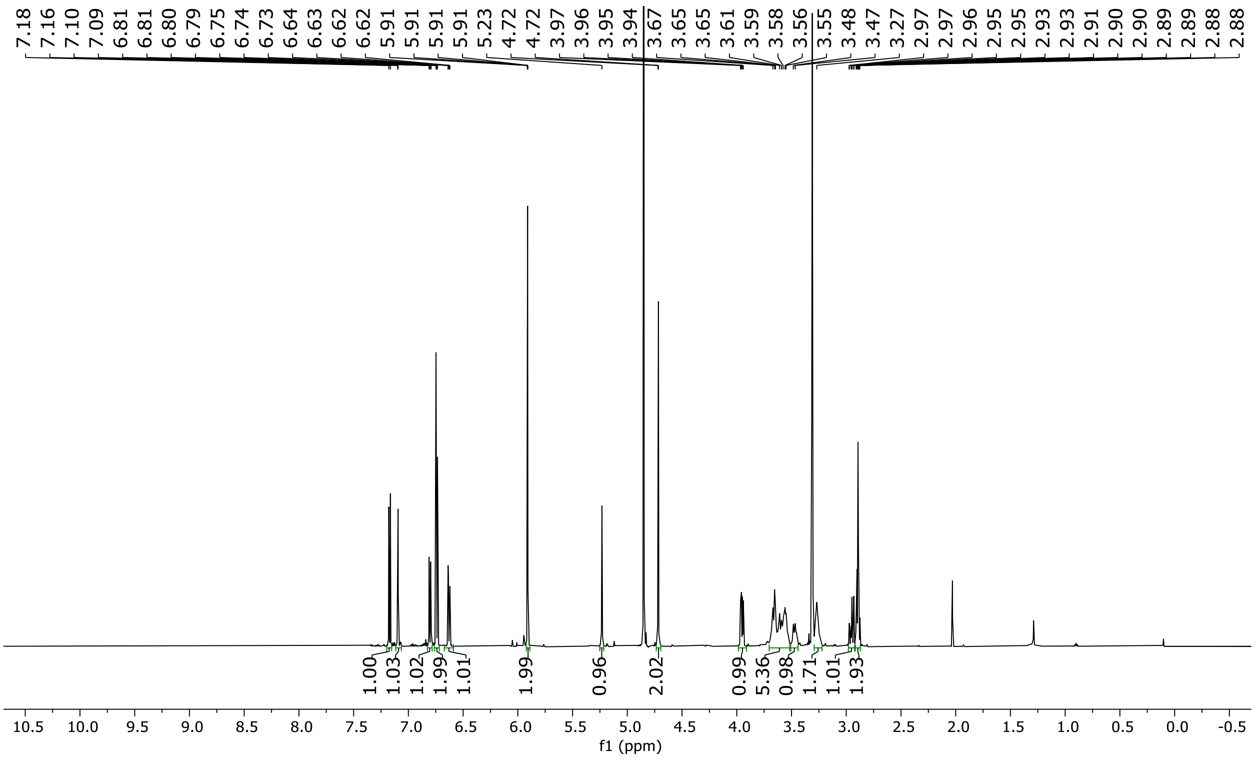

$^{13}C$ NMR (151 MHz, $CD_3OD$) for ent-S15.1.

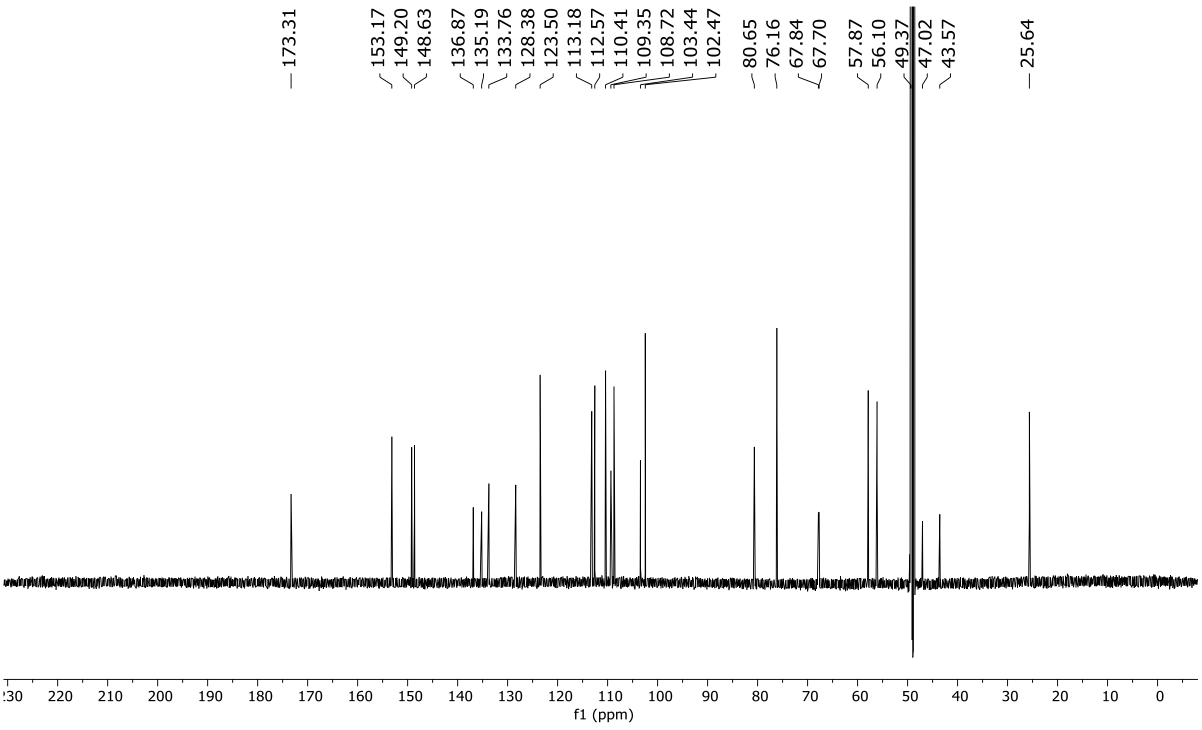

$^{1}H$ NMR (600 MHz, $CD_3OD$) for CJR-6B.

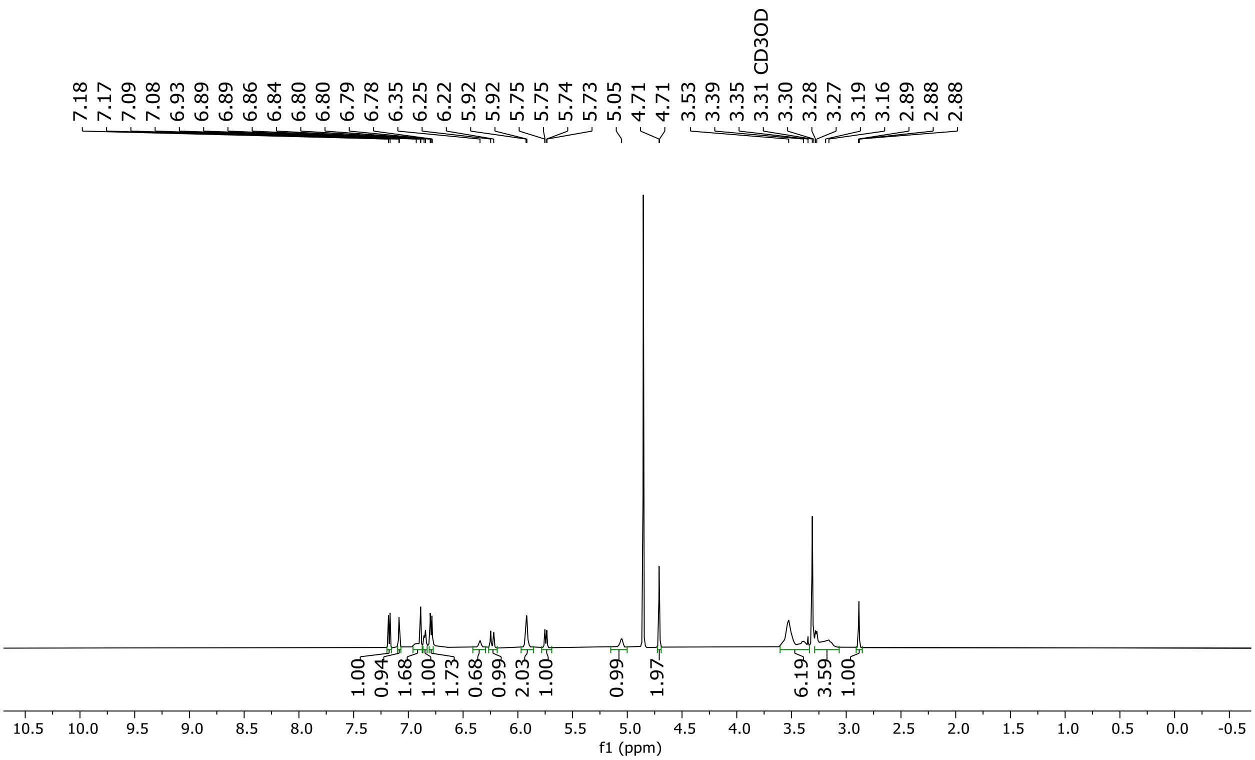

$^{13}C$ NMR (151 MHz, $CD_3OD$) for CJR-6B.

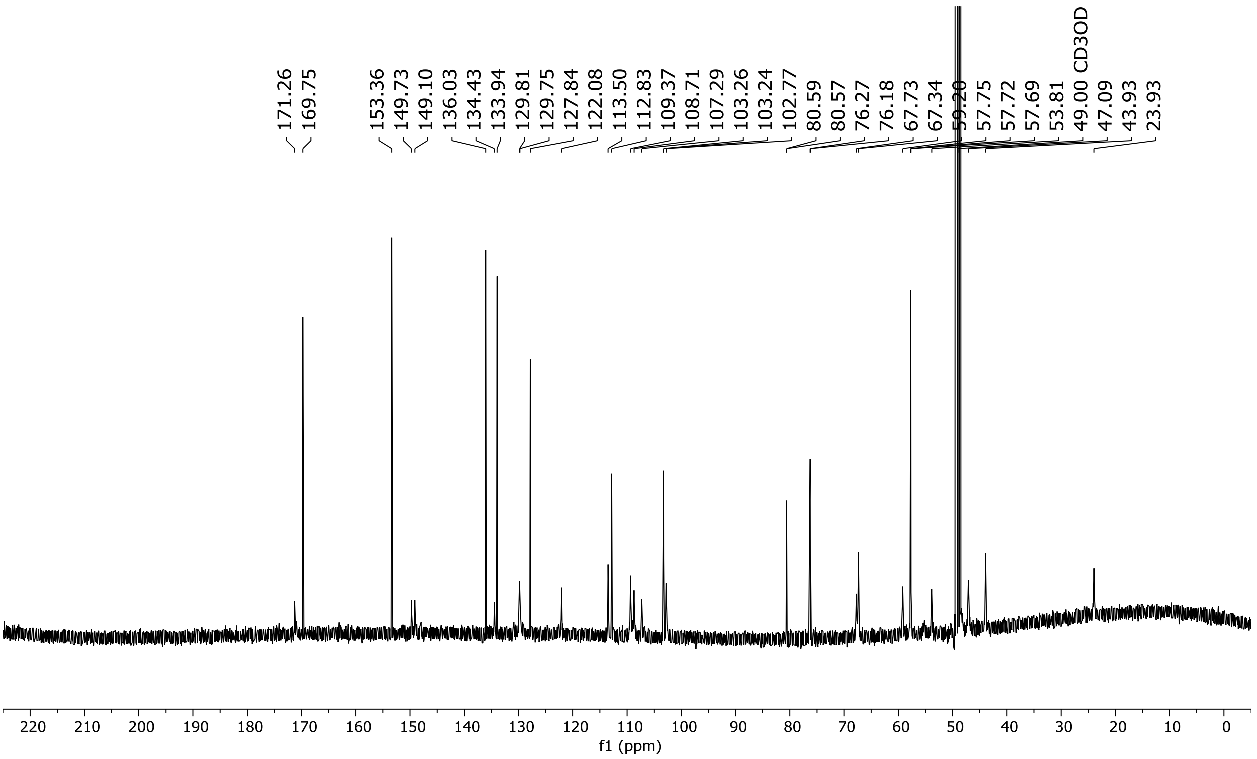

**Chiral-phase SFC data.**

Chiral-phase SFC chromatogram for mixture of CJR-6A (ent2) and CJR-6B (ent1).

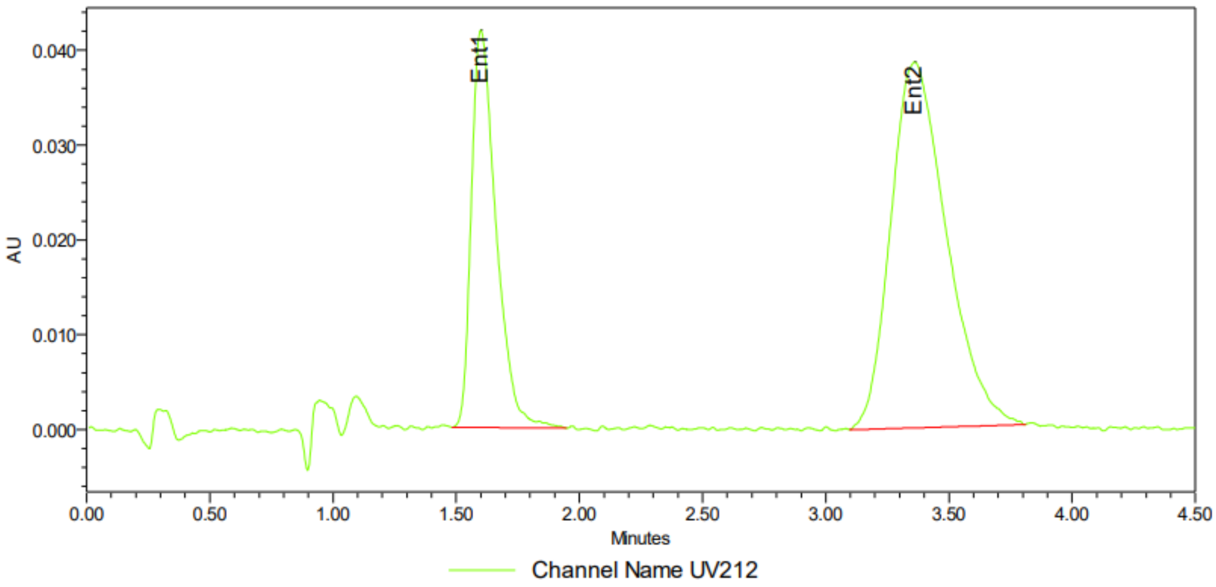


Peak Info

| | Channel Name | Name | RT | Area | Height (µV) | ent1 | ent2 | ee |
|---|---|---|---|---|---|---|---|---|
| 1 | UV212 | Ent1 | 1.60 | 291172 | 41914 | 32.40 | 67.60 | -35.20 |
| 2 | UV212 | Ent2 | 3.36 | 607473 | 38610 | 32.40 | 67.60 | -35.20 |

Chiral-phase SFC chromatogram for CJR-6A (ent2).

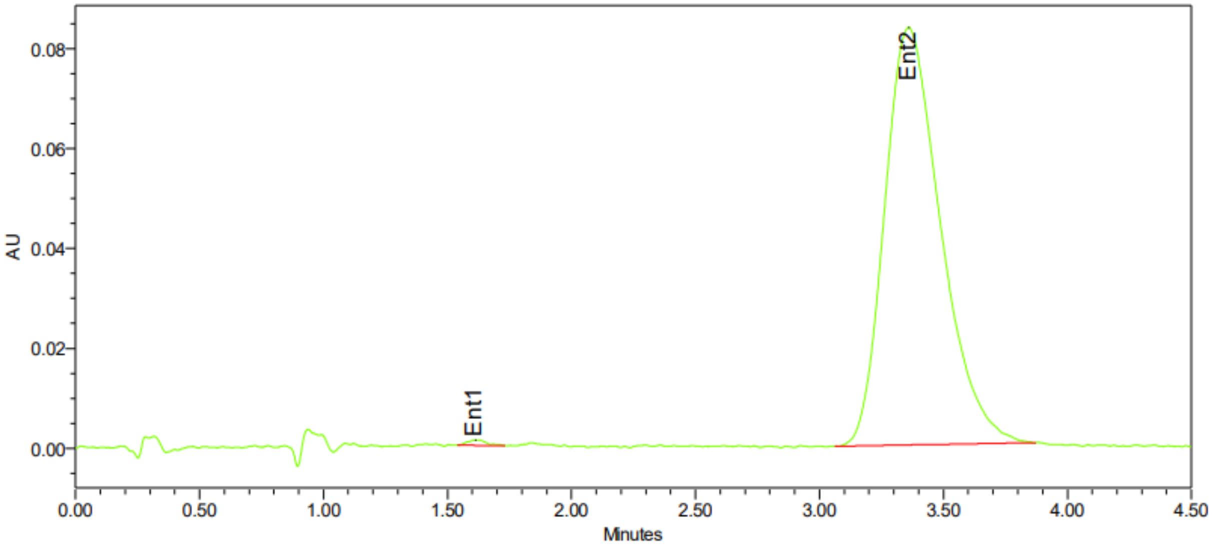


Peak Info

| | Channel Name | Name | RT | Area | Height (µV) | ent1 | ent2 | ee |
|---|---|---|---|---|---|---|---|---|
| 1 | UV212 | Ent1 | 1.61 | 5871 | 1061 | 0.44 | 99.56 | -99.11 |
| 2 | UV212 | Ent2 | 3.36 | 1314113 | 83649 | 0.44 | 99.56 | -99.11 |

Chiral-phase SFC chromatogram for CJR-6B (ent1).

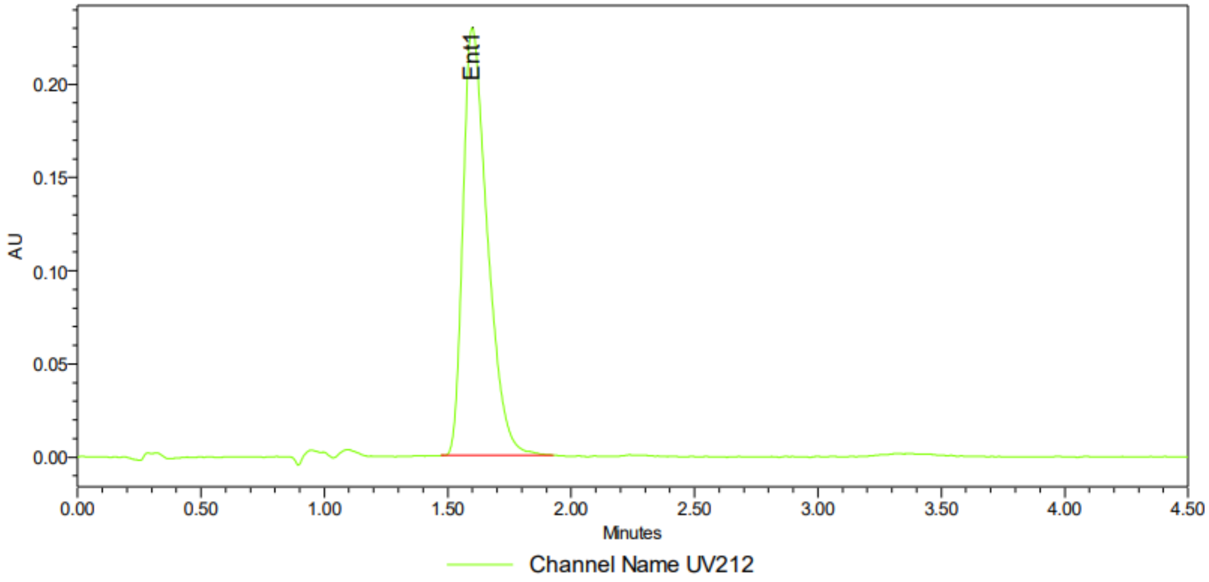


Peak Info

| | Channel Name | Name | RT | Area | Height (μV) | ent1 | ent2 | ee |
|---|---|---|---|---|---|---|---|---|
| 1 | UV212 | Ent1 | 1.60 | 1582133 | 229500 | 100.00 | | 100.00 |
| 2 | UV212 | Ent2 | 3.30 | | | 100.00 | | 100.00 |
